# Ecological genomics in the Northern krill uncovers loci for local adaptation across ocean basins

**DOI:** 10.1101/2023.04.15.537050

**Authors:** Per Unneberg, Mårten Larsson, Anna Olsson, Ola Wallerman, Anna Petri, Ignas Bunikis, Olga Vinnere Pettersson, Chiara Papetti, Ástþór Gíslason, Henrik Glenner, Joan E. Cartes, Leocadio Blanco-Bercial, Elena Eriksen, Bettina Meyer, Andreas Wallberg

## Abstract

Krill is a vital food source for many marine animals but also strongly impacted by climate change. Genetic adaptation could support populations, but remains uncharacterized. We assembled the 19 Gb Northern krill genome and compared genome-scale variation among 74 specimens from the colder Atlantic Ocean and warmer Mediterranean Sea. The genome is dominated by methylated transposable elements and contains many duplicated genes implied in molting and vision. Analysis of 760 million SNPs indicates extensive homogenizing gene-flow among populations. Nevertheless, we detect extreme divergence across hundreds of genes, governing ecophysiological functions like photoreception, circadian regulation, reproduction and thermal tolerance. Such standing variation may be essential for resilience in zooplankton, necessitating insight into adaptive variation to forecast their roles in future marine ecosystems and support ocean conservation.

**One-Sentence Summary:** Genome-scans of Northern krill link genes for photoreception, reproduction and thermal tolerance to ecological adaptation.

## Main Text

Climate change is affecting all life on Earth and forcing species to move or adapt (*1*). Ocean plankton are crucial to maintaining food webs and fisheries but face many challenges including increased temperatures and acidification (*2*–*4*). Many planktonic species are shifting toward higher latitudes (*4, 5*), and continued warming is expected to impact marine communities and ecosystem services (*6*). The long-term responses to these changes are unclear, but evolutionary adaptation may be important to sustain populations, particularly when physiological and geographical limits have otherwise been reached (*7*). This warrants the need to better understand adaptation in key zooplankton that strongly influence marine ecosystems. Krill (Euphausicea; 86 spp.), or euphausiids, are macrozooplankton crustaceans inhabiting all world oceans. Some species include trillions of individuals and are among the most abundant animals on Earth (*8*– *10*). As grazers of smaller plankton and food for fish and mammals, krill are critical links between primary production and higher trophic levels (*3*). However, polar krill of both hemispheres have declined in recent decades (*11, 12*), while boreal species such as the Northern krill *Meganyctiphanes norvegica* spread into new areas (*13*), impacting native biodiversity (*14, 15*).

The Northern krill is the largest and most abundant North Atlantic krill species, possibly structured into 3–4 basin-scale gene pools (*9, 16*). While many krill species are stenothermal and have narrow latitudinal ranges, it has unusually broad thermal tolerance and range (*17, 18*). It occurs across a 2–15°C temperature gradient (Fig. 1A) and breeds within 5–15°C (*9*), much wider thermal envelopes than for example the Antarctic krill *Euphausia superba* that is constrained within -2.0°C to +4.0°C and reproductively challenged already at +1.5°C (*19*).

**Fig. 1.**
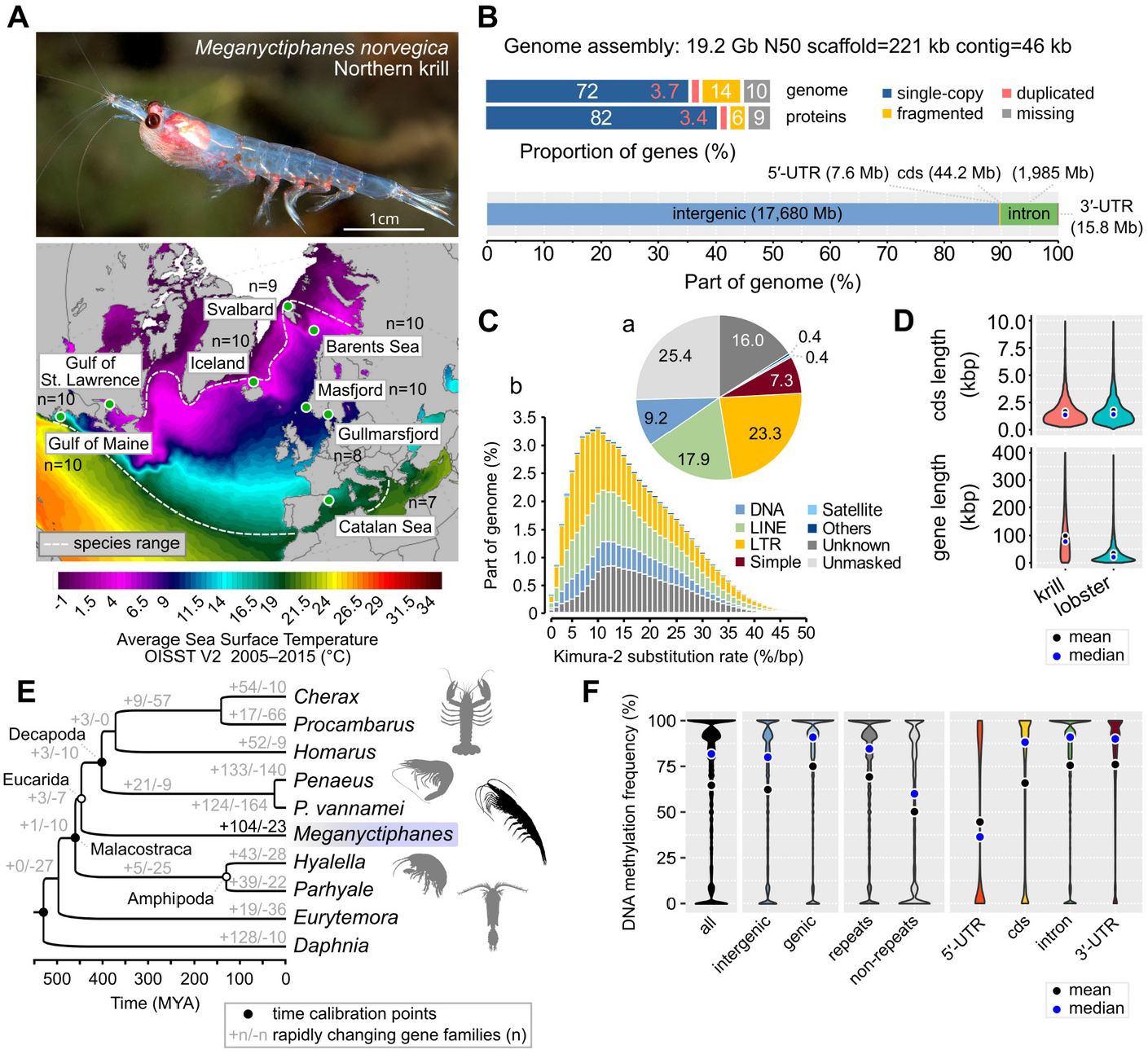
Sampling, genome assembly and genome analyses of the Northern krill *Meganyctiphanes norvegica*. (**A**) Photo: Adult specimen from the Norwegian Lysefjord (©Rudolf Svensen; approximate scale). Map: Atlantic and Mediterranean sample locations (n=sample-sizes). Sea Surface Temperatures from Climate Reanalyzer (*29*). (**B**) Genome assembly statistics in kilobases (kb), megabases (Mb) or gigabases (Gb). Top: completeness and duplications of BUSCOv5 genes (n=1,013) across the genome assembly and protein models. Bottom: sizes of genomic regions (UTR=untranslated exonic regions; cds=coding). (**C**) The repeat landscape. (a) Proportions of repeat-masked and unmasked bases (%). (b) The divergence landscape of interspersed repeats. (**D**) The coding and full sequence lengths of 7,150 single-copy orthologs between *M. norvegica* and the lobster *H. americanus*. (**E**) Time-calibrated species tree (inferred from 1,011 single-copy orthologs; 100% bootstrap support for all nodes) and gene family evolution. (**F**) The proportion of methylated cytosines across 75 M CpG-dinucleotides in the genome (>10× coverage).

Northern krill from different climates vary in metabolism, nutrition, maturation and timing of reproduction that track local seasonal cycles (*17, 20*), ranging from spawning in late winter–early spring in the Mediterranean Sea to summer in the Atlantic Ocean. These phenotypic variations could have genetic bases, making *M. norvegica* an attractive model for environmental adaptation.

Zooplankton generally have large populations with considerable genetic variation, suggesting they have high potential to adapt to changing environments through natural selection (*21*). Many also exhibit extensive larval dispersal and gene flow, which could either interfere with local adaptation or help introduce adaptive variants (*21*). Genomic evidence for mechanisms of adaptation is still critically missing for zooplankton (*22*). In particular, we lack insight into adaptation in euphausiids, due to their large and repetitive genomes ranging between 11–48 Gb 4–16× the human genome), which have hindered genetic analysis until now (*23*). The recently published Antarctic krill genome represented a major step forward but revealed extensive genetic homogeneity and limited adaptive variation in that circumpolar and panmictic species (*24*).

Here we use long-read technologies to assemble and characterize the huge genome of the Northern krill. We perform genome-scale population re-sequencing to map genetic variation and uncover the evolutionary history of this widespread species. Our ultimate aim is to identify genetic adaptations to different environments across its range. Insight into adaptive variation can help monitoring evolutionary processes including migration and adaptation, identify genetic diversity hotspots or offsets and support forecasting resilience of krill stocks under climate change (*25*).

## Results

### A highly repeated genome

The Northern krill is diploid and has 19 large metacentric chromosome-pairs that are homomorphic between sexes (*26*). We assembled and scaffolded the nuclear and mitochondrial genomes with Nanopore long-reads, Chromium linked-reads and RNA data from mostly a single specimen (tables S1–3; figs. S1–3; materials and methods). We tracked contiguity, completeness and accuracy by mapping coding transcripts (figs. S4–5). The finished genome assembly spanned 19.2 Gb (n=216,568 scaffolds/contigs) with ∼90% of BUSCO genes present and ∼0.5 base-level errors/kb (Fig. 1B, table S4) . The genome is GC-poor (%GC=29.9) and repeat-rich. Low-complexity sequences and simple repeats span 15% of the genome and AT-microsatellites alone cover 6% (table S3). Using a custom transposable element (TE) library, we find that 74% of the genome is repetitive (Fig. 1C; table S5). While we are unable to classify all TEs, retrotransposons (LINEs+LTRs) outweigh DNA-transposons 4:1, similarly to the American lobster or black tiger shrimp (*27, 28*). This repeatome is different from the recently characterized genome of the Antarctic krill, which is dominated by DNA-transposons (*24*). Furthermore, TE divergence in the Northern krill is unimodal, without the multiple bursts of proliferation seen in the Antarctic krill. These observations hint at divergent genome composition and lineage-specific evolution of the huge genomes of euphausiids.

### Expansion of cuticular and opsin gene families

We used RNA and comparative data to annotate 25,301 protein-coding genes along with 2,283 TEs (mostly expressed retrotransposons), and also detected another 14,643 potential yet unannotated genes or TEs (tables S6–8; fig. S6). Gene bodies (introns+exons) span 50,276 bp on average and occupy 10% of the genome, while coding sequence covers only 0.22% (Fig. 1B).

Orthologous gene bodies between the Northern krill and crustaceans with smaller genomes (table S9) are 3–8× longer in krill, but have similar amounts of coding sequence (Fig. 1D; fig. S7, table S10). Compared to the Antarctic krill (*24*), genes are ∼2.5× longer in the Northern krill, suggesting proliferation of retrotransposable elements has produced long and highly repeated introns (fig. S6B). We estimated high synonymous divergence (*dS*=0.46) between the two species (table S10). Using a decapod molecular clock (*30*), this divergence suggests they split from a common ancestor ∼130 MYA, underscoring separate evolution over long time-scales.

We built a crustacean species tree and analyzed gene family evolution. We found 104 rapidly expanding gene families in the Northern krill (p<0.05; Fig. 1E; fig. S8A–B; table S11), including those related to chitin, cuticular metabolism, regulation of the molting cycle, which are important processes for growth and reproduction in crustaceans. This is notable as renewal of the exoskeleton is unusually frequent and plastic in euphausiids (*9, 31*), and similar expansions were independently detected in the Antarctic krill genome (*24*). Moreover, we detected expansions of the opsin gene repertoire, which encodes the light-sensitive receptors in ommatidia. Fourteen opsins have previously been identified from RNA in the Antarctic krill *E. superba* (*32*), which are thought to enable vision under the divergent light conditions experienced throughout its life cycle and vertical migrations (*33*), while 16 opsins have recently been inferred from *M. norvegica* RNA (*34*). We queried our *M. norvegica* gene-set and the KrillDB^2^ *E. superba* RNA database against the curated crustacean opsin dataset in ref 35. We detected 19 genes in the former species and 15 putative genes in the latter (fig. S9–10), including new visual middle wavelength-sensitive (MWS) opsins and non-visual arthropsins. All *E. superba* opsins have homologs in the *M. norvegica* genome and all but one previously identified *M. norvegica* transcripts can now be anchored unambiguously 1:1 to our gene models (fig. S10). Our findings expand the known opsins in both species and suggest that opsin and molting-gene duplications could be common to all euphausiids.

Ancient whole-genome duplication (WGD) may explain the evolutionary origins of large genomes but is not commonly reported for crustaceans. To test for WGD, we interrogated divergences between gene-paralogs and searched for Hox-gene duplications, but found no supporting evidence (Supplementary text). Instead, the huge Northern krill genome has likely evolved through TE proliferation and numerous small-scale duplications.

### An active DNA methylation system

Epigenetic regulation of the genome may contribute to genome evolution, function and plastic responses to environmental change (*35*). For example, CpG-methylation can silence the expression of harmful transposable elements (TEs), paradoxically allowing them to persist and contribute to genome expansion (*36*). DNA methylation (DNAm) of both TEs and protein-coding genes is ancestral in Arthropoda, but has frequently been lost (*37*). Using DNA from muscle, we characterize, for the first time, the DNAm toolkit and genomic patterns in a euphausiid. The genome has low CpG content (CpG_O/E_=0.53), indicating DNAm. We find all genes encoding the canonical methyltransferases and genes responsible for repair or demethylation (*Dnmt1–3, AlkB2, TET2*; fig. S12–16; table S12), hallmarks of functional DNAm (*37*). We scanned the Nanopore-reads for signals of CpG-methylation at 75 million CpG-sites. Overall, DNAm rates are higher in genes than intergenic regions (75% vs. 62% of reads being methylated; Fig. 1F; fig. S17) and positively associated with splice isoform variation (n=2.5±0.05 isoforms/gene with >95% methylation rates vs. 1.8±0.03 isoforms/gene with <5% methylation rates; fig. S17E). In contrast, we observed a lack of methylation in the mitochondrial chromosome (4%). DNAm rates are higher across repeats vs. non-repeated DNA (69% vs. 50%), and appear to target young retrotransposons with similar LTRs, that may have been recently active (S17F). Gene-body methylation is similar to observations in marbled crayfish (*38*), while repeat-oriented methylation reminds of the myriapod *Strigamia maritima* (*37*). The krill methylome thus spans both genes and repeats, suggesting dual roles in gene regulation and silencing TEs.

### Genome-scale variation is shaped by linked selection and pervasive gene flow

To uncover patterns of genetic variation, we collected 74 Northern krill specimens from eight geographical regions separated by up to 5,800 km, covering a range of environmental conditions (Fig 1A; Supplementary text). We re-sequenced whole genomes to ∼3×/specimen (20–30×/population), mapped reads and called 760 million quality-filtered SNPs across accessible sites determined with similar filters (8.4 of 19.2 Gb; fig. S18). We estimate genome-wide nucleotide diversity (π; the average differences between pairs of sequences) to 1.31% perbase, intermediate among arthropods and low compared to marine broadcast spawners like oysters and sea-squirts (*39*). The population mutation rate (*θ*_w_) is 1.62%/bp. Assuming mutation-drift equilibrium and a mutation rate from snapping shrimp (*30*), we estimate the long-term effective population size (*N*_E_) to be 1.53 million, far below the expected census population size of trillions (*9*). However, Tajima’s D is negative (-0.53), indicating excess of low-frequency variants compared to expectation under equilibrium, consistent with population expansion or selection shaping genetic variation. In accordance, we observe ∼30% reduction of variation over genes (π=1.15%), and more so at coding and non-synonymous sites (Fig. 2Aa–b). This effect extends up to 50–100 kb around genes (Fig. 2Ac), suggesting widespread impact of linked selection. Using heterozygous genotypes in the reference specimen (∼5,000 scaffolds >500 kb), we applied the Sequentially Markovian Coalescent to model demographic history (*40, 41*). The results indicate populations expanded half way through the last glacial period (*42*) (Fig. 2B). Levels of variation are similar among populations (fig. S19A). Population structure is limited but recapitulates geography (the fixation index *F*_ST_ is ≈0.06 on average; Fig. 2C) and previously detected mitochondrial gene pools (*9*). Genetic distances (d_XY_) only increase marginally with geographic distance (fig. S19B–D; S20). The Mediterranean Sea sample is the most divergent, but average d_XY_ is only 1.04× higher compared to distances among Atlantic populations (1.71% vs. 1.64%). The majority of non-singleton variants are polymorphic in most populations (fig. S21), indicating extensive gene flow among stocks.

**Fig. 2.**
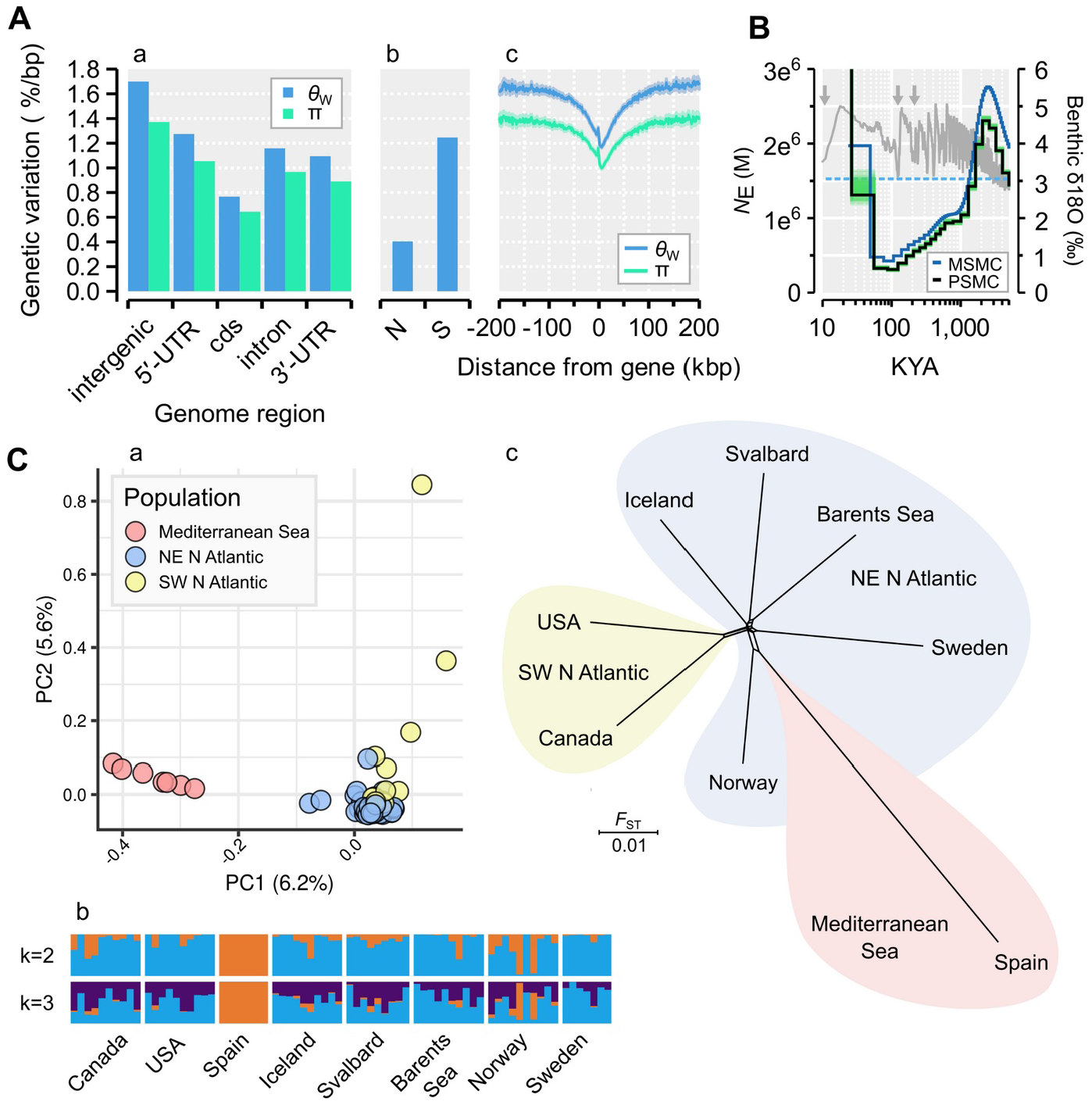
Genome-scale patterns of genetic variation, demographic history and population structure. **(A)** The mean levels of genetic variation across: (a) the genome; (b) at non-synonymous (N) and synonymous sites (S); or (c) across 1 kb non-overlapping windows upstream or downstream of putative protein-coding gene bodies. Shaded regions are 95% confidence intervals (200 bootstrap replicates). (**B**) Historical fluctuations in effective population size (*N*_E_) inferred using PSMC and MSMC (y1-axis; green=100 PSMC bootstrap replicates; KYA=thousands of years ago). The dashed line is long-term *N*_E_. The LR04 benthic δ^18^O isotope (gray line, y2-axis) indicates ice volume and sea temperature (arrows=last three interglacials). (**C**) Sample and population interrelationships. (a) Main clustering of genetic variation in a principal component analysis. (b) Admixture analyses indicating levels of shared ancestry. (c) NeighborNet network of pairwise *F*_ST_-distances.

### Signatures of ancient adaptive divergence across hundreds of genes

To reveal genomic signatures of adaptation, we partitioned the dataset into two major contrasts:

i. Atlantic vs. Mediterranean samples (at/me); and ii) North-Eastern vs. South-Western North Atlantic samples (ea/we; Fig. 3A). For each contrast, we computed pairwise divergence in allele frequencies. While most variation segregates at low differences (*F*_ST(at/me)_=0.056; *F*_ST(ea/we)_=0.017; Fig. 3B; fig. S20A), we also detect about 8× as many highly divergent variants compared to expectations from simulations of neutral drift (figs. S22B–C). Divergent regions (*F*_ST_>0.4) span <1% of the genome and are about 2× enriched for gene sequences and for extended haplotypes, compared to undifferentiated regions (*F*_ST_<0.2; Fig 3B), consistent with gene-centered signatures of selective sweeps (*43*). We compared *F*_ST_ between genes and similarly sized 50 kb flanking regions 50–100 kb away from genes, which may more often evolve neutrally. At extreme levels of divergence (the top 0.1% most divergent flanking regions), genes outweigh flanking regions by 7× (at/me) or 2× (ea/we), respectively, consistent with natural selection driving divergence across many genes (Fig. 3C; table S13; fig. S23).

**Fig. 3.**
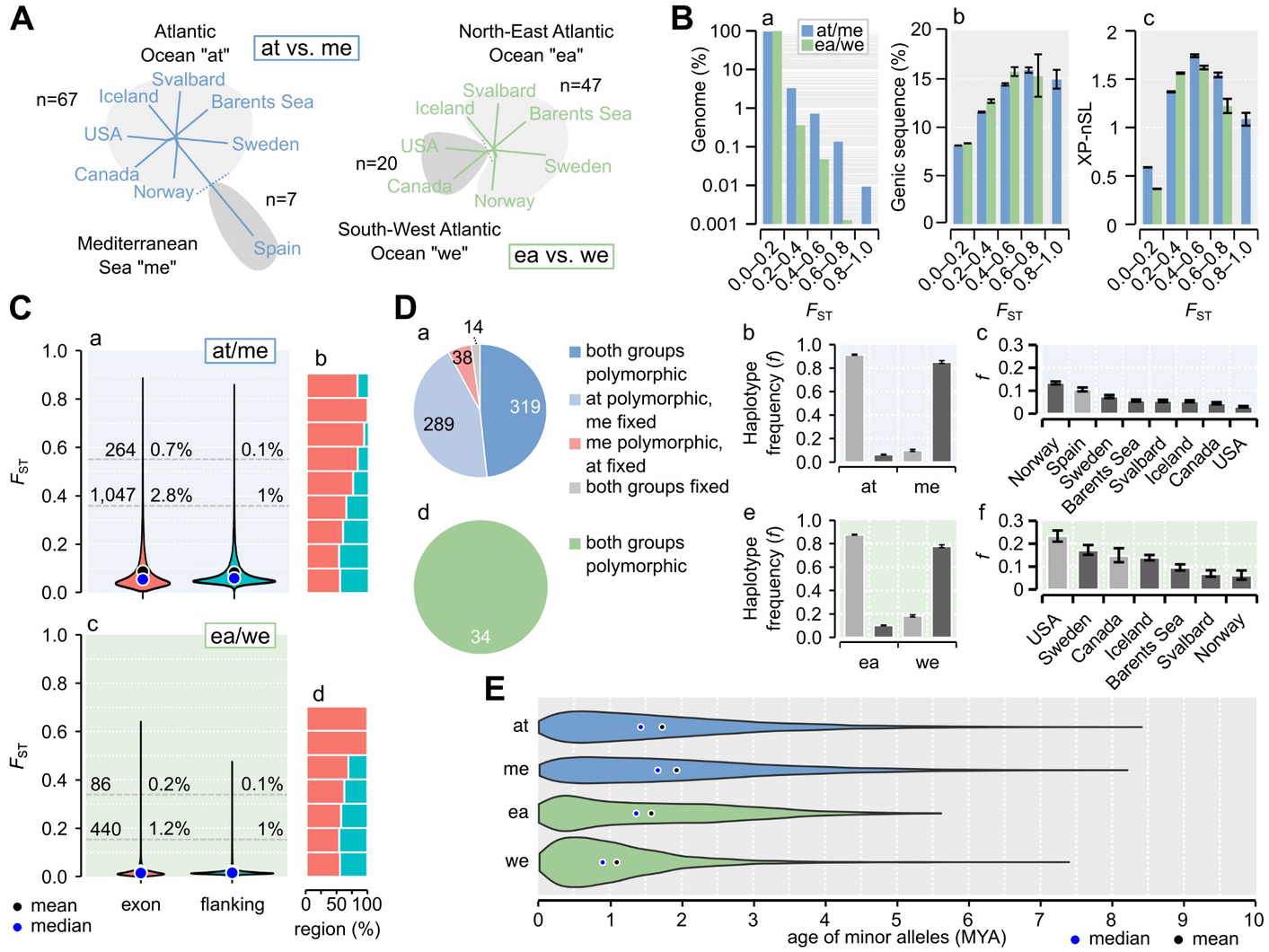
Genetic divergence among Northern krill sampled across the Atlantic Ocean and Mediterranean Sea. (**A**) The two contrasts used to measure divergence. (**B**) Features of genomic regions at increasing divergence (mean *F*_ST_ across 1 kb windows). (a–b) The proportion of the genome or genic sequence (cds+intron+UTRs). (c) The absolute extended haplotype statistic |XP-nSL| scores (95% confidence intervals from 2,000 bootstrap replicates). (**C**) (a–c) Atlantic-Mediterranean contrast: (a) The distribution of per-gene *F*_ST_-values computed across the exons of genes (cds+UTRs; n=36,997) vs. flanking intergenic regions (n=31,101). Gray lines are 1% and 0.1% percentiles of flanking *F*_ST_. Numbers and proportions of genes are indicated at each percentile. (b) The proportion of genes vs. flanking regions at each *F*_ST_-level. (c–d) Eastern-Western contrast: statistics as in a–b (n=36,397 vs. n=31,103). (**D**) Haplotype distribution for divergent genes (exon-wide *F*_ST_>0.4). (a–c) Atlantic-Mediterranean contrast. (a) Numbers of genes with shared or private haplotypes (n=660). Mean frequencies of common and rare haplotypes in each group (b) and population (c) (95% confidence intervals as in (B)). (d–f) as in (a–c) but for 34 genes in the Eastern-Western contrast. (**E**) Age distributions of minor alleles across (at/me: ∼22–25 K SNPs in 660 genes; we/ea: ∼2.5 K in 34 genes).

We analyzed the geographic distribution of putatively adaptive variation by defining gene-level haplotypes (having at least four diagnostic SNPs with *F*_ST_>0.5). At many divergent genes (exon-wide *F*_ST_>0.4), both haplotypes are often present in both groups (n_at/me_=319/660; n_ea/we_=34/34), indicating widespread standing variation (Fig. 3D). Southern or Scandinavian populations are more polymorphic than Barents Sea and Svalbard populations (Fig. 3D), which could reflect genetic drift or ongoing selection at the margin of the Arctic species range (*44*). We estimated the ages of minor alleles on the divergent haplotypes to learn for how long haplotypes may have been segregating in the species. We first estimated a genome-average recombination rate (r=0.32cM/Mb, n=652 scaffolds) and then applied the Genealogical Estimation of Variant Age (GEVA) (*45*). This coalescent method infers the time to the most recent common ancestor using mutation and recombination rates, without requiring *a priori* assumptions about demographic history. Most variation originated over 1 MYA, predating multiple glacial cycles, and adaptive variation segregating between Atlantic and Mediterranean populations may predate that segregating in the Atlantic Ocean (Fig. 3E).

### Candidate genes for ecological adaptation associated with ecophysiological functions

For each contrast, we ranked all genes by exon-wide *F*_ST_. The top gene between Atlantic and Mediterranean krill is *nose resistant to fluoxetine protein 6* (*nrf-6*) (table S13), encoding a membrane protein facilitating lipid transport. It is located within a high-divergence region and the Mediterranean samples are fixed for an *nrf-6* haplotype with low frequency among Atlantic samples (6%). Exon-wide *F*_ST_ is >0.8 and high XP-nSL scores indicate loss of variation consistent with a selective sweep among Mediterranean samples (Fig 4Aa–b). We also detect accelerated protein evolution on the Mediterranean haplotype (*dN/dS*: 0.54 vs. 0.31; fig. S24A– B), consistent with positive selection. We predicted the protein topology and detected 1.7× enrichment of missense variants in its extracellular part compared to synonymous variants (9/16 vs. 5/15; Fig 4Ac; fig. S24C-D; table S14), which may alter its function. This is noteworthy as *nrf-6* is important for yolk transport into eggs in worm (*46*) and ovary development and oogenesis in fly (*47*), and overexpressed in the ovaries of sexually precocious crabs (*48*). In the Northern krill, Mediterranean stocks depend on a short spring phytoplankton bloom to accumulate lipid stores and trigger vitellogenesis and reproduction in early spring (*9, 49*). The Mediterranean *nrf-6* variant could contribute to advanced reproductive timing, making it a strong candidate for local adaptation.

**Fig. 4.**
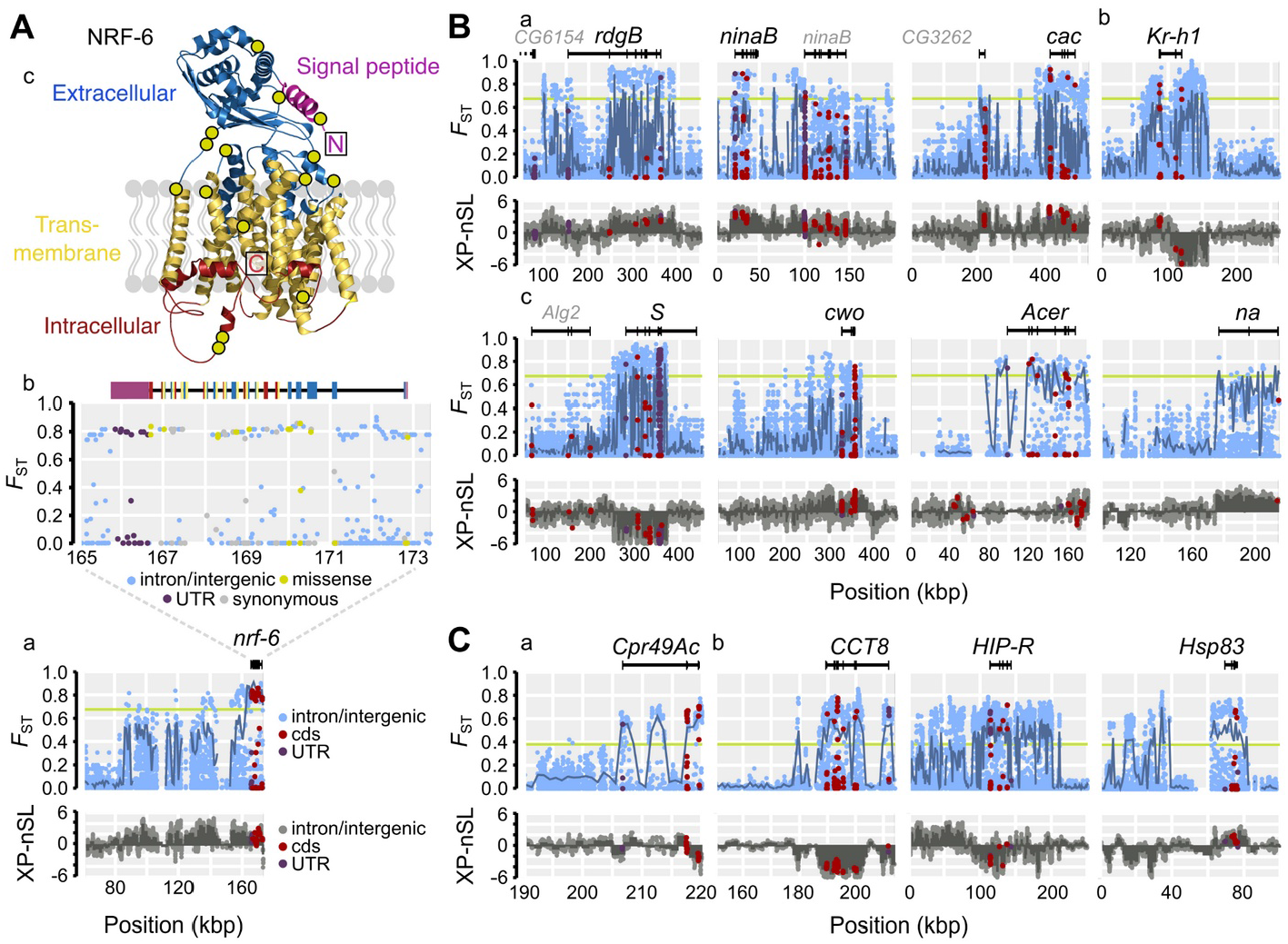
Adaptive divergence and candidates for local adaptation. (**A**) High divergence between Atlantic and Mediterranean samples across the gene *nrf-6*. (a) Top: per-SNP *F*_ST_ along the locus (green line=0.1% percentile for SNPs; dark-blue line=*F*_ST_ for 1 kb windows; black=gene model). Bottom: per-SNP XP-nSL (dark-gray bars=mean window-based XP-nSL). Positive values imply a selective sweep in the Mediterranean sample, negative values mark the Atlantic Ocean sample. (b) Magnified view of *nrf-6*. Exons colored by UTR (purple) or NRF-6 protein topology. (c) Modeled protein structure and topology (including signaling peptide; boxes=N/C terminals; green circles=non-synonymous/missense variants). (**B**)Examples of highly differentiated genes between Atlantic and Mediterranean samples. (a) Photoreception: *retinal degeneration B* (*rdgB*), *neither inactivation nor afterpotential B* (*ninaB*) and *cacophony* (*cac*). (b) Heterochronic development: *Kruppel homolog 1* (*Kr-h1*). (c) Circadian regulation: *Star* (*S*), *clockwork orange* (*cwo*), *Angiotensin-converting enzyme-related* (*Acer*) and *narrow* (*na*). Statistics as in (A). (**C**) (a) The top gene in the Atlantic comparison was *Cuticular protein 49Ac* (*Cpr49Ac*) (b) Three chaperon genes implied thermal tolerance: *Chaperonin containing TCP1 subunit 8* (*CCT8*), *Hsc/Hsp70-interacting protein related* (*HIP-R*) and *Heat shock protein 83* (*Hsp83*).

We next queried the ranked lists for shared gene ontologies using used fly homologs. Atlantic-Mediterranean divergence is enriched for genes involved in oogenesis, muscle function, phototransduction and eye development, regulation of circadian rhythm and heterochronic development (fig. S25A; table S15). These include homologs for *ninaB* that synthesize visual pigment (*50*), *S* that regulate sleep/wake cycles in animals and the transcription factor *Kr-h1* that acts in the juvenile hormone signaling pathway to govern vitellogenesis and reproduction in arthropods (*51*) (Fig 4B). Overall, 30 genes in the 0.1%-percentile (*F*_ST_=0.60–0.72; n=264) are associated with vision-related ontologies, 1.5× more than expected by chance (p=0.033), and 112 genes in the 1%-percentile (*F*_ST_=0.36–0.72; n=1,024), including four MWS opsins (table S15).

Photoreception candidates often involve paralogs (fig. S26) and belong to rapidly expanded gene families, which are otherwise underrepresented in the 1%-percentile (2.98× vs. 0.6×; table S15).

Between the two Atlantic basins, the most divergent gene encodes a larval cuticular protein (homolog of *Cpr49Ac*; Fig 4C), while this contrast is enriched for signaling ontologies (fig. S25A–B). Among the top 20 genes we identify three heat-shock chaperones/co-chaperones, including *CCT8* (Fig 4C), that fold proteins and promote proteostasis under thermal stress. Heat-shock proteins contribute to protective thermal tolerance in krill and many other species (*52*). Chaperonins (*CCT-n*) influence cold shock response in crustaceans and other eukaryotes (*53*), and *CCT8* evolution is implied in freeze-tolerance in Amur sleeper fish (*54*). North-Eastern krill have a sweep-like signature of extended *CCT8* haplotypes, suggesting natural selection for cold tolerance in these stocks.

### Conclusions and implications

We here provide novel insight into the evolutionary history of the Northern krill. Our genome assembly reveals a highly methylated repeatome and many expanded gene families. Paralogous genes may have arisen from ectopic recombination between non-homologous repeated loci, which is more likely to occur when genomes accumulate transposable elements (*55*). The krill genome appears to continuously have evolved new genes, including those involved in molting and vision.

Molting is a crucial process in krill, being interlinked with growth and reproduction, and controlled by environmental cues including light (*9, 56*). Some of the top gene candidates for local adaptation belong to expanded gene families, including two *ninaB* paralogs and four *ninaE* paralogs that synthesize visual pigment or encode MWS opsins. We detect elevated divergence in fifteen genes encoding cuticular proteins, several belonging to expanded families. Expanded cuticular and opsin gene families and functionally diverged paralogs may have enabled development and photoreception under diverse conditions in ancestral krill and provide substrate for adaptation also today (*57*).

We have detected many functionally related genes with small-to-moderate shifts in allele frequencies, consistent with signatures of polygenic adaptation (*58*). Evolve-and-resequence experiments in copepods have similarly found polygenic signals of adaptation to temperature and acidity in laboratory conditions (*59*). Genetic adaptation in zooplankton could commonly involve numerous loci, warranting genome-scale assays to map adaptive variation. Many of our candidate genes have roles in photoreception, circadian rhythm, and oogenesis, ecophysiological functions also implied in adaptation in other widespread pelagic species, such as the Atlantic herring (*60*). The variants may help krill respond to light, temperature and resources in different environments.

Photoreception varies among *M. norvegica* populations: krill from turbid waters around the Gulf of Maine are more light sensitive than those from clearer waters (*61*). Water clarity and light penetration influences behavior in *M. norvegica*. To avoid predators, stocks prefer deeper depths in more clear Norwegian fjords (*62*), while the deepest daytime depths (400–800 m) are known from the oligotrophic Mediterranean Sea (*9*). Moreover, light sensitivities in the North Pacific krill *E. pacifica* are tuned to local conditions. Individuals inhabiting shallow green water in the Saanich Inlet are more sensitive to green light compared to those from the deeper blue water of the San Diego Trough (*63*). At least thirty genes involved in eye function diverge strongly between Atlantic and Mediterranean krill (including 4 *ninaE*/MWS paralogs), and another six genes segregate across the Atlantic Ocean, suggesting heritable variation could contribute to these phenotypes. Eye traits are generally fast-evolving among krill species and associated with ecological niche (*64*). Our candidates could help reconstruct the genomic architecture of vision and behavior in krill.

Seasonal and daily cycles of ambient light are central to zooplankton and influence diapause, vertical migrations and entrainment of endogenous circadian clocks that control the daily rhythm of physiological processes (*65*). *E. superba* and *Thysanoessa inermis* krill have endogenous circadian rhythms (ECR) that oscillate faster than 24 h in the absence of light, or respond to minute irradiance. This may reflect adaptation to extreme photoperiodic variability at high-latitudes where the sun is either below or above the horizon for extended periods (*66, 67*), although the genetic mechanisms of these adaptations are unknown. *M. norvegica* also shows ECR and expresses a full set of circadian clock genes (*68*). We find that genes likely involved in regulating its circadian clock diverge across its range, including homologs of *narrow abdomen, glass* and *clockwork orange*. Functional assessments of the variants could illuminate how biological clocks are set in different environments.

Marine ecosystems are changing at unprecedented rates, causing redistribution of organisms and impacting food webs (*5, 15*). The Antarctic krill is already declining, which could severely impact the Antarctic ecosystem (*11, 14*). The Northern krill could be declining around Iceland (*69*), but increasing in the Barents Sea (*13*). Where will krill thrive in the future? We found many variants that could help krill adapt to new or changing environments, many of which are widely distributed, old and possibly maintained by long-term balancing selection under slowly fluctuating conditions. These adaptive variants could be important for coping with rapid climate change, and Scandinavian stocks in particular may serve as sources of genetic diversity typical to southern populations. The next frontier for the Northern krill is the Arctic Ocean (*13, 15*).

Standing variants supporting physiological processes under darker or colder conditions may help establish it there. The most divergent population is that of the Mediterranean Sea, which is close to its southern limit. This population appears to lack variation at many adaptive loci, which might limit its evolutionary potential, although our analysis is limited by a small sample size.

With the exception of the Antarctic krill (*8*), long-term monitoring of krill abundance is typically performed or reported in aggregate (*12*), obscuring how individual species fare under climate change. Our results suggest that krill may commonly be genetically fine-tuned to their environments, while previous research underscore limited thermal tolerances in many species (*18, 19*). Genetic adaptation could be the major process determining whether krill will persist or perish. The many candidate genes reported here can be used as biomarkers to diagnose and monitor change of adaptive variation also in other species and climates, in order to better forecast their distributions and estimate risks of the great many species that depend on them.

## Acknowledgments

We thank Stéphane Plourde, Geneviève Perrin, Jon Rønning, Monica B. Martinussen and Katharina Michael for providing samples, the staff at Kristineberg Center for Marine Research and Innovation, Zhen Li for constructive discussion and Jessica Heinze, Sarah Demirkale and Ylva Jondelius for lab assistance. The computations were enabled project SNIC 2022/5-472 provided by the National Academic Infrastructure for Supercomputing in Sweden (NAISS) and the Swedish National Infrastructure for Computing (SNIC) at UPPMAX and the PDC Center for High Performance Computing partially funded by the Swedish Research Council through grant agreements no. 2022-06725 and no. 2018-05973.

## Author contributions

Conceptualization: AW

Data curation: AW PU IB

Formal analysis: AW PU ML IB

Funding acquisition: AW

Investigation: AW AO AP OW

Methodology: AW PU AP

Project administration: AW OVP

Resources: AW OVP EE AG HG JC LBB

Software: AW PU

Supervision: AW

Visualization: AW PU ML

Writing – original draft: AW PU

Writing – review & editing: AW PU BM LBB EE AG HG JC CP

## Data and materials availability

The sequence data underlying this article are available in the European Nucleotide Archive at https://www.ebi.ac.uk/ena/browser/home, and can be accessed with project accession PRJXXXXXXX. The genome assembly, annotated SNPs and orthology datasets are available in the SciLifeLab Data Repository at https://doi.org/10.17044/scilifelab.c.XXXXXXX. Public code is available at https://github.com/NBISweden/genecovr and https://github.com/andreaswallberg/Ecological-Genomics-Northern-Krill. Reference specimen tissue is deposited in the LIB Biobank at Museum Koenig Bonn under accession ZFMK-TIS-XXXXX.

## Supplementary Materials

Materials and Methods

Supplementary Text

Figs. S1 to S26

Tables S1 to S15

References (*##*–*##*)

