## Supplementary materials for "Ecological genomics in the Northern krill uncovers loci for local adaptation across ocean basins"

**This PDF file includes:**

Materials and Methods
Supplementary Text
Figs. S1 to S26
Tables S1 to S15

**Table of Contents**

**Materials and Methods** 5
**Biological materials** 5
Processing of samples 5
**Overview of library preparation & sequencing** 5
High-molecular weight DNA extraction for long-read and linked-read libraries 6
PromethION long-read sequencing 6
10X linked-read sequencing 7
RNA extraction from multiple tissues for long-read and short-read RNA libraries 7
Short-read RNA sequencing 7
Long-read cDNA sequencing 7
Salt-isopropanol DNA extraction of multiple specimens for population-scale whole-genome re-sequencing 8
Short-read DNA libraries and re-sequencing 8
A preliminary mitochondrial assembly 8
**Pre-processing of long-read DNA data** 9
**RNA processing and transcriptome assembly** 9
Long-read cDNA processing 9
Short-read RNA-seq processing and transcriptome assembly 10
**Mitochondrial re-assembly** 10
**Genome assembly** 11
Genome assembly 11
Assessment of genome assembly quality and completeness 11
Assembly v1: Production of a preliminary assembly with the wtdbg2 assembler 11
Assembly v2: Long-read polishing with Racon 12
Assembly v3: Long-read polishing with Medaka 12
Assembly v4: Short-read polishing with Pilon 12
Assembly v5: Purging haplotigs with Purge_haplotigs 12
Assembly v6: Mixed-read polishing with HyPo 13
Assembly v7: Scaffolding the contigs with Trinity transcripts using L_RNA_Scaf 14
Assembly v8: Scaffolding with short-read RNA-seq data with a first pass of BESST_RNA 14
Assembly v9: Scaffolding with long-read DNA data using FAST-SG+ScaffMatch 14
Assembly v10: Scaffolding with linked-read DNA data using Scaf10x 15
Assembly v11: Scaffolding with short-read RNA-seq data with a second pass of BESST_RNA 16
Assembly v12: Short-read polishing with Pilon using high-coverage RNA-seq data 16

Assembly v13: Finishing the assembly by removing contaminants and re-inserting contigs 16
Reintroduction of krill genes in to the main assembly 16
Detection of contaminants 17
Flagging residual mitochondrial sequence 18
**Genome annotation 18**
Repeat annotation 19
Detection of simple repeats and low complexity regions 19
Detection and characterization of LTRs using structural searches with LTR_Finder and LTRharvest 19
Detection of diverse transposon domains using TransposonPSI 21
Detection of transposable elements through de novo repeat assembly with dnaPipeTE 22
Detection of transposable elements from high-frequency motifs using RepeatModeler2 22
Evaluation and validation of the repeat library 23
Annotation of repeats across the krill genome with RepeatMasker 23
Gene annotation 24
RNA-based data 24
Comparative data 24
Consolidation of gene models 25
Gene set size and completeness 25
Functional and evolutionary annotations 26
Gene region masking 26
**Evolutionary analyses of protein coding genes 27**
Inference and comparison of orthologs between krill and other species 27
Estimating divergence between the Northern krill and the Antarctic krill 27
Inferences of phylogenetic interrelationships 28
Analysis of gene family evolution 28
Preparing the krill gene set 29
Analyses with SwiftOrtho to cluster genes into families 29
Analyses with CAFE to trace gene family expansions or contractions 30
Gene ontology (GO) enrichment analysis of expanded gene families 31
Analyses with wgd to test for signatures of whole-genome duplication (WGD) 31
Characterization of the Hox gene complement 32
Characterization of genes involved in DNA methylation 32
Predicting protein structures 33
**Analysis of DNA methylation levels 33**
**Population genetic data processing and analyses 34**
Processing re-sequencing data 34
Mapping, read-group tagging and duplicate marking datasets 34

Mapping-depth profiles and masking of inaccessible sites 35
Calling and phasing single-nucleotide polymorphisms (SNPs) across the genome 35
Calling and processing variants 35
Imputing and phasing SNPs 36
Annotating SNPs 36
Estimating patterns of variation 37
Levels of variation 37
Effective population size (NE) and its historical demographic trends 37
Counting alleles and estimating allele frequency divergence, population structure and selection 39
Simulations of divergence 40
Signatures of selective sweeps 41
Enrichment analyses 42
Estimation of haplotype ages 42
Assessment of molecular evolution in nrf-6 and the topology of its encoded protein 43
**Supplementary Text 45**
Geographical variation in the Northern krill and regions used to sample material 45
Gulf of Maine (USA) - The SW North Atlantic Ocean 45
Gulf of Saint Lawrence (Canada) - The SW North Atlantic Ocean 45
Masfjord (Norway) - The NE North Atlantic Ocean 46
Gullmar Fjord (Sweden) - The NE North Atlantic Ocean 46
Iceland - The NE North Atlantic Ocean 47
Barents Sea & Svalbard - The NE North Atlantic Ocean 47
Spain - The Mediterranean Sea 47
**No signature of whole-genome duplication (WGD) in the Northern krill 48**
**Supplementary figures 49**
**Supplementary tables 86**
Table S1. See separate file. 86
Table S2. 87
Table S3. See separate file. 88
Table S4. 89
Table S5. See separate file. 90
Table S6. 91
Table S7. See separate file. 92
Table S8. 93
Table S9. 94
Table S10. See separate file. 95
Table S11. See separate file. 95

Table S12. See separate file. 95
Table S13. See separate file. 95
Table S14. See separate file. 95
Table S15. See separate file. 95
**References** **96**

### Materials and Methods

#### Biological materials

Adult specimens of the Northern krill (*Meganyctiphanes norvegica*) were collected from eight geographical regions across the North Atlantic Ocean and the Mediterranean Sea by different collaborators and co-authors (table S1; Fig 1A; fig. S19C). Environmental characteristics and sampling at each location is provided as Supplementary Text, while sample details including sampling coordinates and sequencing yields are in table S1.

#### *Processing of samples*

Live specimens were either preserved in 95-99% EtOH or in RNA*later*™ (Invitrogen™) to facilitate either DNA and RNA recovery. Some specimens were split behind the carapace so that the cephalothorax and abdomen could be stored separately in either buffer. For most samples, the sex was determined by inspecting petasma or thelycum for male or female characteristics using a stereo microscope. This was either done at the time of collection or after preservation in EtOH. Altogether, 36 samples were determined to be males, 32 to be females. Seven samples were not sexed because the relevant tissue had been removed or damaged in processing.

A single female specimen (Sample IDs “K20” or “swe_1”; ~3cm long) collected in the Gullmarsfjord in Sweden was used to produce all primary long-read and linked-read DNA data for the main *de novo* genome assembly, as well as most novel short-read and long-read RNA/cDNA resources used for scaffolding and annotation of the genome sequence. In order to reduce the risk of contamination from food-particles, this specimen and others sampled at the same location were kept in aquaria with filtered running deep water at ambient temperatures (10°C) at the Kristineberg Marine Research Station without access to food for 24–48 hrs before they were preserved. Sequences from other specimens were used to assist in specific tasks for the genome assembly (table S2; fig. S2), as well as to produce a comprehensive population dataset to map genetic variation.

#### Overview of library preparation & sequencing

For the genome assembly and annotation, we produced DNA and RNA/cDNA data using Oxford Nanopore Technologies (ONT) MinION (hereafter: MinION) and PromethION long-read sequencing (hereafter: PromethION). Linked-read DNA data was produced using 10x Genomics Chromium sequencing (hereafter: 10x). RNA was also sequenced using Illumina RNA-seq. Low-coverage whole-genome population-scale re-sequencing (WGS) was carried out using Nextera DNA Flex libraries or Illumina DNA PCR-Free libraries. All DNA/RNA extractions and the MinION sequencing were performed at the Dept. of Medical Biochemistry and Microbiology at Uppsala

University. All other library preparation and sequencing was carried out by the National Genomics Infrastructure (NGI) at Science for Life Laboratory (SciLifeLab), Sweden. Extractions were quantified for purity, yield and fragment lengths using the Thermo Scientific NanoDrop and Qubit systems, standard 1% agarose gels, and Agilent TapeStation and Femto Pulse systems, respectively. Summary reports with basic sequence statistics were produced using `PycoQC` for long-reads (*1*) and `MultiQC` for short-reads (*2*) with the high-throughput `FastQC` backend (Simon Andrews, Babraham Institute in Cambridge), respectively, according to standard procedure.

#### *High-molecular weight DNA extraction for long-read and linked-read libraries*

We produced DNA data using PromethION and 10x sequencing of a single female specimen for the genome assembly. We dissected five abdominal segments for all ethanol-preserved tail muscle tissue and extracted each segment individually (K20-1 through K20-5). Each piece was snap frozen in liquid nitrogen and grinded using a pestle and mortar. High-molecular weight (HMW) DNA was extracted using the Genomic-tip 100/G Midi kit (Qiagen) according to the manufacturer's protocol for tissues. Briefly, the pieces of tissue were lysed in presence of Protease (Qiagen) and RNAse A (Qiagen) to degrade proteins and RNA, respectively. The DNA was bound to the Genomic-tip resin, purified from RNA, protein and other contaminants, eluted and desalted using isopropanol precipitation. The DNA extractions were finally eluted in 70 µL of TE buffer (Qiagen AE buffer). The extraction yielded approximately 30 µg DNA (Qubit), with a peak of DNA fragment lengths at about 130 to 180 kbp, and with similar yields and qualities across the five abdominal segments (fig. S1A–B).

#### *PromethION long-read sequencing*

The ONT SQK-LSK109 Ligation Sequencing Kit was used by NGI to prepare DNA libraries using all five extractions (K20 1–5; 90% of DNA was consumed), which were then sequenced across ten PromethION FLO-PRO002 R9.4 flow cells. The DNA was sheared to 20 or 30 kb fragments with a Diagenode Diagnostics Megaruptor before preparing most remaining libraries to find an optimal balance and tradeoff between read lengths and yields (fig. S1C). One HMW library was made with the Megaruptor set to shear to 75 kb, followed by size selection for fragments >15 kb using the Sage Science BluePippin system. Library preparation for nanopore sequencing was done using the LSK-109 kit, with the following changes to the original protocol: i) End-prep / FFPE incubation times at 20 and 65 degrees were prolonged up to 10 or 15 minutes, and ii) adapter ligation time was prolonged to up to 55 minutes. Sequencing was performed on a PromethION beta machine, using R.9.4 flow cells.. In order to increase the overall yield, we reused the flow cells. We first washed them using standard wash kit buffers and then performed nuclease flushes to digest old libraries, before applying new DNA. Two flushed flow cells were run with cDNA libraries instead of DNA (see below).

Sequencing was performed using ONT `MinKNOW v3.1.18`. Initial basecalls were carried out using MinKNOW/Albacore but were updated with calls using ONT `Guppy v2.3.7` and the more accurate "flip-flop" algorithm (the dna_r9.4.1_450bps_flipflop_prom model) (*3*). The updated basecalls produced 593.8 Gb of DNA data (13% produced on flushed cells), representing 13% more data than the original basecalls. The Guppy-recall took 10 days of GPU-compute time across four NVIDIA 1080 GPUs. We sequenced 84.4 M reads with overall $N_{50}$=12 kb. About 57% of sequences had mean base qualities (Q-scores) above 10 using flip-flop (table S3), compared to 16% in the original call. Assuming a genome size of 18.6 Gbp (*4*), the PromethION data corresponded to 32× coverage across the genome, with ~20× coverage of reads 10 kb or longer (table S3; fig. S1D).

#### *10X linked-read sequencing*

Four separate 10x GEM libraries were made from the K20-5 HMW DNA extraction by NGI following the manufacturer's recommended protocol and had mean library fragment lengths of 530–540 bp. Paired-end reads (2×150 bp) were sequenced on a single Illumina NovaSeq 6000 S4 lane, generating 897 Gb of data across 2.97 B read-pairs (~46× coverage; 12.6% marked as duplicates; %GC=34.7). The standard `LongRanger` (v2.2.2; 10x Genomics) "`basic`" command was used to extract the 16 bp barcodes from the forward/R1 reads for each library individually. About 3.5 M of the 4.8 M barcodes (4M-with-alts-february-2016.txt; 10x Genomics) were found to be associated with at least one read in each library and ~96% of all sequences were barcoded. We rewrote the "BX:Z" style barcodes of each of the four libraries using a unique suffix ("_1" through "_4") in order to keep them separate in downstream analyses and to de-interleave the FASTQ files.

#### *RNA extraction from multiple tissues for long-read and short-read RNA libraries*

We produced RNA data using PromethION/minION cDNA-sequencing and Illumina short-read RNA-seq using the K20 specimen for gene and genome annotation. Six libraries were prepared using RNA from small pieces of tissue preserved in RNAlater: i) head/brain; ii) dorsal/caudal muscle tissue on cephalothorax; iii) thoracic exopodite legs (distal segments of the periopods); iv) ventral soft tissue of cephalothorax including gills; v) hepatopancreas (the crustacean digestive gland); and vi) soft-tissue surrounding it. RNA was extracted using the RNeasy Lipid Tissue Mini Kit (Qiagen) according to the standard protocol, using ceramic beads and a BioSpec Products Mini-Beadbeater for tissue homogenization, and eluted in 30 µL RNAse-free water. This yielded RNA extractions with concentrations between 50–240 ng/µL and with RIN-values between 9.1–10. We used both raw RNA-seq reads and Trinity assembled transcripts (*5*) in downstream analyses.

#### *Short-read RNA sequencing*

NGI produced six libraries from the six K20 RNA extractions using Illumina TruSeq Stranded mRNA Library Prep kit with polyA selection, which were sequenced as 2×150 bp paired-end reads on one Illumina NovaSeq 6000 SP lane. This generated 459 M read-pairs that were demultiplexed by NGI (table S3).

#### *Long-read cDNA sequencing*

One barcoded and amplified Nanopore cDNA library was produced from each of the six K20 RNA extractions, using the PCR-based ONT Strand-switching protocol with poly-A selection and the LSK109 ligation kit to add barcodes and adapters. We performed cDNA sequencing on PromethION (ONT `MinKNOW v3.1.18`) using two nuclease flushed flow cells and one new FLO-MIN106D R9.4.1 MinION flow cell (3 M reads) and the standard loading kit and buffers for each instrument. The data was basecalled using ONT Guppy v4.4.1 and HAC (high accuracy) model (dna_r9.4.1_450bps_hac_prom.cfg). The two PromethION runs generated 8.99 M and 3.91 M reads (10.1 + 5.2 Gb of cDNA data), with median Q-scores of 10.5 and 10.7 and median lengths of 962 bp and 1,060 bp, respectively. The MinION run produced 3.0 M reads (3.5 Gb of cDNA data) with median Q-score of 12.6 and median lengths of 915 bp.

#### *Salt-isopropanol DNA extraction of multiple specimens for population-scale whole-genome re-sequencing*

To produce the population dataset, we extracted DNA from 75 *M. norvegica* specimens using a salt-isopropanol precipitation protocol. We extracted DNA from abdominal muscle tissue from 1–2 muscle segments of krill samples. Samples were selected based on geographic origin and quality of preservation. We sequenced 6–10 specimens per population. We used a column-free salt-isopropanol precipitation protocol to extract DNA. Briefly, tissues were lysed over night in ATL Lysis buffer (Qiagen) together with Proteinase K (Qiagen) over night, followed by RNAse A (Qiagen) treatment, protein separation using Protein Precipitation Solution (Qiagen) and precipitation and washing of the DNA using isopropanol and ethanol, respectively. The washed DNA pellet was finally eluted in 70 µL MilliQ-water. The extractions produced about 7 µg of DNA on average per specimen (Qubit). The protocol is available as a separate document in this publication and at the online platform protocols.io:

- https://www.protocols.io/workspaces/wallberg.lab.imbim.uu/resources/isopropanol-hmw-dna-extraction-method

#### *Short-read DNA libraries and re-sequencing*

Barcoded Nextera DNA Flex or Illumina DNA PCR-Free libraries (Illumina) were prepared from 75 DNA extractions (74 specimens for re-sequencing + the reference specimen) by NGI/SciLifeLab according to the manufacturer's instructions. Both protocols use a bead-linked transposome to fragment DNA and incorporate adapters in a single step. About 350 ng of DNA was used as input per Nextera library and 300 ng per PCR-Free library. We aimed for ~3× depth of coverage per specimen and sequenced the libraries across six Illumina NovaSeq 6000 S4 lanes, generating 15.1 B 2x150 bp read-pairs with valid library barcodes (18% or 23% of reads marked as duplicates with Nextera or DNA PCR-Free, respectively; %GC=32.9; 88% of reads with Q-scores above 30). One library ("sva_1" from the Svalbard area) failed to sequence (0.03× coverage; table S1) and was discarded from population genetic analyses (hence 74 samples were carried forward).

#### *A preliminary mitochondrial assembly*

A minION test run of specimen K4 from the Gullmarsfjord produced 1.5 Gbp of long-read across 270,185 reads (N50=8,229 bp; <0.1× across the genome) ahead of the other sequencing runs. We downloaded mitochondrial *M. norvegica* template accessions (16S=AY744910; COI=FJ581747; CYTB=AF149775; NADH=AF149775; tRNA_ser=AF149775; tRNA_leu_16S.AF149775) and used BLASTN 2.2.31+ to identify 235 putatively mitochondrial reads shorter than 16 kbp that aligned to any of the templates. These reads were assembled in

`Canu v1.7` (*6*) with "`genomeSize=17k`" that produced a 24.8 kbp contig. We blasted the read pool against this contig and collected additional reads (n=520 in total) and performed a second round of assembly with Canu, generating a new 24.802 bp contig from 83 aligned reads (37× depth of coverage) that the assembler suggested was circular. Manual inspection of the sequence indicated that it was the full mitochondrial chromosome partly repeating itself and we circularized the sequence by hand, producing a 17,360 bp long sequence. We next mapped the long-reads to the mitochondrial sequence with `minimap2 v2.17-r941` (*7*) (109× mean depth of coverage) and polished it with `Nanopolish v0.11.3` (*8*) using the "`variants`" and "`vcf2fasta`" subcommands. We took the high coverage as an indicator of high mitochondrial:nuclear ratio in the extracted muscle tissue. Finally, we mapped RNA-seq data from the published TI787-MN-F specimen to the polished sequence using `Bowtie2 v2.3.4.1` (*9*) and performed one round of polishing with `Pilon v1.22` (*10*) with the "`--fix indels,gaps,local`" setting. The resulting mitochondrial sequence was 17,508 bp with %GC=28. Automated annotation was performed with the online MITOS2 tool (*11*). We used this sequence to screen out potential mitochondrial long-reads from the K20 reference datasets ahead of genome assembly and used a similar approach to assemble the mitochondrial chromosome from this specimen (see mitochondrial re-assembly below).

**Pre-processing of long-read DNA data**

We used `Porechop v0.2.4` (*3*) to trim the LSK109/NSK007 top and bottom adapters from the PromethION long-reads. Reads with internal adapters can be chimeras and either be split or removed. We first ran the program without splitting potentially chimeric reads:

```
porechop -t 20 --no_split --adapter_threshold 60 --end_threshold 70 --end_size
200 --in fastq.gz --out fastq.trimmed.gz
```

We then repeated the scan without "`--no_split`" to note which reads had middle adapters (n=50,368; 0.06% of all reads). We next mapped all reads against our preliminary mitochondrial sequence with `minimap2 v2.17-r941` (*7*) and found that 382 K reads mapped to this chromosome for a mean depth coverage of 33,662× (vs about 32× across the nuclear genome). We filtered out both the putative chimeric and mitochondrial long-reads for the reference PromethION FASTQ read-pool, to reduce the risk of downstream interference in the main genome assembly.

**RNA processing and transcriptome assembly**

#### *Long-read cDNA processing*

The barcoded ONT PromethION and minION cDNA reads were de-multiplexed with ONT `guppy_barcoder v4.4.1` and then processed with ONT `Pychopper v2.5.0` (https://github.com/nanoporetech/pychopper) to find, orient and trim full-length cDNA reads using the SSP/NVP primer sequences (cDNA_SSP_VNP.fas). The resulting 8.7 M reads were combined into a single archive and later used for gene annotation (see below). We also

produced a non-redundant set of sequences to represent potential genes in the krill genome. Here we extracted long and high-quality cDNA reads (length>500bp and Q>12) from both runs and clustered them into putative genes with `isONclust2` (*12*) using the "`-x sahlin`" mode. We then error-corrected the clustered cDNA reads with `isONcorrect v0.0.6` (*13*). Lastly, for each error-corrected cluster with four or more cDNA reads we generated a consensus sequence with
`vsearch v2.14.1` (*14*):

```
vsearch --cluster_fast <clusters.fa> --threads 40 --id 0.95 --clusterout_id --
clusterout_sort --sizeorder --sizeout --consout <clusters.fa.cons>
```

This produced 24,632 consensus sequences with $N_{50}$=1,810 bp. These non-redundant transcript sequences were later used for RNA-based scaffolding (see below).

#### *Short-read RNA-seq processing and transcriptome assembly*

The Illumina RNA-seq data from the reference specimen and two published RNA-seq archives, TI787-MN-M (adult male; SRR3657321) and TI787-MN-F (adult female; SRR3657320), were trimmed for adapters and base qualities using `Trim Galore! v0.6.1` (https://github.com/FelixKrueger/TrimGalore/) and `Cutadapt v2.3` (*15*):

```
trim_galore -j 8 --gzip --length 50 --trim-n --paired
```

All read pairs and 89–99% of bases were retained. The trimmed reads were later used in genome assembly scaffolding and annotation (see below). We used `Trinity v2.5.1` (*5*) to assemble a transcriptome from all 459 M trimmed read-pairs from the reference specimen, producing 573,869 with $N_{50}$=900 bp. Using `BUSCO v3.0.2b` (*16*) with the Arthropoda odb9 lineage set, we detected almost all expected genes (C:97.2%[S:42.4%,D:54.8%], F:1.6%, M:1.2%, n:1066). A subset of 500 bp or longer transcripts were used in scaffolding (see below). We processed all transcripts with `Trinotate v3.1.1` (*17*) and identified 60,677 possibly coding transcripts with best hits against metazoan templates from parsing the "Trinotate.xls" output and used these transcripts for gene annotation (see below). A subset of 16,509 non-redundant transcripts with best hits against arthropods were used to measure assembly quality scores (see below).

#### Mitochondrial re-assembly

We re–assembled the mitochondrial chromosome for the K20 reference specimen similarly as described above. PromethION reads identified during the screening process and that were also between 1–16 kb long were used to assemble the chromosome in `Canu v2.0-development` (*6*) with "`genomeSize=17k`" that produced a 26.1 kb contig that the assembler suggested was circular. We next mapped the 369,946 mitochondrial long-reads to the sequence with `minimap2 v2.17-r941` (*7*) and polished it with `Racon` using the "`variants`" and "`vcf2fasta`" subcommands. We circularized the sequence manually, producing a mitochondrial sequence that was 17,840 bp. We then mapped a subset of the 10x Chromium data (656,771 read pairs) to the chromosome with BWA and performed one round of polishing with `Pilon v1.23` with the "`--fix all`" setting, followed by a round of polishing with K20 RNA-seq data. The final mitochondrial sequence was 17,944 bp with %GC=28. Automated annotation was again carried out with the online MITOS2 platform as before and features were visualized with `DNAPlotter` (*18*) (fig S3).

**Genome assembly**

The hybrid genome assembly underwent multiple steps of polishing, scaffolding and refinements, in which we prioritized achieving high gene completeness and gene contiguity. We mainly used genetic information from the reference specimen, but also included data from other specimens for specific tasks (such as identifying redundant haplotigs). Nanopore long-reads were used for assembly, polishing (reducing base-level errors) and scaffolding, while 10x barcoded linked-reads and RNA data were used for polishing (in highly expressed regions) and scaffolding.

#### *Genome assembly*

Scaffolding was performed with top priority on recovering gene bodies using RNA from the reference specimen and additional RNA-libraries (*19*), followed by genome-wide scaffolding using Nanopore synthetic mate-pairs with `FAST-SG` (*20*) and 10x Chromium barcodes with `Scaf10x`. Some inadvertently removed contigs with genes were manually re-inserted into the genome. Screens against the SILVA ribosomal database (*21*) were used to isolate contigs from bacterial contaminants. The assembly was carried out in 13 steps and we used `BUSCO` (*22*) and `genecovr` (https://github.com/NBISweden/genecovr) throughout the pipeline to assess gene detection, completeness and sequence error.

#### Assessment of genome assembly quality and completeness

We used `BUSCO v3.0.2b` with the Arthropoda odb9 lineage set or `BUSCO v5.0.0` (*22*) with the Arthropoda odb10 lineage set, respectively, to track improvements in gene detection rates between genome assembly versions, and to assess the overall completeness and levels of duplication in the final genome. In addition, we assessed overall gene completeness by mapping a set of 16,509 trinity transcripts to the assemblies using `gmap` version 2020-06-30 (*23*). We parsed the psl output and summarized various statistics related to mapping quality, including number of insertions, number of mismatches, number of distinct contig hits, alignment length and alignment depth, to obtain "gene coverage" statistics for each transcript. Compilation of the statistical summaries and plots were implemented in a custom R package `genecovr`:

- https://github.com/NBISweden/genecovr.

#### Assembly v1: Production of a preliminary assembly with the wtdbg2 assembler

To assemble the krill genome, we used the `wtdbg2 v2.5` assembler (*24*) and 6–300 kbp long reads with Q-scores ≥10 selected with NanoFilt v2.6.0 (*25*) and that had been pre-screened for internal adapters or mitochondrial sequence, corresponding to ~16× depth of coverage across the genome. We used a short kmer setting (“`-p 17`”; as recommended for noisy data by the developers) and sparse kmer subsampling (“`-S 2`”) for sensitive read analysis and binning. Assembling the genome using these settings (“`-A -p 17 -S 2 -s 0.05 -L 6000  -g 18g -t 168`”) took 16 days on a 168 core machine with 4.7 TB RAM. Consensus contigs were exported with `wtdbg-cns`, resulting in genome assembly v1, which spanned 21.8 Gbp across 797,490 contigs with $N_{50}$=41 kbp.

#### Assembly v2: Long-read polishing with Racon

The PromethION long-reads were mapped in parallel chunks to the v1 genome assembly using `minimap2 v2.17-r941` set to use a split prefix and without secondary alignments (“`-L -ax map-ont -y -t 16 --secondary=no --split-prefix`”). Data was first saved as gzipped SAM files and then sorted according to coordinates into BAM files with `samtools v1.10` (*26*), followed by merging into a single BAM file with samtools. The genome assembly was split into 248 100 Mbp FASTA chunks with a custom Perl script and the mapped reads were partitioned into corresponding SAM and FASTQ files using samtools. We used one round of `Racon v1.4.11` with the average base quality threshold set to 8 (“`-q 8`”) to polish all 248 chunks in parallel. The polished contigs were concatenated into genome assembly v2.

#### Assembly v3: Long-read polishing with Medaka

We followed the guidelines from the medaka source documentation (https://github.com/nanoporetech/medaka#improving-parallelism) that suggests running the medaka components independently and in parallel for large genomes. Briefly, the PromethION long-reads were mapped in 250 parallel chunks to the v2 genome assembly using `minimap2 v2.17-r941` with parameters “`-x map-ont --MD --secondary=no -t1`”. This was followed by the consensus algorithm `medaka consensus` with parameters “`--model r941_flip235 --batch_size 100 --threads 1`” to generate models over each assembly chunk. Finally, consensus sequences were generated with `medaka stitch` and merged to obtain assembly v3.

#### Assembly v4: Short-read polishing with Pilon

The 10x Chromium paired-end linked-reads were contained in 16 paired FASTQ libraries that were mapped to the v3 genome assembly with `BWA v0.7.17` (*27, 28*). First, the genome reference sequence was indexed (“`bwa index -a bwtsw`”) and the data was then mapped with the “`bwa mem`” algorithm, piping the output to samtools which sorted the data on the fly before saving a BAM to disk. The 16 BAMs were merged with `samtools`. As in the Racon step above, both the reference genome and the BAM were split according to chunks of contigs spanning 100 Mbp. Polishing was performed in parallel with `Pilon v1.23` set to polish putative base-level errors and specifying a diploid dataset (“`--fix snps,indels --diploid`”). The short-read polished contigs were concatenated into genome assembly v4.

#### Assembly v5: Purging haplotigs with Purge_haplotigs

Since our genome assembly was about 3 Gbp larger than the expected genome size (21.8 Gbp vs ~19 Gbp), we next sought to identify and remove potential haplotigs from the assembly using `Purge_haplotigs v1.1.0` (*29*). The tool uses sequence similarity among contigs as well as a bimodal signature of read mapping depths to isolate and remove alternative alleles of the same locus from highly heterozygous diploid genome assemblies. It also identifies "junk" contigs with extreme mapping depths (low or high) that may be assembly artifacts or otherwise intractable in downstream analyses.

We first mapped the ~32× PromethION long-reads using `Minimap2` to the v4 assembly, as in previous steps, and then used the "`purge_haplotigs hist`" command to generate a depth-of-coverage histogram across the genome. However, this relatively low-coverage dataset did not result in a clearly bimodal mapping profile with 1x and 0.5x mapping peaks on its own. We therefore also mapped the 10x Chromium linked-reads and a preliminary set of ~140× population re-sequencing data to the genome using `BWA`, in order to use all available data to specify depth thresholds to flag offending contigs. The short-read BAMs were marked for duplicates with `Picard MarkDuplicates`. We then exported per-base depth-of-coverage data running "`samtools depth`" across all positions in the genome assembly ("`-a`") and generated a depth histogram with a discernable "1x" peak at ~150×, presumably corresponding to properly haplotype-fused contigs. The expected position of the 0.5x peak at half of this depth was less clear and possibly intermixed with genomic regions of low mappability. Given the depth profile, we then configured `Purge_haplotigs` to use 20× and 240× as lower and upper cut-offs, respectively, and 125× as the mid-point. We specified that contigs with 100% or less of their depths at diploid level of coverage to be flagged as suspected haplotigs (i.e. to comprehensively evaluate all contigs) and that those that had depths outside of the upper/lower cut-offs across >50% of their lengths to be flagged as junk:

```
purge_haplotigs cov -l 20 -m 125 -h 240 -s 100 -j 50 -i <coverage_file> -l 20
-m 125 -h 240 -o <coverage_stats_output>
```

Purge_haplotigs infers contig similarities from the proportion of sequence that can be aligned between two contigs using minimap2, and can be set to disregard alignments across repeat sequence. We therefore produced a preliminary catalog of repeat sequence intervals in BED format using `Red` (*30*), masking 11.7 Gb (54%) of the genome. We used the default alignment score cut-off (≥ 70%) to mark contigs for reassignment as haplotigs:

```
purge_haplotigs purge -align_cov 70 -repeats <repeats_file>
```

The purge subcommand executes the script "`purge.pl`". We modified this script to increase performance in the handling of our large, repeated and relatively fragmented genome assembly by: implementing faster parsing of tabular data; a string-based instead of array based data structure for hit summaries; and faster internal repeat-handling instead of executing the external tool bedtools from within the script. This code is available at: [https://github.com/andreaswallberg/popgenome-tools/](https://github.com/andreaswallberg/popgenome-tools/).

Using these settings, we purged 168,316 contigs ($N_{50}$=13,074 bp) spanning 1.95 Gbp as potential haplotigs and 49,996 contigs ($N_{50}$=14,236 bp) spanning 565 Mbp as junk/artifact contigs, respectively, removing 12% of the original genome assembly. The remaining main assembly had 586,516 contigs ($N_{50}$=46,208 bp) spanning 19.18 Gbp and was taken as genome assembly v5.

#### Assembly v6: Mixed-read polishing with HyPo

We aimed to polish the genome further after purging haplotigs and mapped both the PromethION long-reads and 10x Chromium linked-reads from the reference specimen back to the genome as above. We next split the genome assembly, BAMs and FASTQ reads (using samtools) into ten chunks of contigs spanning 2 Gbp of the genome each and ran the polisher `HyPo v1.0.3` (*31*) separately on each chunk, which uses both short and long reads to polish contigs in a single run. The polished contigs were taken as genome assembly v6.

#### Assembly v7: Scaffolding the contigs with Trinity transcripts using L_RNA_Scaf

At this stage, quality assessments indicated that we had many broken gene models with exons distributed across more than one contig. Preliminary trials suggested that scaffolding contigs together based on information in RNA-mappings before long-read or linked-read DNA scaffolding recovered more complete gene models for this genome than vice-versa. We started by scaffolding contigs with non-redundant full-length cDNA transcripts (n=24,632; see section above) and Trinity-assembled transcripts, i.e. contiguous RNA evidence that was assembled independently from the genome assembly itself, using `L_RNA_Scaffolder` (*32*). We first filtered the Trinity transcripts to only include the longest isoform of each gene and kept only the longest isoforms that were >500bp long (n=73,422; $N_{50}$=1,482 bp). L_RNA_Scaffolder typically uses BLAT to infer query alignments, which does not support genomes larger than 4 Gbp. We therefore instead used `NCBI BLAST v2.2.31+` (*33*) to build a database:

```
makeblastdb -dbtype nucl -in <reference>
```

We then queried both sets of transcripts using `MEGABLAST`:

```
blastn -query <transcripts.fa> -task megablast -db <reference> -outfmt 5 -
perc_identity 80 -max_target_seqs 100 -num_threads 36
```

We then converted the XML output into the expected PSL format using UCSC `blastXmlToPsl v412` (http://hgdownload.soe.ucsc.edu/admin/exe/). L_RNA_Scaffolder scaffolding generated 12,277 scaffolds from joining 20,641 contigs. The scaffolds had $N_{50}$=158,477 bp, compared to $N_{50}$=44,758 bp among the unscaffolded contigs (n=553,073). Scaffolds and contigs were joined into genome assembly v7.

#### Assembly v8: Scaffolding with short-read RNA-seq data with a first pass of BESST_RNA

We next used `BESST_RNA` (https://github.com/ksahlin/BESST_RNA) to perform additional RNA-based scaffolding using paired-end RNA-seq data. We first mapped the 755 M processed

Illumina short-read RNA-seq read-pairs derived from the reference specimen itself and two published specimens (one male with ID TI787-MN-M; one female with ID TI787-MN-M; table S2; fig. S2) (*19*) against the assembly with `HISAT2 v2.2.1` (*34*) (mapping rates 81–86%) and merge the data into a single BAM file using samtools. We then performed scaffolding with `BESST_RNA` requiring three or more witness links with a maximum of 200,000 bp distance (“`-e 3 -T 200000 -k 500 -d 1 -z 10000 -g 1`”), which resulted in a total of 15,519 RNA-based scaffolds (including those from the previous step) and reduced the number of sequences to 557,087 in genome assembly v8.

#### Assembly v9: Scaffolding with long-read DNA data using FAST-SG+ScaffMatch

We hypothesized that there could be residual unresolved long-range contact information in the PromethION DNA data and used the `Fast-SG` scaffolder (*20*) with `KMC v3.0.0` (*35*) for k-mer counting to accomplish further scaffolding. First, we extracted a subset of long-reads ≥ 5 kbp and with average Q-scores above 10 from the read pool. Next, we generated synthetic mate-pairs in Fast-SG with insert lengths ranging from 4,000 to 40,000 bp (n=10) using 32bp kmer matches spaced along reads and the genome to enable downstream detection links between contigs. The mate-pair settings were specified in a single-line read configuration file:

```
long ont <reads.fq> 4000,5000,6000,7000,8000,10000,15000,20000,30000,40000 1
```

Fast-SG was run as:

```
FAST-SG.pl -k 32 -l <read_configuration.txt> -r <genome_assembly.fa> -p
results -c $KMC -t 40
```

This produced two SAM files per synthetic mate-pair group, containing perfect forward/reversed kmer matches along reads and contigs at the approximate insert lengths specified for each insert group. The empirical mean and standard deviation of the insert lengths was computed from the first 1 M observations in each group as per the FAST-SG manual (https://github.com/adigenova/fast-sg/wiki/Hybrid-scaffolding-of-NA12878).

The statistics and SAM mappings were next processed in `ScaffMatch v0.9` (*36*) to generate and resolve a scaffolding graph in order to scaffold the genome assembly:

```
python2.7 $PATH/scaffmatch.py -w ONT-K32 -c <genome_assembly.fa> \
            -s 1461,1734,1907,2137,2194,2646,3736,7711,10503,13425 \ # std-dev
            -i 3957,4964,5965,6958,7938,9921,14634,16513,26175,34772 \ # mean
            -p fr,fr,fr,fr,fr,fr,fr,fr,fr,fr \
            -m1 <ont.I4000.fwd.sam>, <ont.I5000.fwd.sam>, … \
            -m2 <ont.I4000.rev.sam>, <ont.I5000.rev.sam>, … \
             2>&1 | tee scaffmatch.log
```

Fast-SG+ScaffMatch produced a scaffolded genome assembly spanning 247,751 sequences ($N_{50}$=142,666 bp), out of which 131,888 were scaffolds and 115,863 were contigs, and was taken as genome assembly v9.

#### Assembly v10: Scaffolding with linked-read DNA data using Scaf10x

Having produced scaffolds with RNA data and long-reads and reduced the overall number of sequences in the genome assembly, we next explored the possibility to scaffold sequences based on signatures of shared 10x Chromium linked-read barcodes between the outer edge regions of scaffolds and contigs. Scaffolding was performed using `Scaff10X v4.2` (https://github.com/wtsi-hpag/Scaff10X).

First, we mapped the linked-reads to the genome assembly using BWA, as in previous steps. For performance and memory reasons, we next pre-filtered the BAM file to only include reads mapping to the outer 20 kbp regions of scaffolds or contigs using samtools and a custom Perl script. To reduce the influence of ambiguous mappings to highly repetitive sequences in the outer regions, we also filtered the reads to only include read-pair mappings with MAPQ≥20 in the same script. We then ran one iteration of Scaff10x, and configured the program to require at least six reads per barcode ("`-reads 6`"), eight shared barcodes as evidence of links between sequences ("`-link 8`"), edge length to match our estimated linked-read lengths ("`-edge 20000`"), with other settings at their default values:

```
scaff10x -nodes 20 -gap 100 -longread 1 -reads 6 -link 8 -edge 20000 -plot
<mappings.bam>.png -bam <mappings.bam> <genome_assembly.fa>
```

This resulted in scaffolded genome assembly v10, spanning 219,207 sequences with $N_{50}$=214,496 bp, including 108,310 scaffolds.

#### Assembly v11: Scaffolding with short-read RNA-seq data with a second pass of BESST_RNA

After scaffolding the genome assembly with long-read and linked-read DNA data, we applied a second round of RNA scaffolding using `BESST_RNA` and RNA-seq data, as above, to find gene-based links that were not resolved in previous steps. `BESST_RNA` reported that 2,294 new scaffolds had been formed. This round of scaffolding further reduced the number of sequences from 219,207 in v10 to 216,722 ($N_{50}$=220,983 bp) in genome assembly v11, out of which 106,444 sequences were scaffolds.

#### Assembly v12: Short-read polishing with Pilon using high-coverage RNA-seq data

The Illumina RNA-seq data from the reference specimen was re-mapped to the genome assembly (88.3% mapping rate) and one final round of base-level polishing Pilon ("`--fix snps,indels`") was applied across transcribed regions with at least 100× depth of coverage ("`--mindepth 100`"), resulting in sequences with minor adjustments that were taken as genome assembly v12.

### Assembly v13: Finishing the assembly by removing contaminants and re-inserting contigs

Some qualitative adjustments were carried out to reassign sequences between the main genome assembly and other classes of sequence in order to maximize gene completeness and remove putative non-krill contaminations and mitochondrial artifacts. We screened the main assembly (v12), as well as the "artifact" (~565 Mb) and "haploid" (~1.95 Gb) sequences classified by purge haplotigs (produced in genome assembly v5) to identify candidate sequence for removal or inclusion.

#### Reintroduction of krill genes in to the main assembly

We identified missing genes in the main assembly using both comparative and experimental evidence. By running `BUSCO` on the artifacts and haplotigs we identified two complete single-copy BUSCO genes that had no corresponding BUSCO `tblastn` hit in the main assembly. In addition, we used `GMAP` to identify ten contigs from the haplotig class that had unique Trinity transcripts mapped with high quality (>90% identity across >90% of the transcript) but with no hits in the main assembly. We considered these 12 contigs to have been inadvertently misclassified and transferred them back into the main assembly.

Ribosomal gene sequences occur in high copy numbers in many metazoans and may be difficult to target accurately with short-read data. We therefore used the nucleotide-nucleotide `BLAST` (blastn) and the extensive SILVA LSU and SSU ribosomal RNA databases (release 132) (*21*, *37*) to identify "artifact" sequences with ribosomal loci from the krill. In the majority of contigs where hits were detected, the top scoring krill alignment was against the expected *M. norvegica* templates (LSU accession AY744900.1.2423 [2,423 bp]; SSU accessions AY781434.1.1853 [1,853 bp] or DQ900731.1.1884 [1,884 bp], respectively), although sometimes templates for other krill had better matches (typically from the Antarctic krill *Euphausia superba*), which occurred in particular when matches were short and fragmented. For example, without considering the quality and lengths of the reported hits against the main assembly and the "artifacts'', the *M. norvegica* LSU template had the best krill match in n=70/96 [73%] of cases, while the SSU templates were the best krill matches in 15/27 [56%] of cases. Requiring hits to be at least 1,500 bp with 98% identity against the template raised the on-target *M. norvegica* best-hits to 91% (n=10/11) and 100% (n=8/8) for LSU and SSU. We found that most sequences with long motifs matching *M. norvegica* ribosomal templates (>1,500bp; >90% identity) had been placed into the artifact class (LSU: $n_{artifacts}$=14 vs $n_{main}$=5; SSU: $n_{artifacts}$=10 vs $n_{main}$=5), likely due to aberrant mapping depths across these multi-copy loci. We transferred these "artifact" contigs (n=18) with long matches back into the main assembly.

#### Detection of contaminants

We searched the SILVA LSU and SSU ribosomal RNA databases to identify possible eukaryotic or bacterial contaminants using `blastn` and considering the top-scoring alignment for each assembly sequence. Among contigs flagged as artifacts, we detected long and high-identity ribosomal matches (>1,500 bp and >90% identity) against *Enterobacter ludwigii*/*cloacae*, *Lactobacillus reuteri* and *Delftia tsuruhatensis*. Querying the Nanopore sequence data itself against the SILVA database (see below), we re-classified the *Delftia* matches as *Delftia acidovorans*. We also identified several sequences in the main assembly with partial SSU low-identity hits against *Aphelidium desmodesmi* (n=204; average identity=81%), a eukaryotic endoparasite of freshwater green algae. However, this SSU accession (KY249641.1.3521 [3,521 bp]) also contains a partial LSU sequence with relatively high identity towards euphausiids (~85% identity across 550–650 bp). A combined search using both SSU and LSU simultaneously completely masked the hits against this accession in favor of *M. norvegica* LSU hits.

We downloaded the corresponding bacterial genome sequences from NCBI Genbank (CP027618.1 for *Enterobacter cloacae* [5.0 Mbp genome]; CP015408.2 *Lactobacillus reuteri* [2.0 Mbp] and CP019171.1 for *Delftia acidovorans* [6.6 Mbp], respectively). We then used blastn to screen the krill genome assembly for contigs and scaffolds matching these sequences. Among artifact contigs, we recovered multiple high-scoring alignments between assembly sequences and the *Enterobacter* (n=68; average alignment length=48 kb; average identity=92.5%) and *Delftia* (n=75; average alignment length=44 kb; average identity=92.4%) genomes, respectively. Among the sequences in the main assembly, we detected less *Enterobacter* (n=7; average alignment length=24 kb; average identity=99.2%; genome coverage=3%) and *Delftia* (n=12; average alignment length=88 bp; average identity 96.2%; genome coverage=4%). Matches against the *Lactobacillus* tended to score lower, covering only 0.6% of the bacterial genome among "artifact" sequences (n=7; average alignment length=1.9 kbp; average identity 86%) and 2% among main assembly sequences (n=17; average alignment length=2.1 kbp; average identity 97%).

We then queried up to 1 M Nanopore reads produced by each PromethION flow cell and sequencing-run (original or flushed) against each bacterial genome (in total, 14.8 M reads out of 84.4 M reads) using blastn and estimated that we had sequenced about 73 k reads from *Enterobacter* for a total of about 60 Mb or 12× average coverage across its genome, 66 k reads of *Lactobacillus* (~82 Mb; 40× coverage) and 14 k reads of *Delftia* (~180 Mbp; 27× coverage). Taken together (assuming no overlap between the datasets) we estimate that about 0.02% of reads and 0.05% of data was derived from these bacteria rather than the krill, respectively. Because of the low frequencies and sporadic appearance of reads among flow cells, we hypothesize that the bacteria were technical contaminants rather than being naturally associated with the krill itself. *Delftia acidovorans* has previously been associated with biofilm contamination of BluePippin size selection cassettes (*38*). We used a basic blastn criterion to move main and artefact assembly sequences with at least one top-scoring alignment with >90% identity towards any of the bacteria across 500 bp into a "contamination" class (*Enterobacter* n=51; *Lactobacillus* n=9; *Delftia* n=54), spanning about 98–99% of aligned sequence.

**Flagging residual mitochondrial sequence**

We had already screened the Nanopore dataset for mitochondrial reads against a draft assembly of the mitochondrial chromosome. Nevertheless, we probed the genome assembly for sequences matching the *M. norvegica* mitochondrion with >90% identity across at least 2 kb (longer than any individual mitochondrial gene) and detected 15 sequences harboring either mitochondrial genes or fragments of the AT-rich control region. Three of these were scaffolds, each of which with a single contig containing the long match but with short matches on one or more other contigs. We split these scaffolds and extracted only the offending contig while transferring the remaining contigs back into the assembly. We can not exclude that some of these sequences are nuclear mitochondrial DNA (NUMTs) rather than misassembled sequences and therefore put these sequences into their own "mitochondrial artifact" class.

**Genome annotation**

We first applied multiple tools to find simple and interspersed repeats, using structural signatures or protein domain homology for classification of transposable elements, and then produced a non-redundant repeat library to characterize the repeat landscape. To annotate genes, we generated gene models using RNA-seq data and cDNA or Trinity transcripts. We also mapped transcripts or gene sequences from seven other Malacostracans to find unexpressed genes and consolidated overlapping gene models into common loci, from which high-scoring non-redundant isoforms were selected as the canonical representatives of genes.

#### *Repeat annotation*

We aimed to build a custom non-redundant repeat library for the Northern krill and use it to estimate the distribution of repeats across the genome. We applied several specialized or general standard repeat detection pipelines to annotate simple, tandem and interspersed repeats and transposable elements (TEs) in the krill genome, using structural or homology-based searches against characteristic motifs, *de novo* detection and *de novo* assembly of repeats from short-reads. Intermediate repeat libraries were interrogated against the genome and/or for TE domains and masked or clustered to remove redundancy, before a final non-redundant interspersed repeat library was incrementally produced and used to annotate the genome. Some intermediate steps to filter or rename repeats based on the output from the respective tools were facilitated with custom Perl scripts. The procedure to detect interspersed repeats was inspired by two online protocols:

1. MAKER WIKI: http://weatherby.genetics.utah.edu/MAKER/wiki/index.php/Repeat_Library_Construction-Advanced
2. Berriman Lab Group (Sanger Institute) protocol on Protocol Exchange (*39*): https://doi.org/10.1038/protex.2018.054

### Detection of simple repeats and low complexity regions

We used `SciRoKo v3.4` (*40*) with default settings to estimate the degree of microsatellites in the genome assembly and identify common microsatellite motifs. We then used `SDUST v0.1` (https://github.com/lh3/sdust), a fast reimplementation of the symmetric DUST algorithm in NCBI Dustmasker (*41*), using defaults to estimate the proportion of simple repeats and low-complexity regions in the genome. Lastly, we used `TideHunter v1.4.4` (*42*) to detect tandem repeats with a minimum period of 6 bases that occurred in at least two copies ("`-p 6 -c 2`") and estimate the overall tandem repeat content in the genome assembly.

### Detection and characterization of LTRs using structural searches with LTR_Finder and LTRharvest

Characteristic structural features of LTR (Long Terminal Repeat) retrotransposons were searched with `LTR_Finder v1.07` (*43*) using the `LTR_FINDER_parallel` wrapper v1.1 (*44*) and `LTRharvest` (*45*). The output of each tool was processed with `LTR_retriever v2.9.0` (*46*) to remove putative low-quality false positive LTR hits. Internally, LTR_retriever used `RepeatMasker v4.1.2-p1` (*47*) and `CD-HIT v4.8.1` (*48*) ("`-cdhit [ -c 0.80 -n 5 -M 0 -aS 0.80 -G 0 -g 1 ]`"). We used a mutation rate inferred from snapping shrimp in (*49*) of $2.64e^{-9}$ substitutions per site per year ("`-u 2.64e-9`"), "`-missmax 10000`" to allow LTRs to span contig gaps, "`-notrunc`" to discard truncated LTRs and nested LTRs and "`-noanno`" to skip whole-genome LTRs annotation at this stage.

`LTR_retriever` filtering resulted in intermediate libraries with 10,740 LTRs from `LTR_Finder` (headers tagged "FIN") and 4,235 LTRs from `LTR_Harvest` (headers tagged "HAR"), respectively. We next mapped these motifs back to the genome using MEGABLAST in `RepeatMasker RMBLAST v2.11.0` and carried forward only those motifs that had two or more hits with >80% identity towards the genome across >80% of the repeat motif ("80/80" rule) (*50*). Many LTRs are autonomous and contain several genes to facilitate their retrotransposition. The LTRs in each library were therefore characterized through homology to known LTRs or LTR protein domains using multiple tools:

1. `RepeatClassifier` in the `RepeatModeler v2.0.2` suite (*51*) to find matches against sequences in the RepBase RepeatMasker Edition or Dfam DNA databases, respectively, using the `RMBLAST v2.11.0` search engine and default search thresholds.
2. `TEsorter v1.3` (*52*) to detect matches against TE protein domains using the REXdb-metazoa database (*53*).
3. `TransposonPSI v08222010` (http://transposonpsi.sourceforge.net/) with the `BLAST v2.2.26` search engine using the "PSI-BLAST" task (*54*, *55*) to query the LTRs against TE family profiles using the default search parameters.
4. `HMMER3 v3.3` (http://hmmer.org/) searches against Dfam LTR profiles. Here we first generated a list of profiles matching LTRs from the curated release Dfam_3.5 using the command:

    ```
    grep -P "(\tLong\sterminal\srepeat|LTR|Gypsy|Copia|retrotransposon)"
    ```

```
Dfam_curatedonly.hmm.list.csv | grep -v -P "(L\d+|LINE)" >
Dfam_curatedonly.hmm.list.csv.LTR.csv
```

We then used hmmfetch to extract the actual profiles and hmmsearch (“`--notextw --cpu 40 -E 1e-5 --domE 1e-5`”) to search a version of the LTRs where we had first masked out simple repeats with dust to reduce spurious hits driven by microsatellite motifs.

We next used a custom Perl script to parse the output from the four tools and only keep LTRs with a significant hit detected using at least one of them. The LTRs were classified according to the detected LTR type (superfamily or similar level) and tagged with the detection-tool in the following order of priority: i) RepeatClassifier (“ReC”); ii) TEsorter (“TEs”); iii) TransposonPSI (“PSI”); and iv) HMMER+Dfam (“DFA”).

LTR retrotransposons have two similar 5′ and 3′ LTR regions and we divided the LTRs into a pool of repeats with highly identical LTR regions (99–100% identities; putatively “young” motifs) and a pool of repeats with more divergent regions (85–99% identities; possibly older repeats). Sequences in each pool were then translated into peptides LTR_retriever’s `Six-frame_translate.pl` tool and then searched separately with hmmsearch (`--notextw --cpu 40 -E 0.01 --domE 0.01`) and the GyDB2 database (*56*) for complete Copia or Gypsy LTRs with all expected internal genes still present, i.e. those with simultaneous hits against all five typical protein domains: Capsid “GAG”, Aspartic proteinase “AP”, Integrase “INT”, Reverse transcriptase “RT” and RNase H “RH”. We then applied a step-wise procedure to reduce redundancy in the LTR libraries, while putting top priority on complete LTRs with high LTR region identities:

1. We combined the complete LTRs with 99% LTR region identities from LTR_Finder and LTRharvest into a single set and removed redundant 80/80-hits with CD-hit (`-c 0.80 -n 5 -M 0 -aS 0.80 -G 0 -g 1 -T 40`).
2. We then combined the complete LTRs with 85–99% identical LTR regions from LTR_Finder and LTRharvest and masked this set with RepeatMasker using the preceding non-redundant set from step 1. LTRs with <80% sequence masked were kept and clustered to remove redundancy with CD-hit as above. The representative sequences were then concatenated to the set from step one to create a non-redundant set of “complete” LTRs.
3. We next combined the incomplete LTRs with 99% LTR region identities and repeat-masked them with the “complete” LTR library, adding only <80% masked and CD-hit clustered non-redundant LTRs to the library. This was followed by a final iteration of masking and adding LTRs using the incomplete 85–99% LTR library as above.

Our final non-redundant and homology-based LTR library contained 365 full-length or near full length retrotransposon sequences (length N50=6,421bp) (table S5).

### Detection of diverse transposon domains using TransposonPSI

We carried out PSI-BLAST homology searches using the broad library of transposon ORF profiles (including both LTRs, LINEs, DNA transposons and other elements) in `TransposonPSI v08222010` (using `BLAST v2.2.26`) across the whole krill genome assembly. For efficiency, the genome assembly was first split in 100 chunks, and the chunks were scanned in parallel in TransposonPSI. High divergence between query and profile may result in fragmented HSP (High Scoring Pairs) hits interspersed by non-matching sequence. We compiled a preliminary repeat library from the "`*.TPSI.allHits.chains`" files, which contains chains of one or more collinear HSPs hits against transposon profiles. i.e. series of HSPs that align collinearly between a specific TE ORF and the genome, that are not interrupted by ORFs of other TEs. As in the LTR scans above, we then queried the library against the genome and kept only repeats with two or more 80/80-hits against the genome (n=190,408). Based on the initial classification assigned by TransposonPSI, we subdivided the repeats into broad groups: LTRs, LINEs, DNA transposons and Rolling-Circle transposons (Helitrons). LTRs were masked with the preliminary LTR library (see above) using RepeatMasker and only sequences with <80% masked positions were kept. The sequences of each group were then translated into peptides using Six-frame_translate.pl and either the DNA or peptide sequence was then used to re-classify the repeats using the programs and databases also used to classify the LTRs in the previous section:

1. hmmsearch was run with both the GyDB2 database to detect Copia or Gypsy domains and the REXdb-metazoa database to detect other TE domains (`--notextw --cpu 40 -E 0.01 --domE 0.01`). Putative LINEs, DNA transposons and helitrons were kept only if they did not have significant matches against LTR domains. The sets were concatenated and clustered with CD-HIT to remove redundant sequences (`-c 0.80 -n 5 -M 0 -aS 0.80 -G 0 -g 1 -d 0 -T 40`) before proceeding to the next step.
2. RepeatClassifier was run using both RepBase RepeatMasker Edition and Dfam DNA databases, as above.
3. TEsorter was run (`-st nucl -p 40 -tmp tmp -eval 0.01`) with the GyDB2, REXdb-metazoa and REXdb-tir dabases.

We used a custom Perl script to parse the output from the classification tools. Putative LINEs, DNA transposons and helitrons were kept only if they had at least one match against the expected group (class/order) and did not have significant matches against LTR domains. The repeats were classified according to the detected type (i.e. superfamily) and tagged with the detection-tool in the following order of priority: i) RepeatClassifier ("ReC"); ii) TEsorter ("TEs"); iii) TransposonPSI ("PSI", the original classification of the repeat). Compared to the refined structural LTR library, we detected more but shorter LTR fragments using this approach (table S5).

### Detection of transposable elements through *de novo* repeat assembly with dnaPipeTE

Repeats and TEs that occur at high frequency in the genome can be detected through *de novo* assembly of low-coverage short-reads. This approach is implemented in dnaPipeTE (*57*), which uses Trinity to assemble repeats from short-reads independently of a reference genome and

BLASTN to classify them against its own database. We extracted 16 independent batches of 10 million single short-reads from our 10x Chromium Illumina read libraries and assembled repeats independently for each batch using dnaPipeTE v1.3.1 (`-genome_size 19000000000 -genome_coverage 0.079 -sample_number 2`). We then parsed the "`one_RM_hit_per_Trinity_contigs`" output-file from each run with a custom Perl script and kept only those Trinity-assembled repeat contigs where a reported BLASTN match against a dnaPipeTE repeat template spanned >20% of length of both the contig and the template. The repeats of each run were concatenated into a single preliminary library (n=4,799) and subdivided into major groups (DNA transposons, LTRs, LINEs and SINEs). As in the preceding repeat scans, we queried each group against the genome and kept only repeats with two or more 80/80-hits against the genome (n=3,381). From manual inspection, we noticed that candidate TEs frequently consisted nearly exclusively of simple repeats / microsatellites. We therefore subjected also this preliminary library to reclassification with the same tools used to process the previous repeats:

1. The sequences of each group (DNA transposons, LTRs, LINEs and SINEs) were masked with RepeatMasker using the two LTR and TransposonPSI libraries from previous annotation steps, respectively, and only sequences with <80% bases masked were kept. Each group was clustered separately with CD-hit as above to reduce redundancy.
2. Each group was re-classified using the same tools and procedures outlined in the TransposonPSI pipeline above.

Our filtering and clustering resulted in 311 dnaPipeTE repeats being added to the growing repeat library, most of which appeared to be relatively short TE fragments (table S5).

#### Detection of transposable elements from high-frequency motifs using RepeatModeler2

We ran RepeatModeler2 (*51*) across a random subset of contigs corresponding to 10% of the krill genome assembly and detected n=3,799 putative consensus repeat sequences occurring at high copy numbers. We applied a similar approach as in preceding steps to remove redundant repeats from this preliminary library:

1. The sequences were masked with RepeatMasker using the three finalized LTR, TransposonPSI and dnaPipeTE libraries, respectively, and kept only RepeatModeler sequences with <50% bases masked. These were then clustered with CD-hit as above to reduce redundancy, generating n=2,091 repeat sequences.
2. The sequences were re-classified using RepeatClassifier as above. Only 156 could be annotated with RepeatClassifier, suggesting that RepeatModeler2 tended to detect high-frequency repeats with low homology to known transposable elements motifs in the repeat databases, possibly due to excess uncorrected sequence error or evolutionary degeneration or divergence among repeat copies detected throughout the genome.
3. To improve the annotation rate of this library, we scanned individual repeat copies detected by each consensus sequence. First, we mapped consensus sequences to the

genome assembly with MEGABLAST and kept up to n=1,000 hits against the genome with >90% identity across 80% of the consensus sequence. We then queried each copy against a curated protein database in RepeatMasker using its utility RepeatProteinMask (which uses BLASTX), registering alignment scores with p-values <0.01. We summed the alignment scores across copies and annotated the consensus sequence with the order/superfamily classification of the RepeatProteinMask template with highest total score (i.e. the majority vote). We required that the majority vote was based on at least 10 observations and otherwise left consensus sequences unannotated. This approach added annotations to 644 consensus repeats, more than double the annotation rate compared to annotating the consensus sequences directly with RepeatProteinMask (n=254).

Our RepeatModeler library thus spanned 2,091 interspersed repeat sequences, out of which 800 were annotated to class and subfamily (table S5).

#### Evaluation and validation of the repeat library

We concatenated all libraries with interspersed repeats into a single library (n=10,908; 13.8Mb). As a means to evaluate the annotation and characteristics of the library with independent information, we queried the library against a set of 1,000 common transposon-associated protein domains derived from the NCBI Conserved Domain Database (CDD). The domain data was available in the TransposonProteinNCBICDD1000 tool in the TransposonUltimate annotation suite (*58*). We used the "Reverse Position-Specific BLAST" RPSBLASTN tool to query the library and
detect significant hits against pre-calculated Position-Specific Score Matrices (PSSMs) of these domains (*59*, *60*), allowing for low-identity hits with e-values <1:

```
rpstblastn -query <library.fa> -db
proteinNCBICDD1000/selection/Selection1000Library -outfmt "7 stitle evalue
qstart qend" -evalue 1 -outlibrary.fa.txt
```

#### Annotation of repeats across the krill genome with RepeatMasker

We split the genome assembly into 100 chunks and used `RepeatMasker v4.1.1` to annotate each chunk in parallel in the high-performance Uppmax computing environment:

```
RepeatMasker -pa 14 -norna -e rmblast  -a -u -gff -xsmall -gccalc -lib
<repeat_library.fa>"
```

We used the RepeatMasker tools `calcDivergenceFromAlign.pl` to calculate the Kimura 2-Parameter divergence metrics from repeat alignments and `createRepeatLandscape.pl` to generate an overall repeat landscape distribution. We also used a custom script to parse the RepeatMasker .out file and estimate the amount of every TE superfamily (or similar).

Statistics about the repeat library and repeat annotation is available in table S5.

#### *Gene annotation*

We used both RNA data and comparative data from other genome-sequenced crustaceans to annotate genes in the krill genome, with the ultimate aim to produce a non-redundant set of protein-coding genes that could be used for genome and SNP annotation.

##### RNA-based data

We mapped 755 M Illumina short-read RNA-seq read-pairs derived from the reference specimen itself and two published specimens (one male; one female) (*19*) against the assembly with `HISAT2 v2.2.1` (*34*) (mapping rate 88.2%). We then mapped 8.7 M full-length PromethION cDNA reads produced from the reference specimen against the assembly using minimap2 v2.17-r974 (mapping rate 95.8%). `StringTie v2.2.0` (*61*) was used in "mixed" mode to generate reference-guided RNA transcript models in GTF-format from the short-read and long-read mappings simultaneously.

Trinity transcripts that had been assembled independently from the genome assembly (see above) were used as a second line of RNA-based evidence of genes. Only putatively coding transcripts with long open reading frames and domain hits against Metazoan templates (n=60,677) were carried forward in this analysis. `SPALN v2.4.6` (*62*) was used to build a DNA database of the assembly (-KD) and to produce spliced alignments of the Trinity transcripts in GFF3 format ("`-Q7 -LS -d<genome database> -O0,1,3`").

##### Comparative data

We aligned peptide sequences from the KrillDB reference transcriptome from the Antarctic krill *Euphausia superba* (*63*) and published gene models from six genome-sequenced crustaceans (n=304,549; table S6) against the genome assembly using SPALN, with the ambition to potentially recover additional gene models not well-represented by our RNA data. A protein database was built for the assembly with SPALN (-KD) and spliced protein alignments were then produced in GFF3 format ("`-Q7 -LS -d<genome database> -O0,1,2,3,4`"). For downstream compatibility, a custom Perl script was used to ensure that start coordinates for all features were always smaller than stop coordinates, regardless of strand orientation. `GFFCOMPARE v0.12.6` (*64*) was used to combine the protein-based GFFs and assign the alignments to a set of common loci with different isoforms. We used `TransDecoder v5.5.0` to only keep candidate alignments that retained significant protein domain homology by: i) converting GTF to GFF3 (`gtf_to_alignment_gff3.pl`); ii) generating matching sequences in FASTA format (`gtf_genome_to_cdna_fasta.pl`); and iii) search for long open reading frames (`TransDecoder.LongOrfs`) in strand specific mode ("`-S`").

The candidate ORFs were searched for homology against known proteins in the Swissprot database (timestamp 2021-06-16) with `BLASTP v2.9.0+` (*33*) and the Pfam database (release 34.0) using `HMMER3 hmmscan v3.3` (http://hmmer.org/), respectively, according to standard procedure (https://github.com/TransDecoder/TransDecoder/wiki). `TransDecoder.Predict` was used to identify the peptide alignments with significant hits, `cdna_alignment_orf_to_genome_orf.pl` to write a GFF3 file with the corresponding genome coordinates and GFFREAD (*64*) to convert in into GTF format. To reduce the risk of carrying forward low-quality protein alignments and with the aim to generate a non-redundant set of gene models, we used custom Perl scripts to parse the GFFCOMPARE locus *.tracking file in order to: i) discard loci labeled by only one crustacean species; and ii) to select one representative alignment from each locus with the highest Swissprot BLASTP score from the preceding step; and iii) generate an updated GTF file.

#### Consolidation of gene models

We combined the three sources of potential gene models generated above (*i*=StringTie RNA-seq transcripts; *ii*=SPALN-aligned Trinity transcripts; *iii*=non-redundant comparative crustacean models) with GFFCOMPARE and predicted coding transcripts with Transdecoder (as in the previous section). We next queried all predicted peptide sequences against the full invertebrate NCBI RefSeq database (timestamp 2021-03-05) using `DIAMOND v0.9.9` (*65*). From each gene locus identified with GFFCOMPARE, we selected the gene model/isoform with the maximum DIAMOND alignment score to a RefSeq sequence, in order to produce a set of high-quality and non-redundant gene models across the krill genome assembly. The gene labels of both the non-redundant and redundant gene sets were tagged to reflect the source of the evidence, including the best model of each locus.

#### Gene set size and completeness

Altogether, 202,138 models were generated from combining the datasets as above, out of which 118,528 models were found to be putatively protein-coding with TransDecoder. After removing redundancy by selecting the single best DIAMOND+RefSeq model or isoform, 42,227 non-redundant gene models were retained. Of these, 30,766 (73%) models were selected on the basis of a highest-scoring reference specimen StringTie model (assigned "REF_STRG" tag in the sequence header), 8,063 (19%) models from a reference specimen Trinity transcript ("REF_TRIN" tag) and 3,398 (8%) models from comparative peptide data from other crustaceans. TransDecoder annotated genes as "*complete*", "*3prime_partial*" (i.e. missing terminal exons or stop codons), "*5prime_partial*" (missing start codon) or "*internal*" (missing both start and stop codons). 26,448 models were annotated as "*complete*" or "*3prime_partial*" (63%), 7,379 were annotated as "*5prime_partial*" and 8,400 annotated as "*internal*" (table S7). We used a custom Perl script to detect missing in-frame stop codons directly downstream of models annotated as "*3prime_partial*" or "*internal*", restoring stop codons in 5,884 gene models, primarily derived from transcript or peptide data (table S7).

We then used both `BUSCO v3.0.2b` (*16*) with the Arthropoda odb9 lineage set and `BUSCO v5.0.0` (*22*) with the Arthropoda odb10 lineage set, respectively, to assess the completeness of the gene models (Fig 1B, fig. S4).

#### Functional and evolutionary annotations

We applied several tools to accomplish functional annotations of the gene sets. First, we used `EnTAP v0.10.7` (*66*) with the SwissProt and invertebrate RefSeq peptide databases referred to above. To this base resource, we added gene models from multiple genome-sequenced arthropod species not represented in our release of RefSeq (table S8). We ran EnTAP in protein-mode (“`--runP`”) with default settings, using the TransDecoder-predicted peptide sequences derived from the krill gene models and EnTAP’s pre-configured EggNOG framework resources (*67*) to annotate genes. This resulted in 27,584 of the 42,227 gene models being annotated, while 14,643 remained unannotated (table S7). Second, we downloaded the *Drosophila* FlyBase database release FB2021_01 (*68*) and used `BLASTP v2.9.0+` (*33*) to find *Drosophila melanogaster* (dmel_r6.38; n=30,724 peptides) homologs of our krill peptides using. For each krill peptide sequence, the best scoring *Drosophila* homologue was retained as a gene label for downstream annotation of genes or SNPs without further resolving best reciprocal relationships. The “FBgn” Flybase IDs were used to compute Gene Ontology enrichments for gene family expansions and for divergence signals between populations (see sections below).

### Gene region masking

The non-redundant set of protein coding gene was used to generate a mask across the whole genome with a custom Perl script, in which every base was substituted for the type of gene region it occurred in (1=intergenic; 2=intron; 3=3′-UTR; 4=exon; 5=5′-UTR; 6=cds including start/stop codons). This was used to enumerate the lengths of different gene regions and to classify specific positions (e.g. in the subsequent DNA methylation analyses or assessments of variation across the genome).

We then estimated repeat-content for each gene to test for correlations with annotations (as a control) and expose potential transposon models among unannotated gene models. We therefore used the krill repeat library to mask the coding sequence (CDS) of all gene models with RepeatMasker (using the same settings as the whole-genome masking). We aimed to measure and contrast the repeat content between “regular” gene models that had been annotated for function and homology towards other invertebrate genes using EnTAP against those matching transposable elements in other species or those that were unannotated. To distinguish between regular gene annotations and transposable element annotations, we applied keyword searches against: i) each (per-gene) line in the EnTAP output; and ii) the descriptions matching the COG (Clusters of Orthologous Genes) and eggNOG tags reported by EnTAP (field number 31 “EggNOG Member OGs” in the EnTAP summary table). To match a putative transposon annotation, either the whole line itself or the COG/eggNOG ortho-group descriptions had to:

i) have a match to either of the keywords: “retrotrans” or “transposon” or “transposable” or “transposase or “reverse transcriptase” or “RVT_1”;

ii) but not a match to the keywords: “retrotranslocation” or “piRNA” or “miRNA” or “prevent” or “repress”, since these keywords were associated with genes responsible for inhibiting transposable element activities.

This resulted in the identification of 25,301 regular protein-coding gene models and 2,283 transposable element annotations and those models that were unannotated in EnTAP (n=14,643). Their respective lengths, repeat content and other features are provided in table S7 and fig. S6.

#### Evolutionary analyses of protein coding genes

We aimed to assess gene and gene family evolution in the krill by comparing the set of coding genes in our annotation to orthologs of nine other crustaceans (table S9). We prepared the data by removing redundant gene sequences from published gene models of the other species, keeping only the isoform with the longest peptide sequence for every gene.

##### *Inference and comparison of orthologs between krill and other species*

We performed a search for orthologs using `ProteinOrtho v6.0.14` (*69*) with DIAMOND v0.9.30. We used the 26,448 non-redundant Northern krill gene models that had been annotated as complete in this analysis. We then enumerated all 1:1 single-copy ortholog pairs between the krill and the other species. We used 1:1 orthologs between the krill and those of *H. americanus*, *P. monodon*, *H. azteca* and *E. affinis*, respectively, for direct comparative analysis of the lengths of coding sequence and full gene bodies in R (*70*). Gene length comparisons are provided in Fig 1., fig S7 and table S10. For the Antarctic krill, comparisons were done against the mean length reported in the Antarctic krill genome paper (*71*) (no gene models were available at the time of preparing these analyses).

##### *Estimating divergence between the Northern krill and the Antarctic krill*

We estimated per-base divergence at synonymous sites (*dS*) between the Northern krill and the Antarctic krill as a proxy for nearly neutral divergence comparably unaffected by selection. We then used this estimate to date when the two species split from their most recent common ancestor.

To this end, we first used DIAMOND to detect 13,371 pairwise Reciprocal Best Hits between 140 K Antarctic krill transcriptome protein models from the KrillDB2 and 26,395 non-redundant Northern krill protein coding gene models (see section below about the gene models). RBH protein sequences were aligned with `MAFFT v7.453` (*72*) using settings recommended to avoid over-aligning regions with poor homology (*73*):

```
mafft --globalpair --allowshift --unalignlevel 0.8 --leavegappyregion --
anysymbol seq.aa.fasta > seq.aa.fasta.ginsi.fasta
```

For each alignment, we fitted the corresponding nucleotide sequences with `PAL2NAL v14` (*74*) and then used `KaKs_Calculator v1.2` (*75*) with the approximate “YN” method to estimate synonymous and non-synonymous sites and substitutions and compute *dN*, *dS* and *dN/dS*, while accounting for unequal base frequencies, transition/transversion rates and multiple substitutions between sequences (*76*). We summed the total synonymous sites and substitutions to derive a single overall *dS* estimate across all 13,371 genes.

In addition, we concatenated the genes and computed the observed and Jukes-Cantor-corrected distances between the two species across full sequences or 1st, 2nd and 3rd positions separately using Seaview v5.0.5 (*77*).

Using the *dS* estimate of 0.46, we next estimated divergence time assuming the mutation rate inferred from snapping shrimp in (*49*) of $2.64e^{-9}$ substitutions per site per generation and a 1.5 year generation time (1 year in Northern krill and 2 years in the Antarctic krill) (*78*, *79*). We used a standard linear equation assuming a constant molecular clock:

$$T = \frac{D * g}{2u}$$

Here, *D*=divergence per base (*dS*=0.46), *g*=generation time (1.5), *u*=mutation rate ($2.64e^{-9}$).

Results of these calculations are provided in table S10.

#### Inferences of phylogenetic interrelationships

We detected 1,011 single-copy orthologs (OGs) across all ten species and used these genes to produce a time-calibrated phylogenomic crustacean species tree. We first aligned the peptide sequences of each OG with MAFFT and filtered the alignments with `Gblocks` (*80*):

```
Gblocks <OG.fa> -t=p -b1=6 -b2=6 -b4=5 -b5=h -b6=y -v=10000 -d=y -e=.gb
```

We then inferred phylogenetic interrelationships with `IQ-TREE v2.1.0` (*81*) , applying the best-fit LG+F+I+G4 model (Modelfinder+BIC) across all filtered ortholog alignments as a supermatrix and running 1,000 ultrafast bootstrap replicates (*82*). The crustacean phylogeny was converted into an ultrametric chronogram using `r8s v1.8.1` (*83*). Input for r8s was generated using a helper script from the `CAFE v5.0` (*84*), where we specified that the phylogeny had been inferred from 522,746 alignment sites. We used three calibration points, setting: i) the oldest split in the tree (between *D. magna* and all others) to have occurred 530 M years ago (*85*); ii) the split between the lobster *H. americanus* and the crayfish *C. quadricarinatus* to be at least 372 M years old (i.e. older than the radiation of lobsters) and the split between krill and decapods to be at least 447 M years (i.e. older than the radiation of decapods) according to molecular dating in (*86*). We ran r8s using the penalized likelihood method:

```
divtime method=pl algorithm=tn cvStart=0 cvInc=0.5 cvNum=8 crossv=yes
```

#### Analysis of gene family evolution

We used `SwiftOrtho` (https://github.com/Rinoahu/SwiftOrtho) (*87*) to broadly cluster genes into putative gene families and used CAFE to trace the expansion or contraction of these families along the time calibrated ultrametric tree inferred in the previous step. We then used `wgd` (*88*) to test for signatures of divergence between genes that could indicate a history of whole genome duplication.

### Preparing the krill gene set

The full krill gene set spanned n=42,227 putative gene models, out of which n=27,584 genes had been annotated with EnTAP and 18,962 genes had been tagged as complete using TransDecoder (table S7). Because not all gene models were recognised for function or homology with EnTAP or complete with TransDecoder, the full gene set could contain spurious gene models (e.g. non-coding RNA, pseudogenes or transposable elements) or incomplete gene models (e.g. fragments of the same gene locus separated on unscaffolded contigs) that could artificially inflate gene numbers and estimation of gene family expansions.

From the set of annotated genes, we therefore first removed 2,283 potential transposable elements (TEs) that had been annotated as TEs using EnTAP, leaving 25,301 models in this set. We then performed a relaxed homology search between unannotated krill genes and the nine crustaceans (table S9) with DIAMOND using an e-value threshold of $1e^{-5}$ and detected significant hits for 1,934 hitherto unannotated genes. These genes were added to the annotated set for a total of 27,235 genes. This set was then checked for potentially fragmented models that may belong to the same underlying gene locus through queries against two contiguous sets of RNA sequences:

a) We mapped the gene sequences back to the Trinity transcripts (using the longest isoform of each putative Trinity gene) to identify genes that mapped to different parts of the transcripts. For each gene, the RNA transcript with the longest total BLAST hits was registered. If gene models mapped to the same transcript and their respective alignments did not overlap at all or only overlapped marginally (at most 10% of the length of the gene), they were taken potential separate parts of the same gene body in the genome. In these cases, the shorter and potentially redundant gene fragment(s) were removed from the set, so that only one representative gene model was kept.
b) We also mapped the gene models to the full length Nanopore cDNA sequences that had previously been clustered with VSEARCH (see above; n=25,484), and detected short and potentially redundant fragments of the same underlying gene using the same approach.

This procedure identified and removed 839 gene fragments from the gene set, the majority of which had been marked as incomplete by TransDecoder, resulting in a final krill gene set spanning 26,395 non-redundant gene models that were carried forward for gene family analysis.

### Analyses with SwiftOrtho to cluster genes into families

SwiftOrtho inference of gene families was performed in three steps for each dataset:

i) find similar matches between sequences in the non-redundant crustacean gene sets:

```
find_hit.py -p blastp -i <species.fa> -d all_proteins.fa -a 20 -e 1e-5 -s
111111 -o <species.fa.out>
```

ii) infer potential orthology relationships between genes using low taxon coverage ("-c 0.3") and identity thresholds ("-y 30"):

```
find_orth.py -i <all_matches.out> -a 16 -c 0.3 -y 30 > <all_matches.out.orth>
```

iii) cluster similar orthologs or paralogs into groups/gene families using the Affinity Propagation Cluster (APC) algorithm:

```
find_cluster.py -i <all_matches.out.orth> -t 16 -a apc -I 1.5 >
<all_matches.out.30_30.orth.apc>
```

The clustering resulted in a gene count table spanning 17,119 gene families with sequences from two or more species. This set included 11,259 families with at least one gene from the krill and 6,141 families where both the krill and the outgroup *D. magna* were represented. For comparison, the lobster *H. americanus* and amphipod *H. azteca* were represented in 12,274 and 9,149 gene families, respectively.

### Analyses with CAFE to trace gene family expansions or contractions

The SwiftOrtho gene family dataset was reformatted into the expected input for CAFE using a custom Perl script and analyzed in CAFE5. The software uses a maximum-likelihood framework to estimate the rates and events of gene family evolution using a birth-death process to model the gain or loss of genes along the branches of the species tree (*84*). These analyses are conditioned on there being at least one gene present in the outgroup taxon for a gene family to be included. Our analysis of gene family evolution was therefore restricted to families with at least one representative in the *D. magna* (n=6,814).

Here we followed the steps outlined by the developers in https://github.com/hahnlab/CAFE5/blob/master/docs/tutorial/tutorial.md:

1. Separating extremely large gene families from the rest to reduce variance in gene copy numbers and improve parameter estimation:

   ```
   python clade_and_size_filter.py -i
   <all_matches.out.30_30.orth.apc.table.ALL.csv> -o
   all_matches.out.30_30.orth.apc.table.ALL.csv.filtered -s
   ```

   In total, 16 large families were separated from the rest in the first size filter step, all of which indicated expansion in other species than the krill. For this reason, those 16 families were not further analyzed.

2. Infer the gene family evolutionary rates (the so-called lambda parameter) and compute gene family evolution across the species tree.

   ```
   cafe5 -i all_matches.out.30_30.orth.apc.table.ALL.csv.filtered -t
   ```

`filtered.iqtree.out.contree.rooted.nwk.r8s_ctl.txt.out.tre`

We compiled distributions of gene family sizes for each species, estimated the optimal lambda across the tree to be 0.0005932 and inferred the patterns of expansion/contractions across the 6,814 gene families. Results are provided in Fig. 1, fig. S7 and table S11.

### Gene ontology (GO) enrichment analysis of expanded gene families

The subset of gene families with significant evidence of expansion in the Northern krill was analyzed for function by assessing the corresponding *Drosophila melanogaster* homologs using ontologies in the online Flybase service. Using the same *Drosophila* homologs, we tested for enrichment of Gene Ontology terms among the expanded families vs the non-expanded ones using the online services `GOrilla` (*89*) and `ShinyGO` (*90*) using False Discovery Rates (FDR) thresholds of 0.05. Analyses of gene families were carried in out in two ways with regards to the *Drosophila* homologs: i) using all homologs associated with a gene family (which may be highly variable in the number of homologs between gene families); ii) using only the most commonly detected homolog for every gene family (only using more than one in case of ties; an approach to reduce variance in the numbers of homologs used to characterize each gene family).

The results of these analyses are presented in Fig. 1 and table S11.

### Analyses with wgd to test for signatures of whole-genome duplication (WGD)

The amount of divergence at neutral sites between two homologous sequences can serve as a proxy for the time that has passed since they originated through duplication. In the case of genes, such divergence can be measured as synonymous substitutions per synonymous site in pairwise alignments between two sequences ($K_S$), assuming that such sites evolve neutrally. Genes can be added to the genome through small-scale duplication events at different points in time or through large-scale duplications including whole-genome duplications, in which many genes originate at the same time. In each scenario, gene duplicates are often lost after some time. The former process can be expected to produce an “L-shaped” exponential $K_S$-distribution among surviving paralogues in the “paranome”, suggesting many observed paralogues are young, while the latter process can produce one or more characteristic secondary peaks along the $K_S$-distribution, indicating that those paralogues originated at a particular event and age (*91*). To study these patterns, we analyzed the gene set of the krill and, for comparison, those of five other crustaceans that have not been associated with whole-genome duplications (*H. americanus*; *P. monodon*; *H. azteca*; *E. affinis*; *D. magna*) with `wgd` (*88*). The tool internally uses DIAMOND to detect similarity among sequences, MCL (*92*) to cluster them into putative gene families, FastTree to produce per-family gene trees (*93*), PAML (*94*) to perform model-based estimation of synonymous divergence between paralogs and Scikit-learn (*95*) to fit mixtures of components to the overall $K_S$-distribution in order to find evidence for mixed distributions that could indicate WGD. For each species, we:

1. Performed clustering of the genes:

```
singularity exec wgd.sif wgd dmd --nostrictcds <cds.fasta>
```

2. Computed and visualized the respective $K_S$-distribution:

```
singularity exec wgd.sif wgd ksd -n 16 --preserve --max_pairwise 250000
wgd_dmd/<cds.fasta.mcl> <cds.fasta>

singularity exec wgd.sif wgd viz -ht barstacked -r 0 5 -b 50 --weighted
-ks <cds.fasta.ks.tsv> -o <cds.fasta.ks.tsv.viz.svg>
```

For the krill, we then performed additional analyses using Gaussian Mixture Models and applying models with 1 to 4 components and applying the Bayesian Information Criterion to test which model best explained the data:

```
singularity exec wgd.sif wgd mix --method gmm -n 1 4 -o wgd_mix_gmm
<cds.fasta.ks.tsv>
```

The results of these analyses are presented in fig. S8 and table S11.

#### *Characterization of the Hox gene complement*

The presence and number of duplicate Hox genes can indicate ancestral whole genome duplication in a lineage, as these genes are often retained in duplicate copies even after diploidization (*96*). To test for duplicated Hox genes in the krill, we used curated Hox homeodomain sequences from a previous analysis (*97*), HomeoDB (*98*) or NCBI. We searched the krill genes against 10 core homeodomain sets from crustaceans and selected insects, spanning 60–80 amino acids and 3–6 species per set, to annotate Hox genes and detect Hox gene duplication. Eight well-curated homeodomain sets were used unchanged from analyses of the amphipod *P. hawaiensis* (*97*) (*lab=labial*, *pb=proboscipedia*; *Hox3*; *Dfd=Deformed*; *Scr=Sex combs reduced*; *Antp=Antennapedia*; *adbA=abdominal-A*; *AbdB=Abdominal-B*), one set was downloaded from HomeoDB (*98*) (*Ftz=Fushi Tarazu*; HomeoDB species: *Drosophila melanogaster* [Dm]; *Apis mellifera* [Am]; *Tribolium castaneum* [Tc]) and the final set was derived from both HomeoDB and NCBI sequences (*Ubx/Utx=Ultrabithorax*; HomeoDB species: *Drosophila melanogaster* [Dm]; *Apis mellifera* [Am]; *Tribolium castaneum* [Tc]; NCBI species: *P. hawaiensis* [Ph] FJ628448; *D. magna* [Dmag] BAE96992.1).

We queried the krill gene against the 10 homeodomain sets using BLASTP. We kept the longest matching peptide substring (the putative homeodomain) for sequences that had at least one hit to a homeodomain template which spanned ≥50 amino acids with ≥50% identity (n=119 sequences). Because the homeodomain is relatively similar across the different genes, we then combined all 10 sets and krill substrings into a single FASTA sequence file and aligned all data at once with MAFFT. We then inferred a homeodomain gene family tree with Fasttree v2.1.11 (*93*) and inspected the phylogenetic positions of the candidate krill genes.

The Hox gene family tree is shown in fig. S11.

#### *Characterization of genes involved in DNA methylation*

The presence or absence of genes that encode enzymes involved in DNA methylation (i.e. DNMT1 or DNMT3), DNA demethylation or associated damage repair (e.g. TET family; ALKB2) vary among arthropods and may indicate whether the DNA methylation pathway is operational (*99*). Since we had detected DNA evidence of genome-wide DNA methylation through Nanopore signal analysis (see below), we searched the krill gene-set for homologs encoding the DNMT1/2/3, TET2 and ALKB2 enzymes. We first identified potential candidates using the gene names of best-matching genes from the EnTAP gene annotation. We then combined arthropod sequences from multiple sources with established or likely orthology, including OrthoDB v10.1 (*100*), EggNOG v5.0 (*101*) and NCBI (mostly crustacean sequences) (table S12) and performed phylogenetic analyses of the position the candidate krill genes in each gene tree. For each gene, we aligned the peptide sequences with MAFFT, trimmed poorly aligned regions with `trimAl v1.2rev59` (*102*), inferred a gene tree with FastTree and inspected the position and the length of the branch at which the krill sequence(s) grouped in the tree. For further validation, we performed analyses using the online NCBI Conserved Domain Search (*59*, *60*) and SWISS-MODEL (*103*) for homology-based inference of structure and domains in each candidate sequence. SWISS-MODEL uses a comprehensive search strategy to detect significant matches, including the composite Qualitative Model Energy ANalysis (QMEAN) scoring method to analyze and score protein structures.

#### Predicting protein structures

To further assess the accuracy of these candidate proteins possibly involved in the DNA methylation system, we predicted their three-dimensional protein structure. The protein sequences were added to the ColabFold server (*104*), where the input multiple sequence alignment is built by a homology search by `MMseq2` (*105*) before modeling by `AlphaFold2` (*106*). No PDB templates were added, and 3 recycles were used. Five models were generated per protein.

#### **Analysis of DNA methylation levels**

Nanopore signals are altered by DNA base-level modifications, including 5-methylcytosine (5-mC) at CpG sites, that can be detected *in silico* using Nanopore data (*107*). We scanned the Nanopore reads for evidence of methylated cytosines across all well-covered CpG sites in the nuclear and mitochondrial genomes (i.e. DNA methylation in the somatic muscle tissue that the sequences were derived from), and cross-referenced methylation frequencies with genomic features.

We first mapped the PromethION long-reads back to the finished krill genome assembly using minimap2 as above and then used the GPU-accelerated program `f5c v0.6` (*108*) to scan for evidence of methylated cytosines across all CpG sites in the genome with 10× or higher read coverage. We used three f5c subcommands:

1. "`index`" to index the raw FAST5 PromethION signal data and reads;
2. "`call-methylation`" to call the likelihoods for CpGs to be methylated or not;
3. "`meth-freq`" to compute the frequency of reads with sufficient evidence of methylation at every CpG.

We then used a custom Perl script to group the reported methylation frequencies according to genomic regions using a genome mask (see above). Depending on how closely spaced CpG sites are, f5c sometimes outputs joint methylation frequencies across a short subsequence with more than one CpG site. In such cases, we split the observation into separate data points with identical methylation frequencies and queried each position against its corresponding genomic region. We then compiled a second round of summary statistics removing potential source of error by: i) masking all CpG sites at which we had observed heterozygous genotypes in the reference specimen; ii) excluding all regions that were marked inaccessible due to high or low depths in the population genomic dataset (see below). We visualized the results in R and computed mean, median and 95% confidence intervals (200 bootstrap replicates).

We cross-referenced the DNA methylation data with information about reads and repeats. First, for every gene model we counted the number of RNA splice isoforms inferred from RNA-seq data and StringTie (see above), keeping genes supported by at least one isoform (n=21,031). We partitioned genes into 5%-bins of average DNA-methylation level and computed the mean number of isoforms in each bin, along with 95% confidence intervals (1,000 bootstrap replicates). We then extracted 1,706 putative LTR retrotransposons originally reported by LTR_Retriever and verified to contain at least one expected LTR domain when queried against the GyDB2 database (as above). Identities between 5'-LTR and 3'-LTR regions estimated by LTR_Retriever were used to partition the LTRs in bins of 5% divergence (here 0 to low divergence between LTR regions is taken as a proxy for being an evolutionarily young and recently inserted repeat).

Results are summarized in Fig. 1 and fig. S17.

**Population genetic data processing and analyses**

#### *Processing re-sequencing data*

The population-scale data was mapped to the genome, read-group tagged and marked for optical duplicates using standard NGS stools. We made a per-base depth profile across the genome to mark positions beyond ±50% of an observed putative diploid peak (fig. S18A) as inaccessible for measuring variation. SNPs and short structural variants were then called, decomposed and quality-filtered for annotation and analyses.

#### Mapping, read-group tagging and duplicate marking datasets

The barcoded short-read re-sequencing data was demultiplexed by NGI, resulting in one or more paired FASTQ archives per specimen (one pair per lane). The genome assembly was indexed using BWA as above. For each sample, we mapped and readgroup-tagged the data using BWA v0.7.17 with the “mem” algorithm and sorted the output on the fly using samtools v1.12:

```
bwa mem -t 18 -R <read_group> <reference> <fwd.fastq.gz> <rev.fastq.gz> |
samtools sort -@ 4 -O bam -m 4G -o <mapped_sample>.bwa.bam -
```

BAM files for samples sequenced on multiple lanes were merged with samtools. The merged data of each sample was then sorted by read name, marked for duplicates using samblaster v0.1.24 (*109*) and again re-sorted by genome coordinates:

```
samtools sort -@ 10 -n -O SAM <mapped_sample.bam> | samblaster | samtools sort
-@ 10 -O BAM -o <mapped_marked_sorted.bam>
```

For the reference specimen, both the high-coverage linked-reads and low-coverage data were used. We mapped the high-coverage linked-reads and processed it in the same way. All BAMs were indexed with samtools.

#### Mapping-depth profiles and masking of inaccessible sites

Our analyses of repeats had indicated that the krill genome assembly was highly repeated (see above) and that short-read mappings tended to result in uneven coverage across the genome, suggesting that many genomic regions could be problematic for SNP-calling. We therefore produced depth of coverage profiles across the genome with the aim to mark regions deemed to be accessible or inaccessible for SNP discovery. To this end, we compiled a per-base depth-track across the whole genome for each specimen, counting reads that mapped with a minimum quality of 10:

```
samtools view -q 10 -b <sample.bam> -b `cat contig.list` | samtools depth -a
--reference <genome_assembly.fa>
```

Using custom Perl scripts, we merged these depth tracks into per-population tracks, as well as one overall depth-track for the whole dataset. We compiled a depth distribution across the whole genome and identified a peak of coverage at ~188× and set lower and upper thresholds to be ±50% (94×) around this peak (fig. S16). Positions where less than 50% of samples were mapped (n<37) were masked as inaccessible. Downstream, we only considered SNPs and estimated patterns of genetic variation from genomic regions with 94–281× depths of coverage (8.43 Gbp) and for which data had been mapped in at least 50% of samples (n=37). We produced a per-base genome mask marking sites as either accessible (“1”) or inaccessible (“0”) and used this mask to correct estimates of the levels of genetic variation in the genome. The accessibility mask was cross-referenced against the mask of genome regions (see above).

### *Calling and phasing single-nucleotide polymorphisms (SNPs) across the genome*

#### Calling and processing variants

We subdivided the genome sequences (scaffolds and unscaffolded contigs) into 160 approximately equally sized chunks specified GTF files to enable SNP-calling different parts of the genome in parallel. We then used `FreeBayes v1.3.4` (*110*) to call variants with a theta prior set to “`-T 0.01`”, “`--use-best-n-alleles 5`” to limit memory usage, “`-m 10`” to limit analysis to read alignments with mapping scores above 10 (matching depth estimates in a previous step), “`-q 20`” as a minimum base quality filter for alleles and “`-E 0`” to limit the generation of complex haplotype variants. We also used inclusive depth filters outside mapping depth thresholds: “`--min-coverage 80`” as a lower threshold and “`-g 474`” as an upper threshold and piped the output to the Vcflib v1.0.1 (*111*) vcffilter tool set to only keep variants with QUAL > 20:

```
freebayes -T 0.01 -E 0 -m 10 -q 20 --min-coverage 80 -g 474 --use-best-n-
alleles 5 -t <genome_part_${NUM}.gtf> -f <reference.fasta> <bam files> |
vcffilter -f \"QUAL > 20\" > <variants_${NUM}.vcf>
```

We processed the called variants to decompose complex haplotypes and only keep biallelic SNPs. First, we used `bcftools v1.12` (*26*) with the “`norm -d all -a`” parameters to left-align and normalize indels, decompose complex variants and only keep one record per position. The output was piped to `vt v0.5772` (*112*) to split multiallelic variants while correcting for read counts (“-s”):

```
cat variants_${NUM}.vcf | bcftools norm -d all -a -f <reference.fasta> | vt
decompose -s - > <variants_${NUM}.decomposed.vcf>
```

We then used two custom Perl scripts to:

i) remove variants that fell outside accessible regions

```
vcf2filtered_vcf_by_coverage.pl --vcf <variants_${NUM}.decomposed.vcf> --
coverage <depth_mask.fasta> --seqs <genome_part_${NUM}.gtf> --verbose
```

ii) keep only biallelic SNPs in which at least 50% of samples had been genotyped and while also ensuring that these SNPs occurred within the desired thresholds:

```
vcf_biallelic2fasta.pl --input variants_${NUM}.decomposed.accessible.vcf --
output variants_${NUM}.decomposed.accessible.finished.vcf --min_fill_position
0.5 --min_depth 94 --max_depth 281
```

#### Imputing and phasing SNPs

Each chunk of SNPs (n=160) was imputed and phased with `BEAGLE v4.0` r1399 (*113*) in a two-step process. We first imputed missing genotypes using genotype likelihoods:

```
java -Xmx48g -jar beagle.27Jan18.7e1.jar nthreads=10 gl=variants_$
{NUM}.decomposed.accessible.finished.vcf out=variants_$
{NUM}.decomposed.accessible.finished.imputed.vcf
```

We then phased the data using the inferred genotypes:

```
java -Xmx48g -jar beagle.27Jan18.7e1.jar nthreads=10 gt=variants_$
{NUM}.decomposed.accessible.finished.imputed.vcf.gz out=variants_$
{NUM}.decomposed.accessible.finished.imputed.phased.vcf.gz
```

This resulted in a dataset spanning ~760 million biallelic SNPs.

#### Annotating SNPs

We used GFFREAD to extract the CDS DNA sequences of the non-redundant protein-coding genes (see gene annotation above):

```
gffread -g <genome.fasta> -x <cds.fasta> <genes.gff>
```

The GFF gene coordinate file and genome and cds sequences were then used to build a custom database with `SnpEff 5.0e` (*114*):

```
java -Xmx96G -jar snpEff.jar build -gff3 -v mnor 2>&1 | tee build.log
```

Each of the 160 phased VCF files was then annotated with SnpEff, generating a new compressed VCF:

```
java -Xmx16g -jar snpEff.jar -v mnor <phased.vcf.gz> -csvStats
<phased.annotated.vcf.gz.summary.tsv> -htmlStats
<phased.annotated.vcf.gz.summary.html> | pigz -p 4 -c - >
<phased.annotated.vcf.gz>
```

### *Estimating patterns of variation*

#### Levels of variation

From the SNPs, we computed the per-base pair population mutation parameter Watterson's theta ($\vartheta_W$; the number of segregating sites) and nucleotide diversity ($\pi$; the average number of pairwise nucleotide differences between a pair of chromosomes sampled from a population) as estimates of genetic variation across the whole population dataset and for each of the eight populations separately. These statistics were computed across non-overlapping windows of 1,000 bp or 100,000 bp, or across whole sequences. The effective length of each window was corrected for the number of actual accessible bases according to sequence depth (see above) and SNPs and accessible bases were further subdivided according to intergenic, UTRs, cds, synonymous/non-synonymous coding positions or intronic sequence to enable estimation of variation across different kinds of genomic regions. The 1 kb window estimates were used to map variation stepping away from genes, while the 100 kb estimates were used to compile genome-average statistics. In addition, we re-used code from `BioPerl` (*115*) to compute Tajima's D (*116*) by comparing $\vartheta_W$ and $\pi$, to characterize the degree to which the genome overall appeared to evolve neutrally ($D_T \approx 0$) or depart from neutrality ($D_T \neq 0$), which may indicate effects of population bottlenecks or selection.

Synonymous and non-synonymous variants were enumerated from the SnpEff annotations while the number of synonymous and non-synonymous sites across gene bodies were estimated using `KaKs_Calculator v1.2` (*75*). As in the window-based analyses, a correction was made for the per-base diversity estimate to take into account the number of accessible sites in the coding sequences.

#### Effective population size ($N_E$) and its historical demographic trends

We inferred long-term $N_E$ under mutation-drift equilibrium for an idealized population, assuming a snapping shrimp mutation rate of 2.64e$^{-9}$ substitutions per site and generation (*49*). This is to the best of our knowledge, the closest malacostracan species with a robust estimate of the mutation rate. We then used the standard equation $N_E = \vartheta_W / 4\mu$ to compute $N_E$ using the genome-wide average value for $\vartheta_W$. We inferred past changes in $N_E$ by estimating haplotype coalescent times using patterns of heterozygous genotypes in the reference specimen alone using two implementations of the Pairwise Sequentially Markovian Coalescent. In each case, we restricted analyses to span the distribution of heterozygosity across scaffolds or contigs longer than 500 kb, reusing the same SNP calls and masks of accessible sites (see above) as used for the full dataset.

I. `PSMC` (*27*): we first applied the Pairwise Sequentially Markovian Coalescent (PSMC) as implemented in `psmc v0.6.5-r67` https://github.com/lh3/psmc (n=4,911 scaffolds; 3.48 Gbp). Using a custom Perl script, we converted our depth of coverage genome mask to the window-based FASTA-like input format used in PSMC. To accommodate the high levels of variation in the krill, we used a window size of 10 bp instead of the default of 100 bp. We encoded each window with at least one accessible base with the symbol "T" (n=156,739,078), while fully inaccessible windows were encoded as "N" (n=169,545,821), and used the VCF files to re-code windows with heterozygous genotypes as "K" (n=21,271,007). We then applied the PSMC `splitfa` tool to split long sequences into fragments and ran 100 bootstrap replicates of PSMC, randomly resampling fragments in each replicate. We used a set of time segments with 12 free segments close to the present and ran each replicate for 25 iterations ("`-N25`"):

```
for NUM in {1..100}; do
psmc -N25 -t15 -r5 -b -p "1+1+10*1+15*2+4+6" -o round-${NUM}
<psmcfa.split>
done
```

II. `MSMC` (*117*): we then applied the Multiple Sequentially Markovian Coalescent (MSMC) algorithm for one sample as implemented in `mcmc2 v2.1.1` (https://github.com/stschiff/msmc2) (n=5,176 scaffolds; 3.63 Gb). For a single diploid sample, this method is similar to the original PSMC method above (*118*). First, we used a custom Perl script to scan the VCF files and our depth of coverage genome mask to generate the expected genotype input format while correcting for inaccessible sites between SNPs. We then ran MSMC2 with a fine-grained set of time segments:

```
msmc2 -t 40 -o <out> -p "10*2+100*1+1*2+1*3" <datasets/*.500kbp>
```

Alternative time series yielded similar profiles for $N_E$.

In both analyses, $N_E$ and the number of generations are re-scaled post-analysis by the per-generation mutation rate μ, which is unknown for this species. We therefore again applied the mutation rate inferred from snapping shrimp in (*49*) of $2.64e^{-9}$ substitutions per site per generation to scale the statistics, assuming a generation time of one year. For PSMC, we concatenated the results as per the online instructions and plotted the variation among the replicates with the PSMC tool `psmc_plot.pl`, allowing the program to auto-select the best-fitting iteration for each replicate

```
psmc_plot.pl -Y 300 -X 5000000 -p -s 10 -u 2.64e-09 -g 1 round-ALL.plot round-
ALL
```

We combined the output from both PSMC and MSMC into a single figure. In this figure we also incorporated the "LR04" benthic $\delta^{18}O$ foraminiferal calcite isotope stack, a record of data that indicates changes in global ice volume and deep ocean temperature across 5.3 million years (*119*). This data was downloaded from: https://lorraine-lisiecki.com/stack.html

### Counting alleles and estimating allele frequency divergence, population structure and selection

Our SNP dataset spanned Northern krill samples collected from eight geographical regions across its natural range (Fig. 1; table S1). We used a custom Perl script to compute allele frequencies at each SNP for each population and across the whole dataset using the genotypes in the VCF GT field in the phased VCF files. The allele counts and frequencies were saved in tabular TSV text files to enable fast access in downstream analyses, and used to compute the folded allele frequency spectrum across the whole dataset. We then estimated the pairwise genetic distances and interrelationships between all specimens (i.e. the $d_{XY}$ statistic), while correcting for accessible sites. Pairwise distances were converted into a neighbor-joining tree using SplitsTree (*120*).

Population structure and ancestry was inferred from the variation among samples using unsupervised PCA and admixture tools, without prior partitioning of samples. We first employed a custom script to subsample one percent of the variant sites, and then prune remaining variants based on linkage disequilibrium to remove correlation between variant sites in windows of 500 sites. Pruning was done with the function `sgkit.ld_prune` (https://pystatgen.github.io/sgkit, version 0.5.1) that generates a maximally independent set of variants. We used an $R^2$ threshold of 0.1, whereby no variant pairs below the threshold were retained. `PLINK v1.90b4.9` (*121*) was used to perform a PCA on the filtered variants with options '`--pca var-wts --double-id --chr-set 46`' whose output was used in a custom plotting function in python to generate plots. The filtered variant set was converted to PLINK binary biallelic genotype table (bed) format to use as input to `ADMIXTURE v1.3.0` (*122*). We varied the number of populations K from 2 to 8 and ran 50 repetitions using different seeds to control initial conditions. We collected the output files (suffix .Q) and uploaded zip archives to the CLUMPAK server (http://clumpak.tau.ac.il/) (*123*) to summarize the output from the 50 runs.

Specimens were next grouped according to their regions of origin. We used the $F_{ST}$ estimator by Reynolds *et al*. (*124*) to estimate pairwise allelic divergence across whole scaffolds or contigs and infer the genome-wide levels of divergence among all of the eight populations. The pairwise distances were converted into a NeighborNet network using SplitsTree. We next partitioned the data into two major contrasts representing krill from different regions:

1. The Atlantic Ocean (n=67 samples) vs. the Mediterranean Sea (Catalan Sea; n=7 samples).

2. The North Eastern North Atlantic Ocean (samples from waters around Iceland, the Barents Sea, Svalbard and Scandinavia; n=47 samples) vs. the South Western North Atlantic Ocean (samples from the Gulf St. Lawrence, Canada; and the Gulf of Maine, USA; n=20 samples)

For each contrast, Reynolds $F_{ST}$ was used to estimate divergence across non-overlapping windows of 100 bp, 1,000 bp or 100,000 bp, whole contigs/scaffolds or at the genome-scale, using a custom Perl script. We used the 1 kbp window-based $F_{ST}$-estimates to compare divergence between genes and flanking regions, and test for association with gene-sequence or selective sweep signals (see below).

The Weir-Cockerham estimator (*125*) was used to calculate the per-SNP $F_{ST}$, in order to visualize data and outlier loci and SNPs with unusually high levels of divergence. Outlier SNPs were also used to identify gene-level haplotypes of putatively selected gene-variants that segregated between ocean basins. For every contrast or population, the frequency of each haplotype was computed.

Results of these analyses are given in Figs 3–4 and figs. S23–25.

### Simulations of divergence

We performed coalescent simulations of neutral divergence under a simple population-split model using ms (*126*) to determine the probability of observing high levels of allelic divergence between basin-scale population samples from neutrality alone, in the absence of natural selection. In this Wright-Fisher model, populations would split without any subsequent exchange of alleles through gene flow and diverge over time assuming constant population size and no recombination.

For each of the two major contrasts (see previous section), we parameterized the simulation with the observed genome-wide estimates of divergence and variation. We first inferred the scaled time *T* and the number of generations *t* since the population split using the following two equations, respectively:

1. $T = \frac{-\ln(1-F_{ST})}{2}$ from (*124*). *T* was then converted into *t*:
2. $t = T * 4 N_E$, using our genome-wide estimation of $N_E$ (1.53 million).

The downsampling of data (see previous section) resulted in 7.35 M SNPs sampled across 8.43 Gb accessible sites across the genome, for an average block length of 1,146 bp per SNP. We therefore aimed to simulate the coalescent process in about 7.35 M 1.1 kbp loci per contrast (making minor adjustments in the case of invariant sites in any of the two contrasts) and export one SNP per locus. Our estimate of population mutation parameter Watterson's theta ($\vartheta_W$) was 1.62% per base, resulting in a $\vartheta_w$ estimate of 18.57 per 1,146 bp long locus. Using these conditions, we generated 7.3 M unlinked SNPs (the same number as the subsampled, unlinked empirical SNPs).

- The Atlantic Ocean vs. the Mediterranean Sea contrast (n=67 vs n=7 diploid samples)

  Genome-wide $F_{ST}$ had been estimated to be 0.056, resulting in estimates of *T*=0.02885 and *t*=176,572 generations. Taking *T*/2 as the measure of time since the split, we simulated 7,349,210 SNPs:

  ```
  ms 148 7349210 -t 18.57 -I 2 134 14 -ej 0.01443 2 1 -s 1 > simulated.out
  ```

  We here specified to sample 148 chromosomes from the population (two times the n=74 sample size), repeat the simulation 7,349,210 times, use the scaled population mutation rate ("`-t 18.57`"), sample two populations of 134 and 14 chromosomes each ("`-I 2 134 14`"), model a join between the two populations at generation time 0.01443 ("`-ej 0.01443`") and export one SNP per simulated locus ("`-s 1`").

- The North Eastern North Atlantic Ocean vs. the South Western North Atlantic Ocean (n=47 vs. n=20 diploid samples)

Genome-wide $F_{ST}$ had been estimated to be 0.0168 in this contrast, resulting in estimates of $T$=0.00825 and $t$=51,752 generations. We adjusted the number simulations to match the number of segregating sites in this sample and ran the matching simulation using the same approach as above:

```
ms 134 7211757 -t 18.57 -I 2 94 40 -ej $VAL 2 1 -s 1 > simulated.out
```

We converted the output from the ms simulations into allele counts and computed per-SNP $F_{ST}$ using the Weir-Cockerham estimator as above. We then compared the simulated $F_{ST}$-spectra to the observed (binning the SNPs in $F_{ST}$-bins of 0.1), in order to test for excess divergence compared to expectation under neutrality, which could be taken as evidence for natural selection.

#### Signatures of selective sweeps

For our two major pairwise contrasts (i: SW vs NE North Atlantic Ocean; ii: Atlantic Ocean vs Mediterranean Sea), we used the cross-population XP-nSL test in `selscan v1.3.0` (*127*) to scan for signatures of extended haplotypes in one group relative to the other. Such patterns could result from local selective sweeps through natural selection on an adaptive variant that reduces linked variation only in the focal population but not in the other, which is not subject to the selection pressure. XP-nSL can detect signatures of both hard and soft sweeps and help pinpoint candidate loci for ecological adaptation. It is conceptually similar to the XP-EHH test but does not require a genetic map or insight into recombination rates.

For each contrast, we first converted the VCF files to compressed TPED files. To limit the number of files on disk at a time, we then implemented a small daemon that generated one TPED subset per population and scaffold/contig and executed selscan:

```
selscan --xpnsl --threads 20 --trunc-ok --tped-ref
<seq_$N.ref_population.tped> --tped <seq_$N.other_population.tped> --out
<results_out/seq_$N.out>
```

For the first contrast, the NE North Atlantic Ocean sample (n=47) was taken as the reference population and the SW North Atlantic Ocean sample as the other population (n=20). For the second contrast, the Atlantic Ocean population (n=67) was taken as the reference population and the Mediterranean sample as the other population (n=7). In this scenario, negative XP-nSL scores for SNPs indicate extended haplotypes in the reference population, while positive scores are associated with the other population. We normalized the results with the selscan command "`norm`" against the genome-wide empirical background, such that extreme XP-nSL scores would be those less than -2 (indicating sweeps in the reference population) or more than 2 (indicating sweeps in the other population). We kept per-SNP XP-nSL scores and also computed the average across windows 1,000bp.

### Enrichment analyses

Genes were ranked from high to low exon-wide $F_{ST}$ and analyzed for enriched Biological Process gene ontologies using Flybase *Drosophila* homologues in GOrilla (*89*). Enrichments, p-values and FDR-corrected q-values were computed by GOrilla. We retained all reported GO:s, which had FDRs q-values of about 0.1 or less.

Results are provided in table S15.

### Estimation of haplotype ages

We estimated the ages of minor alleles on the divergent and putatively selected haplotypes to learn how long they may have been segregating in the Northern krill. For this, we used the nonparametric Genealogical Estimation of Variant Age (GEVA) coalescent method to estimate the time to the most recent common ancestor (TMRCA) between alleles (*128*). GEVA uses both information about mutation and recombination rates to model TMRCA between ancestral and derived alleles, but does require *a priori* assumptions about demographic history.

Recombination rates were not known in krill from before. We therefore used `iSMC v0.0.23` (*129*) to estimate a genome-average recombination rate from the reference specimen alone, as this tool has been shown to provide robust estimates even from single diploid samples. iSMC uses the coalescent with recombination to infer both recombination rates and aspects of demographic history from heterozygous genotypes.

For estimation of recombination rate, we used genotypes from the reference specimen along 650 scaffolds longer than 500 kb and that had 60% or more accessible bases. We provided a VCF and a matching genome mask file. We left most settings at the defaults in the parameter file and used five rho categories for spatially heterogeneous recombination along sequences ("number_rho_categories = 5"), tolerance for numerical optimisation at $1e^{-4}$ ("function_tolerance = 1e-4"), a window size for decoding of 1Mb (instead of the default 3Mb, "fragment_size = 1000000") and 40 threads ("number_threads = 40"). The program was run:

```
ismc params=1.merged_contigs.bpp 2>&1 | tee 1.merged_contigs.run.log
```

It infers the population recombination rate ρ, where $\rho=4*Ne*r$ , and $N_e$ is the effective population size and r is the recombination rate per base pair. Our analyses gave a ρ of 0.013. For the reference specimen and set of 652 sequences, we estimated $\vartheta_W$ to 1.1% and $N_e$ accordingly to 1.02 M. We thus computed $r$ to be $3.2e^{-09}$/bp or 0.32 cM/Mb across the 652 scaffolds, which we took as the genome-wide average.

We next prepared data for analysis in GEVA. We sought to analyze the age of variants on gene-haplotypes that diverge between Atlantic and Mediterranean ("at vs. me"; n=660 genes) samples or between SW and NE Atlantic samples ("we vs. ea"; n=34 genes) (our two major contrasts in these analyses). GEVA is designed to compute ages of derived alleles, which should be set as the ALT allele in re-coded VCF files. In our case, the ancestral and derived alleles were not known as we had not aligned outgroup sequences to the krill genome. We therefore instead re-coded the data assuming that the minor allele was derived (i.e. ALT) and the major allele was ancestral (i.e. REF) using a custom Perl script. For each contrast, we generated two sets of data, one set with the ALT allele taken as the minor allele in the first group (e.g. “at”) and the other set with the ALT allele taken as the minor allele in the second group (e.g. “me”), respectively.

We converted the re-coded VCFs into GEVAs binary format (one file per contig/scaffold containing a gene of interest), specifying the recombination rate:

```
geva_v1beta -t 2 --vcf <data.vcf> --out <data.out> --rec 3.2e-09
```

We then executed the program, specifying $N_e$ and mutation rates and using the program-provided Hidden-Markov files:

```
geva_v1beta -t 2 --Ne 1530000 --mut 2.64e-09 --hmm hmm_initial_probs.txt
hmm_emission_probs.txt
```

Allele ages of focal variants are estimated from a composite posterior distribution and saved in *.sites.txt output files. Each variant has an age based on a mutation clock (M), recombination clock (R) or joint clock (J). For all variants inside the gene coordinates of each gene, we collected the joint clock estimate and computed age distributions across all variants.

Results of these analyses are provided in Fig 3.

### Assessment of molecular evolution in *nrf-6* and the topology of its encoded protein

In our scans for signatures of selection in the Northern krill, a homologue of the *nose resistant to fluoxetine protein 6* (*nrf-6*) gene encoding the NRF-6 protein was top-ranked for high $F_{ST}$ across its exons. To further learn about how selection may have acted on its gene and corresponding protein, we performed a comparative scan for positive selection between haplotypes. We overlaid the SNP variants detected in the Mediterranean samples on the reference (Atlantic) *nrf-6* CDS sequence and aligned them together with the homologous KrillDB sequence for the Antarctic krill *E. superba* (accession: ESS142994) and the decapod *P. vannamei* sequence (NCBI accession: QCYY01000544) using MAFFT. We then produced a 4-way phylogenetic tree with FastTree and analysed the dataset (alignment+tree) in PAML under a free-ratio codon model to infer per-branch *dN/dS* ratios (*130*) and test for elevated *dN/dS* on the Mediterranean haplotype of the gene.

We predicted its protein structure with Alphafold2, using the same approach as implemented for the DNA methylation genes (see above). The *nrf-6* gene encodes a transmembrane acyltransferase that assists in lipid transportation (*131*). To predict the intracellular, transmembrane and extracellular regions of the protein and map the distribution of non-synonymous and synonymous variants along the regions, we used the TOPCONS server (*132*) and PPM v3.0 (positioning of proteins in membranes) (*133*). We used SignalP v6.0 (*134*) to predict the signal peptide. Protein figures were rendered using PyMOL (*135*) and ChimeraX (*136*). The coordinates reported by TOPCONS were then used to annotate the corresponding exons and locations of the detected SNPs.

**Supplementary Text**

### *Geographical variation in the Northern krill and regions used to sample material*

*M. norvegica* occurs over a wide geographic range (Fig. 1) (*78*). Notable general physiological and behavioral characteristics and variations of the Northern krill across this range are that they: i) tend to grow larger and be more fecund in fjord-areas compared to pelagic areas (*137*); ii) adjust to similar levels of oxygen consumption and metabolic rates across widely different prevailing ambient thermal conditions (*78*, *138*); iii) initiate the reproductive season early (winter/spring) in the South and late (spring/summer) in the North to time spawning with phytoplankton blooms (*139*, *140*); iv) tend to have a more carnivorous diet in the North and mixed omnivorous diet in the South (*141*); and iv) prefer deeper depths and perform greater diurnal vertical migrations in oligotrophic (i.e. brighter) environments as a means to avoid predation (*78*, *142*, *143*).

#### Gulf of Maine (USA) - The SW North Atlantic Ocean

The samples collected from the Gulf of Maine (USA) were part of the NSF OCE-1316040 grant and were, together with the Canadian samples, used to characterize the Northern krill from the South-Western range of its distribution. The Gulf of Maine is a large gulf, or inland sea, in the NE coast of the USA, bounded by Massachusetts, New Hampshire, Maine, and the Canadian provinces of New Brunswick and Nova Scotia by land, meanwhile in the ocean limits with Georges Bank and Browns Bank (*144*). There are three major basins within the gulf (Jordan, Georges and Wilkinson Basins – the latter reaching below 250 m depth). At the Wilkinson Basin, where the samples were taken, surface water temperatures oscillate between 16 °C in the summer to about 7 °C in the winter. Below 50 m depth, water stays between 5 °C and 7 °C all year long (*145*, *146*). Similar profiles are found in all the Gulf of Maine, although lower temperatures can be reachedfound in areas under strong influence of the Labrador Sea water. Salinity ranges from 32.0 for those water masses of Scotian Shelf Water origin, to 34.6 in those typical from the Labrador Sea Water and 35.6 of the Warm Slope Water (*146*).

The Gulf of Maine is a high productivity area, with a strong spring bloom, summer surface stratification and fall mixing. Despite the important role and abundance of *M. norvegica* in the Gulf of Maine ecosystem, they show strong net avoidance due to their large size and swimming ability, therefore their true distribution, seasonality and long term-trends are not very well known (*144*, *147*).

#### Gulf of Saint Lawrence (Canada) - The SW North Atlantic Ocean

The Gulf of Saint Lawrence samples were collected within the Atlantic Zone Monitoring Program (AZMP). The Gulf of Saint Lawrence is a semi-enclosed sea in South-East Canada. The deep Laurentian Channel runs from the continental shelf to the mouth of the Saint Lawrence River, surrounded by more shallow basins. In the winter, surface water temperature reaches below 0 °C, with temperature in depth ranging from below 0 °C (in shallow areas) to 8 °C in areas under the influence of slope waters entering the gulf, especially deep channels and basins. In contrast, during the summer, warmer waters are found at the surface (9 °C and warmer), while temperatures oscillate between 2 and 8 °C at greater depths (*148*). Salinity is lower in the surface in summer, and in areas under the influence of the discharge of the Saint Lawrence river, meanwhile is relatively constant (about 34) in depth all year long (*148*).

In this region, *M. norvegica* forms loose aggregations in the deeper areas (*149*), showing feeding preference for copepods, especially *Calanus* spp., accumulating lipids during the fall and early winter from the lipid-rich copepods in diapause, and consuming the reservoirs in spring and summer (*150*).

### Masfjord (Norway) - The NE North Atlantic Ocean

The samples included in the study from Norwegian fjords were taken in Masfjorden, Nordhordaland on the second of October 2019 during a week-long cruise to threshold Fjords with the research vessel, Kristine Bonnevie. The Masfjord is a 24-kilometer long east-to-west directed fjord that empties into the inner part of the larger, and more open, Fensfjorden. It is between 500 to 1,500 meters wide, has a maximum depth of 494 m and the sill depth is 75 m, and is surrounded by relatively steep mountain sides. At the collection site (60°52.575909' N 5°26.34410' E), it is 421 meters deep and the samples were taken with a plankton net (MIK) at a depth between 202 and 306 meters. The temperature at the collection depth was measured to 8.5 °C and the salinity to 35.

*M. norvegica* is a dominant macroplankton in the Masfjord and particularly abundant in the head of the fjord, likely being advected there. They show a diurnal vertical migration pattern in the fjord, with most biomass concentrated in the upper 100 m during winter nights (*151*).

### Gullmar Fjord (Sweden) - The NE North Atlantic Ocean

The Gullmar Fjord (sv. Gullmarsfjorden) is a narrow (1–2 km wide) sill fjord located on the Swedish west coast. It is 28 kilometers long and has a maximum depth of about 120 m, the basin with depths <100 m being about 5 kilometers long. The fjord is characterized by a unique combination of hydrographic conditions, being influenced by both the northward Baltic current, freshwater runoff and North Sea water. It has a brackish surface/upper layer (salinities between 24 and 27), an intermediate layer with salinities of 32–33 between 15–50 m and comparably stagnant deep layer with salinities 34–35 and temperatures that mostly fluctuate between 4–8 °C, and has suffered multiple low-oxygen events in the last 100 years (*152*). The krill for this study was collected in the deep area, near Alsbäck-Fossen, in July 2018.

*M. norvegica* is the dominant euphausiid of the Gullmar Fjord and shows a diurnal vertical migration pattern with krill typically entering the upper/mid layers in the nighttime, and adults typically occurring at deeper depths than juveniles (*153*). It was suggested this habitat is unusual in the sense that both beneficial access to nutrients and risk from predation are greater at deeper depths, reverse to most Northern krill habitats and that local vertical migration patterns reflect adjustment to local conditions (*153*). The krill has high spawning and molting rates in the late summer in this area, followed by oosorption and reproductive diapause in the fall-winter (*137*).

#### Iceland - The NE North Atlantic Ocean

The samples from the Icelandic area were collected during the Icelandic spring cruise in May which is a monitoring survey where measurements are made on hydrography, nutrients, phytoplankton, and zooplankton around the island. The system of oceanic ridges on which Iceland rests divides the oceanic area around the island into different ocean regions or domains (*154*). There is great variability in hydrographic conditions between these regions that in turn affect the distribution, composition and productivity of zooplankton (*155–157*). To the south and west is the Atlantic Domain, characterized by relatively warm and saline Atlantic water with near surface winter temperatures (0-50 m) of ~6°C, and summer temperatures of ~12°C. To the north is the Atlantic/Arctic Domain where the water is usually a mixture of Atlantic and Arctic water. Near surface temperatures are usually ~1°C during winter but may reach ~6°C during summer. To the east is the Arctic Domain with near surface temperatures usually below 0°C and may reach ~4–5°C during summer. The samples for this study were collected in all these three main hydrographic domains.

In Icelandic waters four euphausiid species are most abundant (*158*). *Thysanoessa inermis* tends to be most abundant over the shelves, *M. norvegica* over the shelf edges, while *T. longicaudata* is mainly found in offshore areas. The fourth abundant euphausiid species, *T. raschi* is most common in fjords on the northwest, north and east coasts. In Icelandic waters, *M. norvegica* appears to have a life span of 2 years (*159*).

#### Barents Sea & Svalbard - The NE North Atlantic Ocean

The high-latitude Barents Sea is characterized by a mix of warm Atlantic water (>3 °C), cold Arctic water (<0 °C) and warm coastal waters (>3 °C). The shelf area is about 1.6 million $km^2$ with a mean depth of 230 m, and the region is experiencing extensive warming and "Atlantification" through influx of Atlantic waters (*160*). Krill make up a large part of the mesopelagic biomass and several *Thysanoessa* spp. are native to the region. *M. norvegica* is thought to be increasingly transported into the Barents Sea from the Norwegian Sea (*161*), and is also being advected into the Svalbard Archipelago and fjords (*162*, *163*). The Arctic Ocean represents the northernmost range of the species and no evidence as of yet has been presented indicating it has formed resident reproducing populations in this area.

#### Spain - The Mediterranean Sea

The Mediterranean sea is in general an oligotrophic area. It has constant temperature and salinity in the whole water column at all depths above 150–200 m and epipelagic water temperatures are between 13–28 °C in summer. This high temperature of the water column, in comparison to the neighboring Atlantic Ocean, may provoke a faster degradation of organic matter arriving to the Deep Sea bottom.

In the Mediterranean, *M. norvegica* has northern distribution. The lowest temperature where *M. norvegica* lives and completes its life cycle in the western Mediterranean is ~13 °C. Their highest densities in daytime are at depths >400–500 m*, though at night it occupies surface waters in the epipelagic zone (*164*, *165*), where also larvae (specially early stages) are mainly located (*166*). There are some seasonal patterns in its density: in late winter–early spring, when the species breeds, it can aggregate inshore at the shelf-slope break (*166*, *167*). In the fall, it moves deeper and can in autumn reach a second peak of abundance, downloading to 1000 m (*167*).

*M. norvegica* is the major prey of the Mediterranean fin whale (*168*), and it also occurs in the diet of deep-sea fish and shrimps, including near-bottom fish like juvenile *Merluccius merluccius* at the shelf-slope break (*169*). At higher depths, to 1200 m, especially in autumn, a high variety of deep-sea bottom living fish (e.g. sharks) and crustaceans prey on it (*170*, *171*). This has also been reported in Atlantic deep waters (*172*, *173*).

*as in the Bay of Cádiz, Atlantic Ocean.

#### *No signature of whole-genome duplication (WGD) in the Northern krill*

To test for evidence of WGD in the krill, we investigated duplicated genes. The rate of gene family expansion is high compared to many crustaceans (fig. S8B; table S9), but there is no conclusive evidence from neutral divergence ($K_S$) among paralogs that many of them would have originated at the same time through WGD. Divergence-patterns are similar among all studied crustaceans. A shoulder in the $K_S$-distribution in krill is better modeled by multiple underlying distributions than one (fig. S8C–D; table S10), favoring smaller-scale duplications (*91*). In vertebrates, horseshoe crabs and arachnids that have undergone WGD, many Hox gene ohnologs have been retained even after diploidization (*96*). We found 9 out 10 core Hox genes (*lab*, *pb*, *Hox3*, *Dfd*, *Scr*, *ftz*, *Antp*, *Abd-A*, *Abd-B*, but not *Ubx;* fig. S11) in the krill genome but all but one are single-copy, not supporting WGD. Three putative *Hox3*-like paralogs were found. We therefore find that genome expansion in the krill is likely associated with TE proliferation and multiple small-scale duplications rather than WGD.

### Supplementary figures

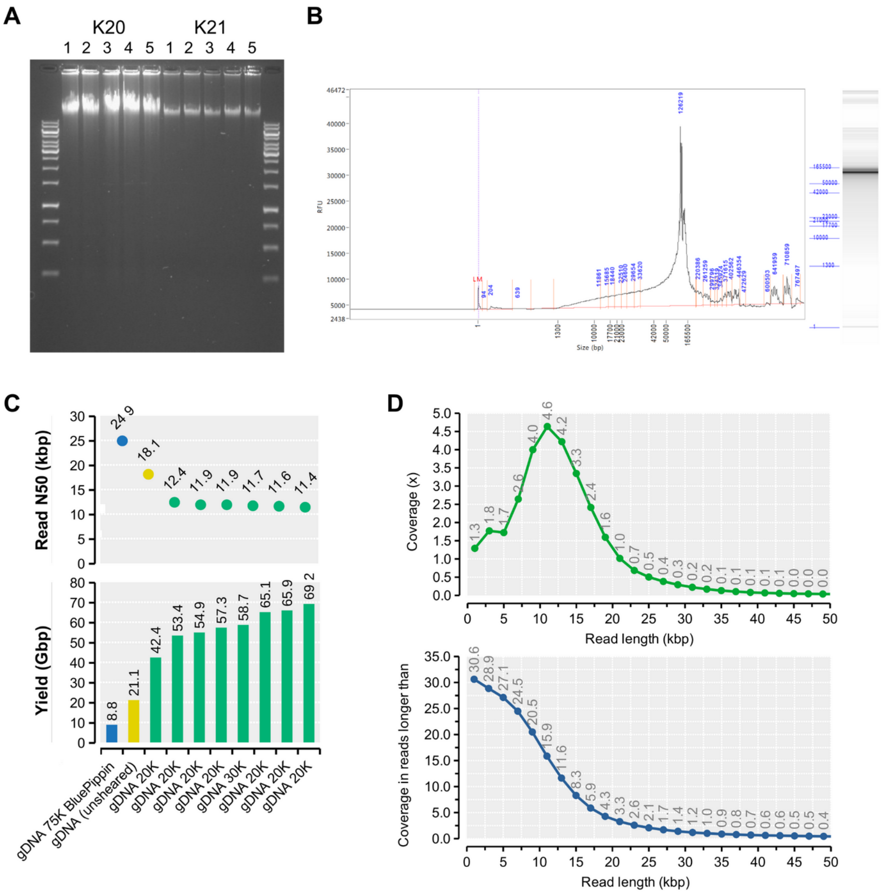


**Fig. S1.**

Genomic Tip DNA extraction of the reference specimen sample K20 used for long-read and linked-read sequencing, as well as backup sample K21. (**A**) Agarose gel image of samples K20 and K21 (ladder: Thermo Scientific GeneRuler™1 kb DNA ladder). (**B**) Femto Pulse fragment length readout of sample K20. (**C**) Nanopore flow cell yields using size-selected, unsheared or sheared DNA molecules. Shearing was done to fragments of 20 or 30 kb. (**D**) Genome coverage for PromethION reads of different length intervals. Top=genome coverage for a particular length class. Bottom=genome coverage for a particular length class or longer reads.

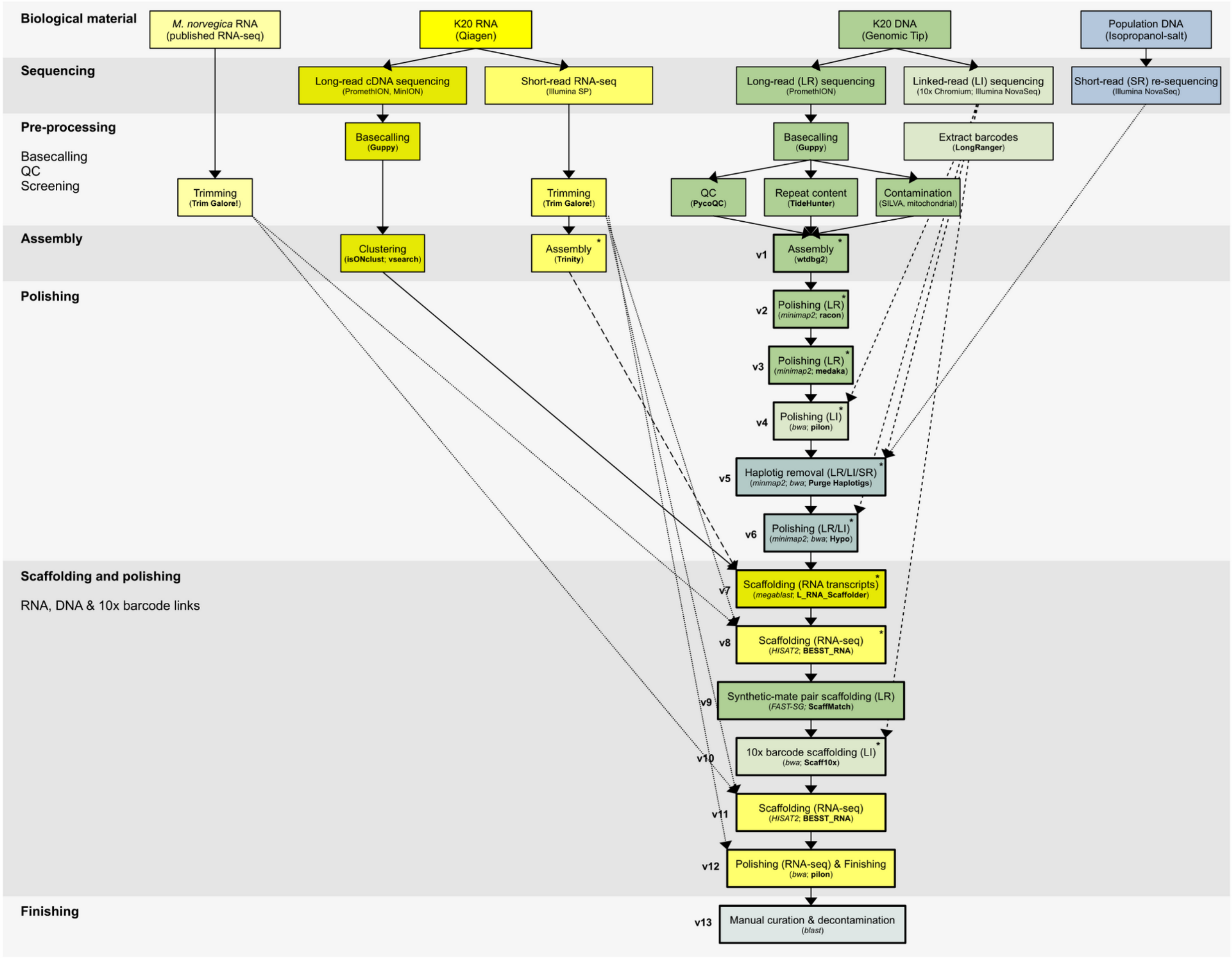


**Fig. S2.**

Genome assembly workflow overview. Biological material included RNA data from published samples, RNA and DNA data from the reference specimen “K20” and population re-sequencing data. For the first two steps (Biological materials and Sequencing), extraction and sequencing methods are indicated in parentheses, while main bioinformatics tools are indicated in subsequent steps. Pre-processing of data encompassed basecalling, trimming, processing of barcodes and basic analytics including quality control (QC) and screening for contaminants (Screening). Colors of boxes indicate important data types: yellow=data and steps involving RNA (e.g. scaffolding and annotation); green=data and steps reliant on DNA derived from the reference specimen (e.g. assembly and polishing); light blue=population data. Arrows indicate how the RNA and DNA materials were used throughout the genome assembly workflow. A final round of manual curation and removal of putative bacterial contaminants generated assembly version 13.

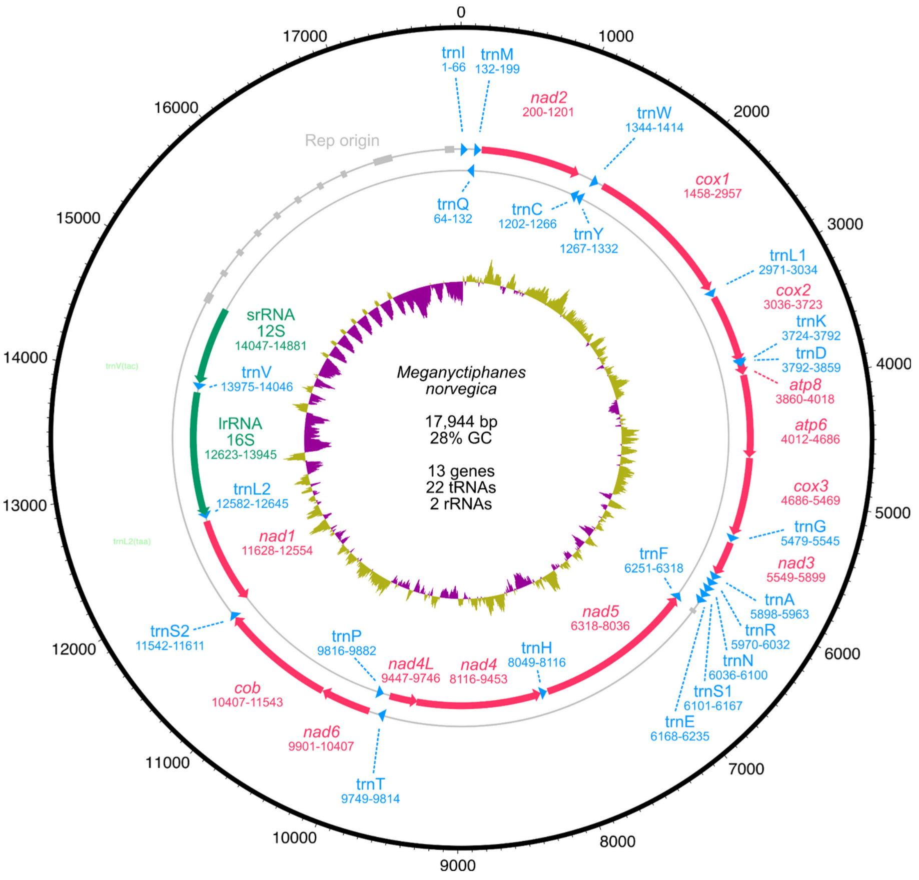


**Fig. S3.**

A 17 kbp mitochondrial chromosome assembly and features derived from PromethION long-reads from the K20 reference specimen. Coordinates are indicated on the outer track (bold black line). Inner histogram indicates GC-content (pink=%GC below mean; green=%GC above mean). Detected features are indicated on the outer heavy/plus strand (clockwise) or the light/minus strand (counter-clockwise), respectively: red arrows=protein coding genes; green arrows=ribosomal RNAs (rRNA); blue arrows=transfer RNA (tRNA); gray arrows=origin of replication and/or AT-rich repeats). Summary statistics are indicated at the center of the plot.

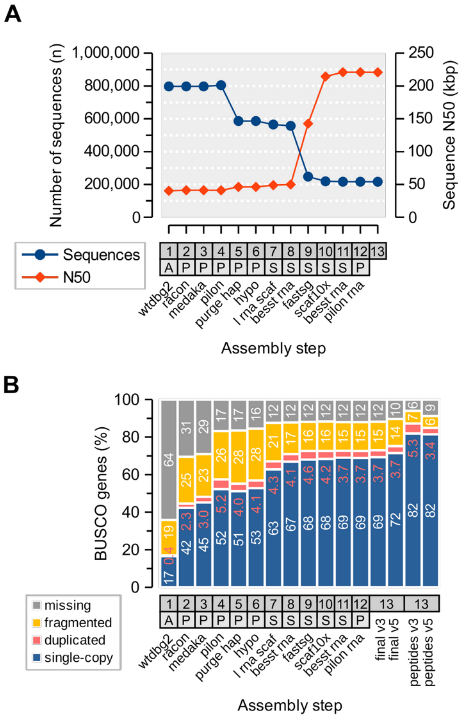


**Fig. S4.**

Genome assembly metrics. (**A**) Number and N50 lengths of genome sequences throughout the assembly pipeline. Steps 1–12 involved bioinformatic tasks (A=assembly; P=polishing; S=scaffolding) while step 13 was a mix of manual curation and bioinformatics to accomplish the final assembly. (**B**) BUSCO gene detection analysis using the Arthropod odb9 gene set and BUSCO v3 in genome mode (steps 1–12) or both BUSCOv3/odb9 and BUSCOv5/odb10 in both genome mode or peptide mode.

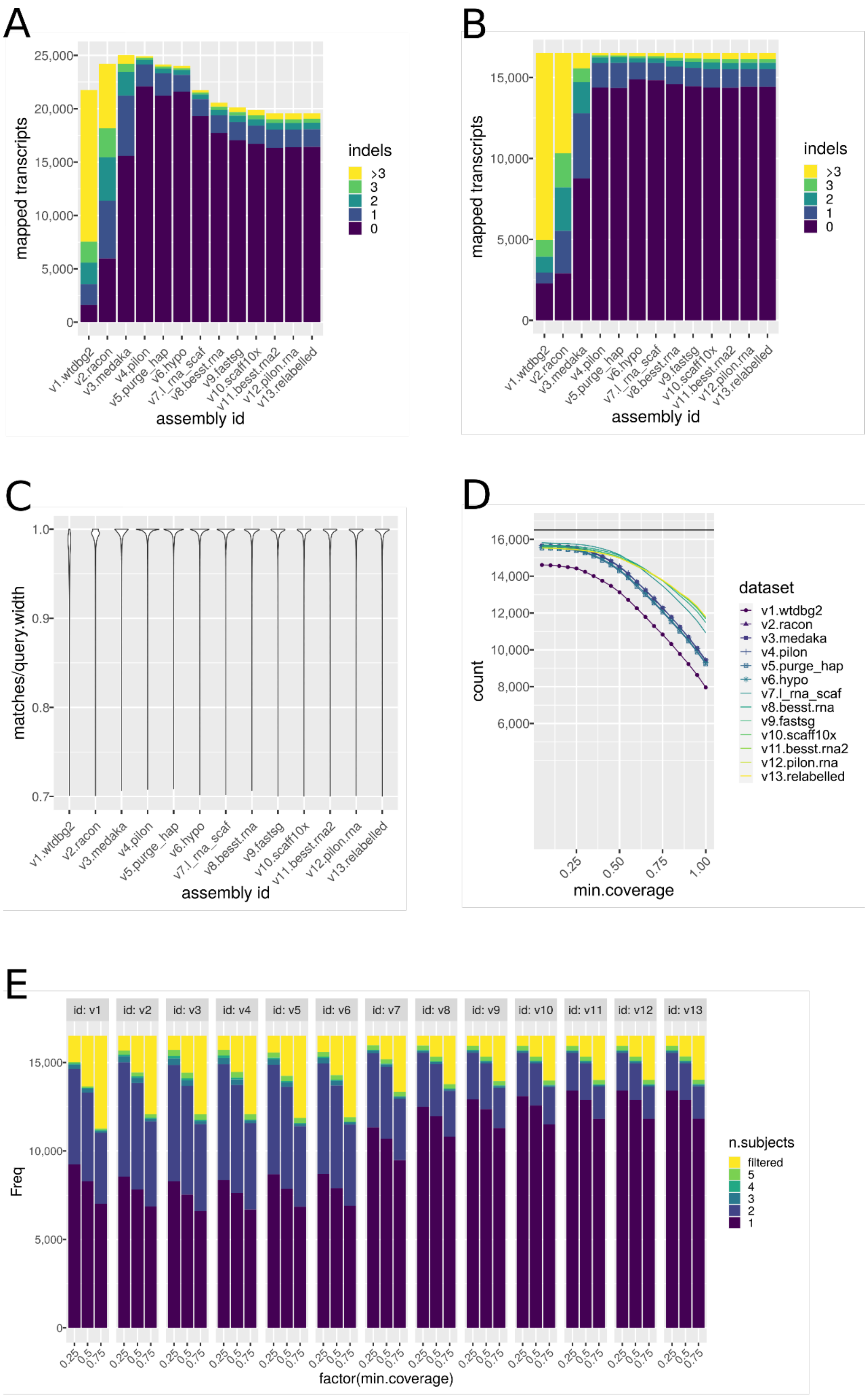


**Fig. S5.**

Genome assembly metrics based on mapping properties of Trinity transcripts (n=16,509 input sequences). Distribution of insertion-deletions (indels) over mapped transcripts (**A-B**) for redundant mappings (**A**) and averaged over each transcript (**B**). The numbers of indels decrease rapidly for the first rounds of polishing and thereafter remain stable during the subsequent scaffolding steps. Similarly, the number of mismatches decreases as a result of polishing (**C**). The number of transcripts mapping to multiple contigs decreases as a result of scaffolding (**D**), indicating an improvement in assembly contiguity. Assembly completeness is assessed by looking at the number of gene bodies covered to a certain extent at 90% identity (**E**).

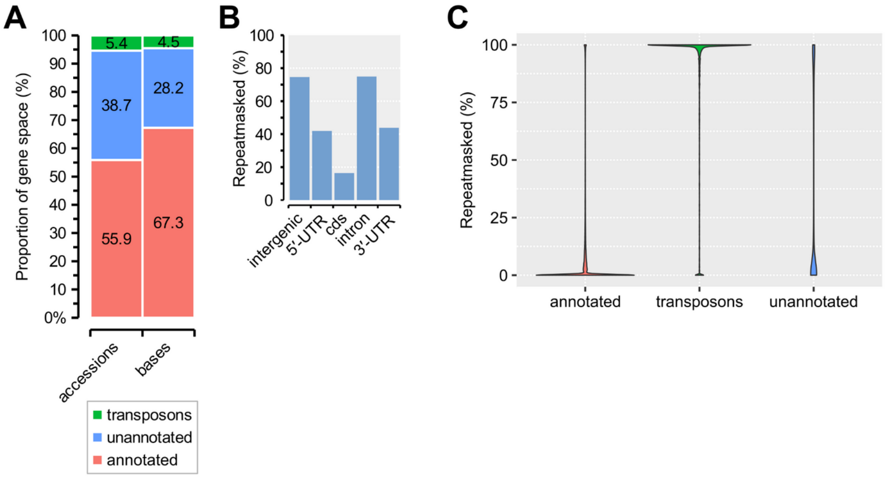


**Fig. S6.**

Overview of 42,227 putative protein-coding gene models in the krill genome detected through RNA and comparative data and functionally annotated using homology information in EnTAP. (**A**) The proportion of regular genes ("annotated"; n=25,301), transposons (n=2,283) and unannotated genes (n=14,643) following functional analysis with EnTAP and searches for transposon keywords ("accessions"=proportion of the total number of all genes; "bases"=proportion of the total number of bases of the coding sequence of all genes). (**B**) The genome-average repeat content estimated for each genomic region (UTR=untranslated exonic regions; cds=coding sequence) estimated with a custom library of interspersed repeats using RepeatMasker. For example, about ~15% of the coding sequence appears repetitive. (**C**) The per-gene distribution of repeat content across the 42,227 gene models. Categories as in (A). The bimodal repeat-content among unannotated genes hints at thousands of additional genes and TEs.

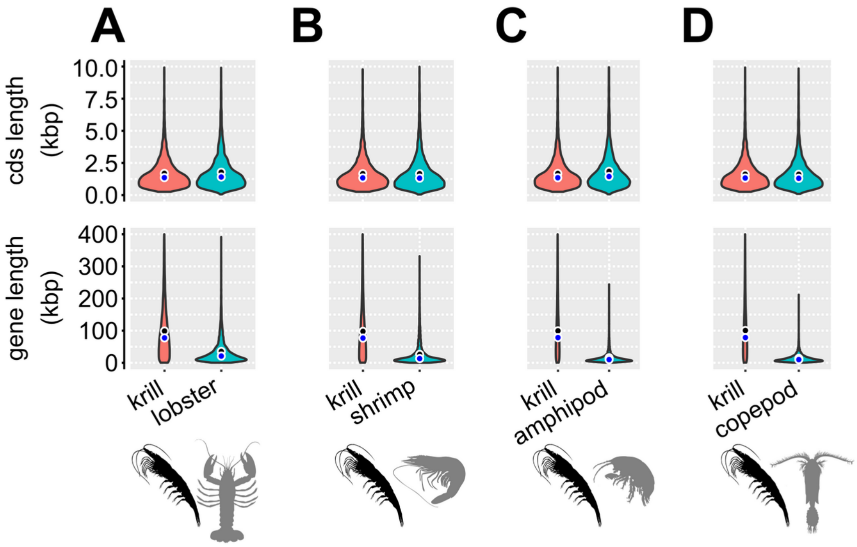


**Fig. S7.**

Comparison of the lengths of genes and coding sequences of 1:1 orthologs between *M. norvegica* and other crustacean species. (**A**) Comparison between *M. norvegica* vs. American lobster *H. americanus* based on 7,150 1:1 orthologs. Upper plot: distribution of the lengths of coding sequences (cds). Black circle indicates mean cds and blue circle indicates median cds. Lower plot: lengths of the full gene bodies in each genome. Mean and medians as in the upper plot. (**B**) Comparison between *M. norvegica* vs. Black tiger shrimp *P. monodon* based on 7,084 1:1 orthologs. Plots and scales as in (A). (**C**) Comparison between *M. norvegica* vs. the amphipod *H. azteca* based on 5,836 1:1 orthologs. Plots and scales as in (A). (**D**) Comparison between *M. norvegica* vs. the copepod *E. affinis* based on 4,551 1:1 orthologs. Plots and scales as in (A).

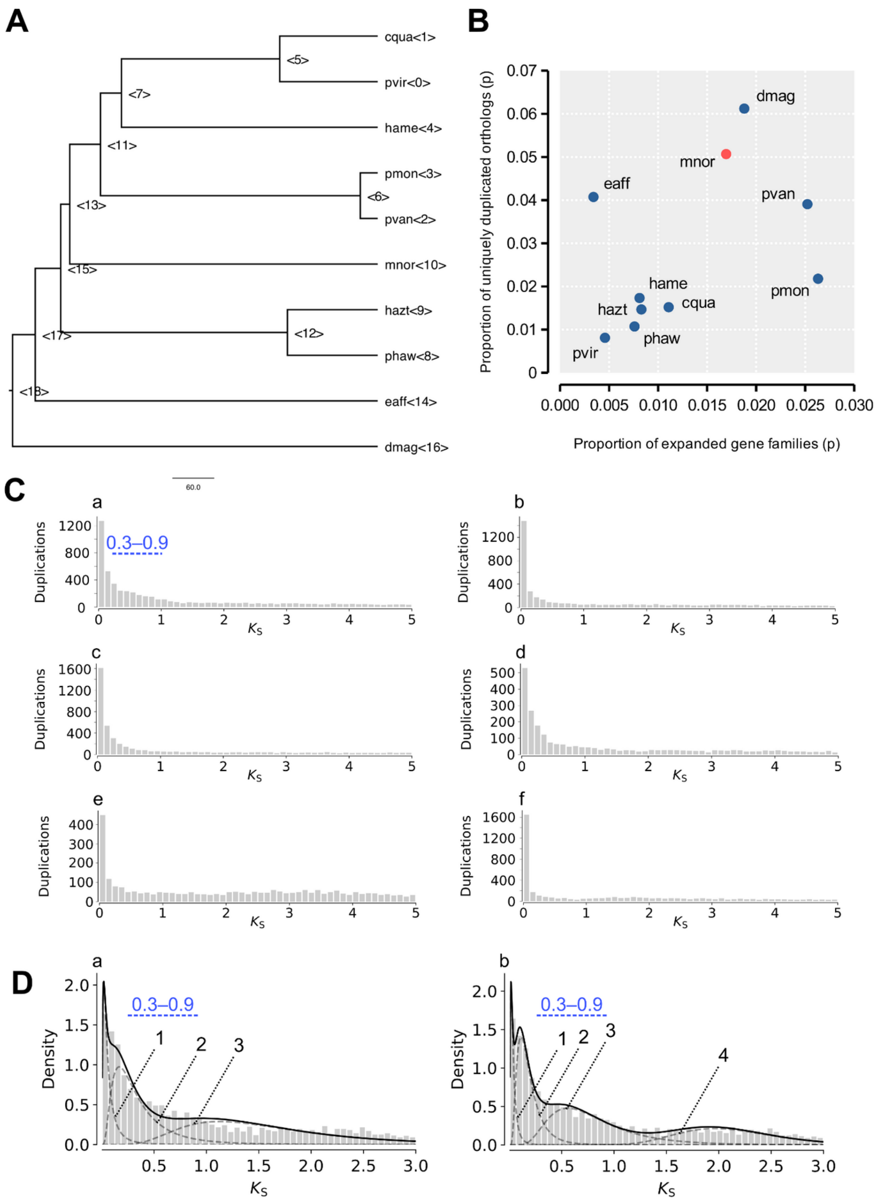


### Fig. S8.

Gene family expansion, gene duplication rates and synonymous (presumably neutral) divergence among gene family paralogs across crustaceans. (**A**) Phylogenetic time-tree used in gene family expansion analyses in CAFE. Node and tip labels are given in brackets and match those in table S11. Labels: mnor=*Meganyctiphanes norvegica*; hame=*Homarus americanus*; cqua=*Cherax quadricarinatus*; pvir=*Procambarus virginalis*; pmon=*Penaeus monodon*; pvan=*Penaeus vannamei*; hazt=*Hyalella azteca*; phaw=*Parhyale hawaiensis*; Eaff=*Eurytemora affinis*; Dmag=*Daphnia magna*. (**B**) Dot plot of lineage-specific duplicated genes and rapidly evolving gene families. X-axis: the rate of gene family expansion, measured as the proportion of rapidly evolving gene families relative to all gene families in that species and inferred through phylogenetic analysis in CAFE ($p<0.05$ that these families are evolving at the same rate as the genome-wide average background rate). Y-axis: proportion of lineage-specific expanded gene families for which orthologs are otherwise only found as single-copy genes in other species (allowing for two species with missing data). Labels as in A. (**C**) The distribution of synonymous divergence ($K_S$) among gene family paralogs in six species using node-weighted calculations in gene family trees. Sub-panels are: a) *M. norvegica*; b) *H. americanus*; c) *P. monodon*; d) *H. azteca*; e) *E. affinis*; f) *D. magna*. A potential peak or shoulder within the $K_S$-range from 0.3 to 0.9 in the krill is highlighted in blue. (**D**) Results of fitting Gaussian mixture models (GMMs) to the observed $K_S$-distribution in the krill using either three components (a) or four components (b) visualized with dashed and dotted lines.

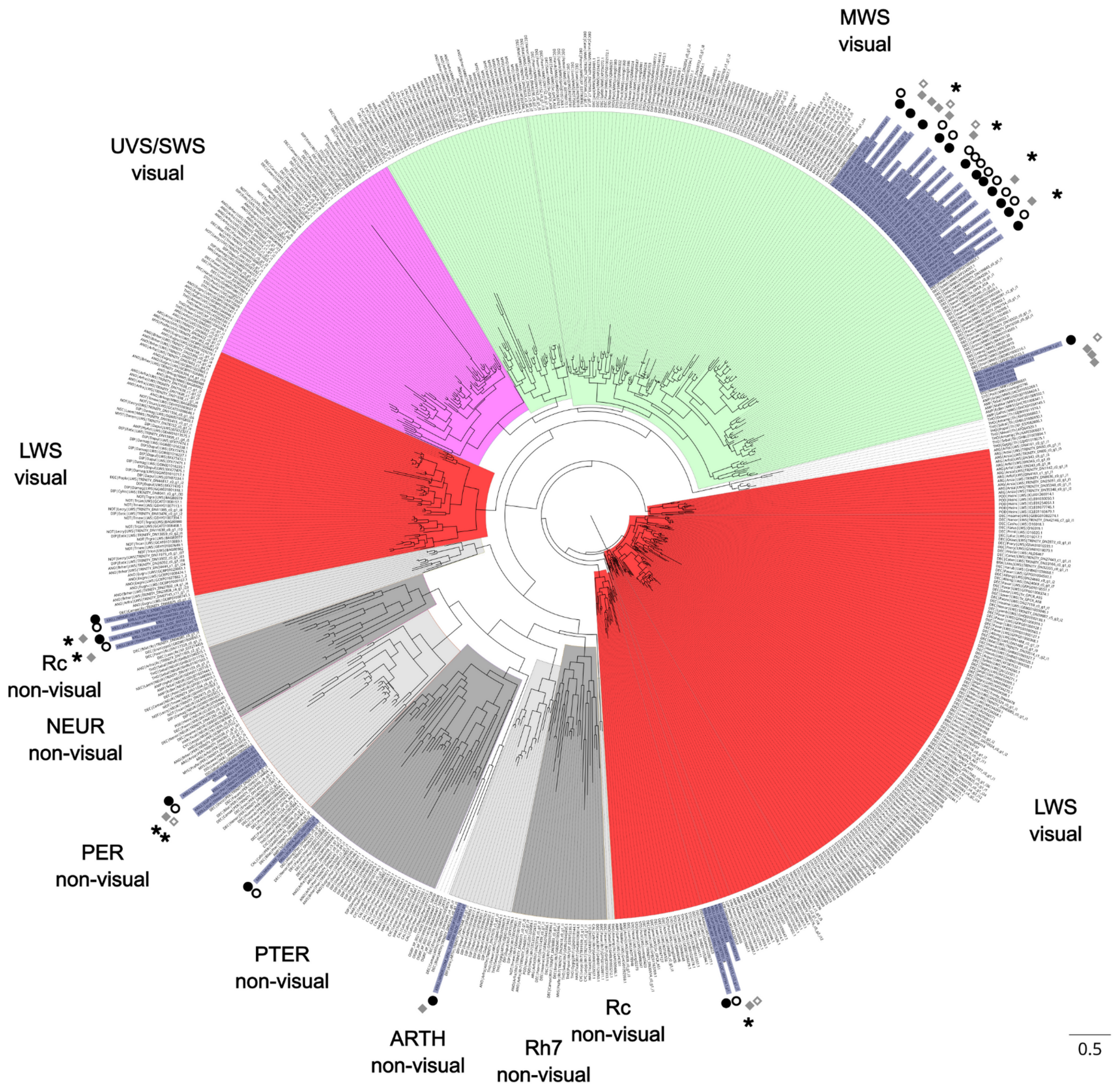


**Fig. S9.**

A maximum-likelihood tree of crustacean opsin gene family proteins. Genes from the Northern krill *M. norvegica* genome assembly and transcripts from Antarctic krill *E. superba* from Urso *et al.* (*174*) were added to the crustacean dataset of Palecanda *et al*. (*175*). Major opsin groups including visual or non-visual function indicated are: LWS=long wavelength-sensitive; MWS=middle wavelength-sensitive; UVS=ultraviolet/short wavelength-sensitive; Rc=crustaceanrhabdomeric opsin; Rh7=rhodopsin 7; ARTH=arthropsins; NEUR=neuropsins; PTER=pteropsins; PER=peropsins. Black circles=*M. norvegica* genes; White circles=*M. norvegica* transcripts from Palecanda *et al*.; Grey diamonds=*E. superba* transcripts from Urso *et al.*; White diamonds=*E. superba* markers from Palecanda *et al*.; stars=*Thysanoessa inermis* krill transcripts from Palecanda *et al*.

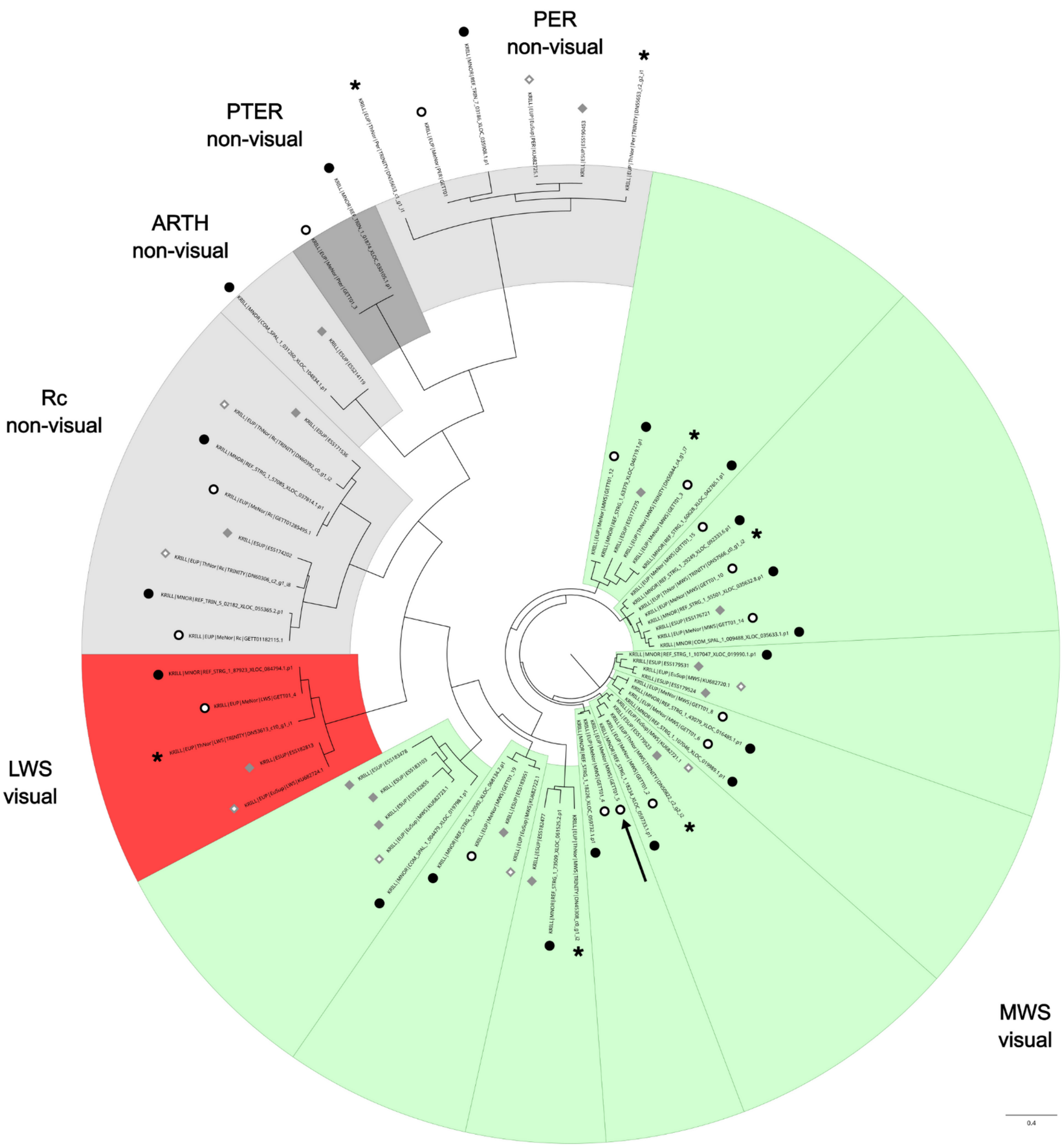


**Fig. S10.**
A maximum-likelihood tree of opsin gene family proteins in the krill. Genes from the Northern krill *M. norvegica* genome assembly and transcripts from Antarctic krill *E. superba* from Urso *et al.* (*174*) were added to the krill subset of the dataset from Palecanda *et al*. (*175*). Major opsin groups including visual or non-visual function indicated are: LWS=long wavelength-sensitive; MWS=middle wavelength-sensitive; Rc=crustaceanrhabdomeric opsin; ARTH=arthropsins; PTER=pteropsins; PER=peropsins. Black circles=*M. norvegica* genes; White circles=*M. norvegica* transcripts from Palecanda *et al*.; Grey diamonds=*E. superba* transcripts from Urso *et al.*; White diamonds=*E. superba* markers from Palecanda *et al*.; stars=*Thysanoessa inermis* krill transcripts from Palecanda *et al*. Arrow indicates an *M. norvegica* RNA transcript that is not unambiguously paired with a gene in the *M. norvegica* genome assembly.

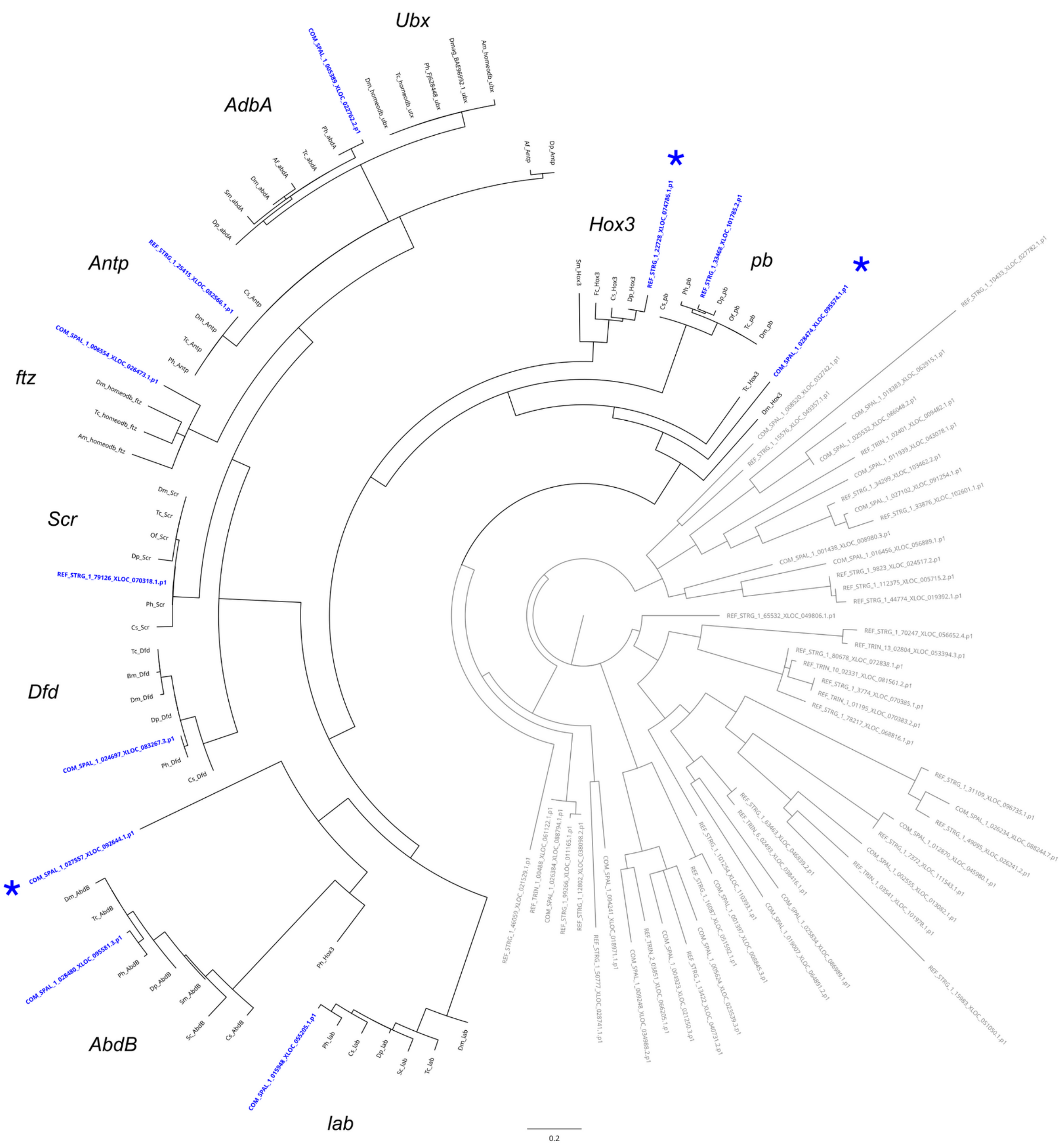


**Fig. S11.**

Phylogenetic inference of Hox genes in the Northern krill genome. A maximum likelihood phylogenetic tree was inferred from homeodomain motifs of 10 core Hox genes in arthropods (crustaceans+insects) taxa together with homeodomain gene candidates in *M. norvegica*. Blue labels indicate Northern krill accessions that fall within the Hox gene clade and respective Hox-gene branches. Non-hox *M. norvegica* accessions in gray. Genes: *lab=labial*; *pb=proboscipedia*; *Hox3*, *Dfd=Deformed*; *Scr=Sex combs reduced*; *ftz=fushi tarazu*, *Antp=Antennapedia*; *Abd-A=abdominal-A*; *Abd-B=Abdominal-B*; *Ubx=Ultrabithorax*. Nine out ten Hox genes were detected in krill, eight of which as single-copy genes. Three putative *Hox3*-like paralogs are indicated with stars. Species: Ag=*Anopheles gambiae*; Af=*Artemia franciscana*; Am=*Apis* *mellifera*; Bm=*Bombyx mori*; Cs=*Cupiennius salei*; Dp=*Daphnia pulex*; Dm=*Drosophila melanogaster*; Dmag=*Daphina magna*; Of=*Oncopeltus fasciatus*; Ph=*Parhyale hawaiensis*; Ps=*Porcellio scaber*; Pc=*Procambarus clarkii*; Sc=*Sacculina carcini*; Sm=*Strigamia maritima*; Td=*Thermobia domestica*; Tc=*Tribolium castaneum*.

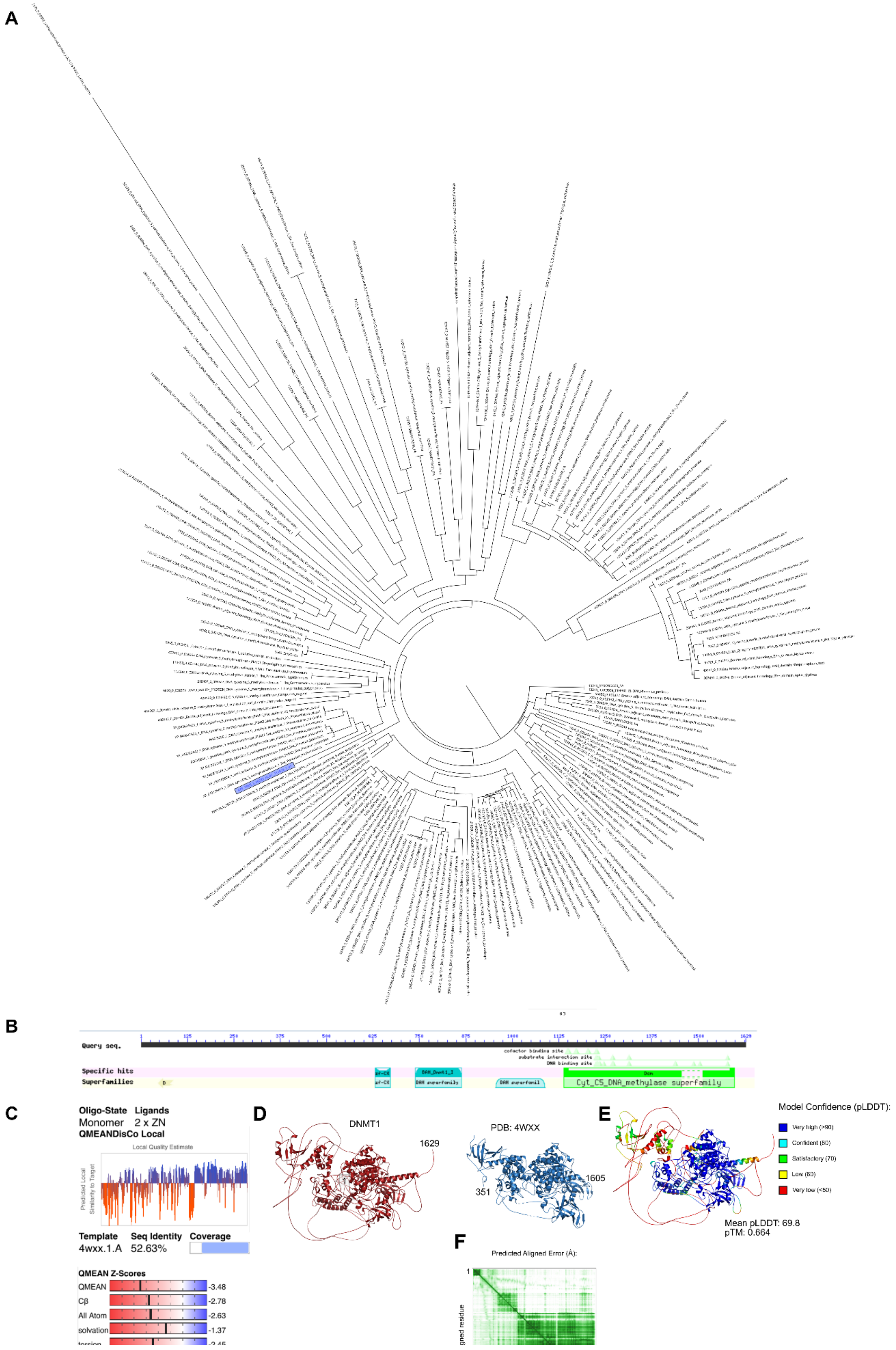
A
B
Query seq.
cofactor binding site
substrate interaction site
DNA binding site
Specific hits
Superfamilies
zf-CX
BAH_Dnmt1_I
BAH superfamily
BAH superfamil
Dcm
Cyt_C5_DNA_methylase superfamily
C
Oligo-State Ligands
Monomer 2 x ZN
QMEANDisCo Local
Local Quality Estimate
Predicted Local Similarity to Target
Template Seq Identity Coverage
4wxx.1.A 52.63%
QMEAN Z-Scores
QMEAN -3.48
Cβ -2.78
All Atom -2.63
solvation -1.37
D
DNMT1
1629
PDB: 4WXX
1605
351
E
Model Confidence (pLDDT):
Mean pLDDT: 69.8
pTM: 0.664
F
Predicted Aligned Error (Å):
1

**Fig. S12.**
Comparative and functional analysis of a candidate gene encoding DNMT1 in *M. norvegica* (REF_TRIN_3_00283_XLOC_111019.1.p1). The gene candidate was first detected in the EnTAP annotations and then analyzed for arthropod homologs. (**A**) Phylogenetic analysis using orthologs from OrthoDB, EggNOG and NCBI. Blue label indicates the position of the candidate krill sequence. (**B**) Domains indicated from querying the peptide sequence against the NCBI Conserved Domains database. (**C**) The best homology hit (model 4wxx.1.A) from querying the peptide sequence against the SWISS-MODEL database. (**D–F**) Modeling of the DNMT1 protein. (**D**) Comparison of the ColabFold predicted DNMT1 protein model (red) to the crystal structure of the human DNMT1 (PDB: 4WXX, blue) (*176*). (**E**) Per-residue confidence coloring of the top ranked predicted model of DNMT1. The mean predicted local distance difference test (pLDDT) value and pTM score are annotated. (**F**) Residue-residue alignment plot of the predicted DNMT1 model.

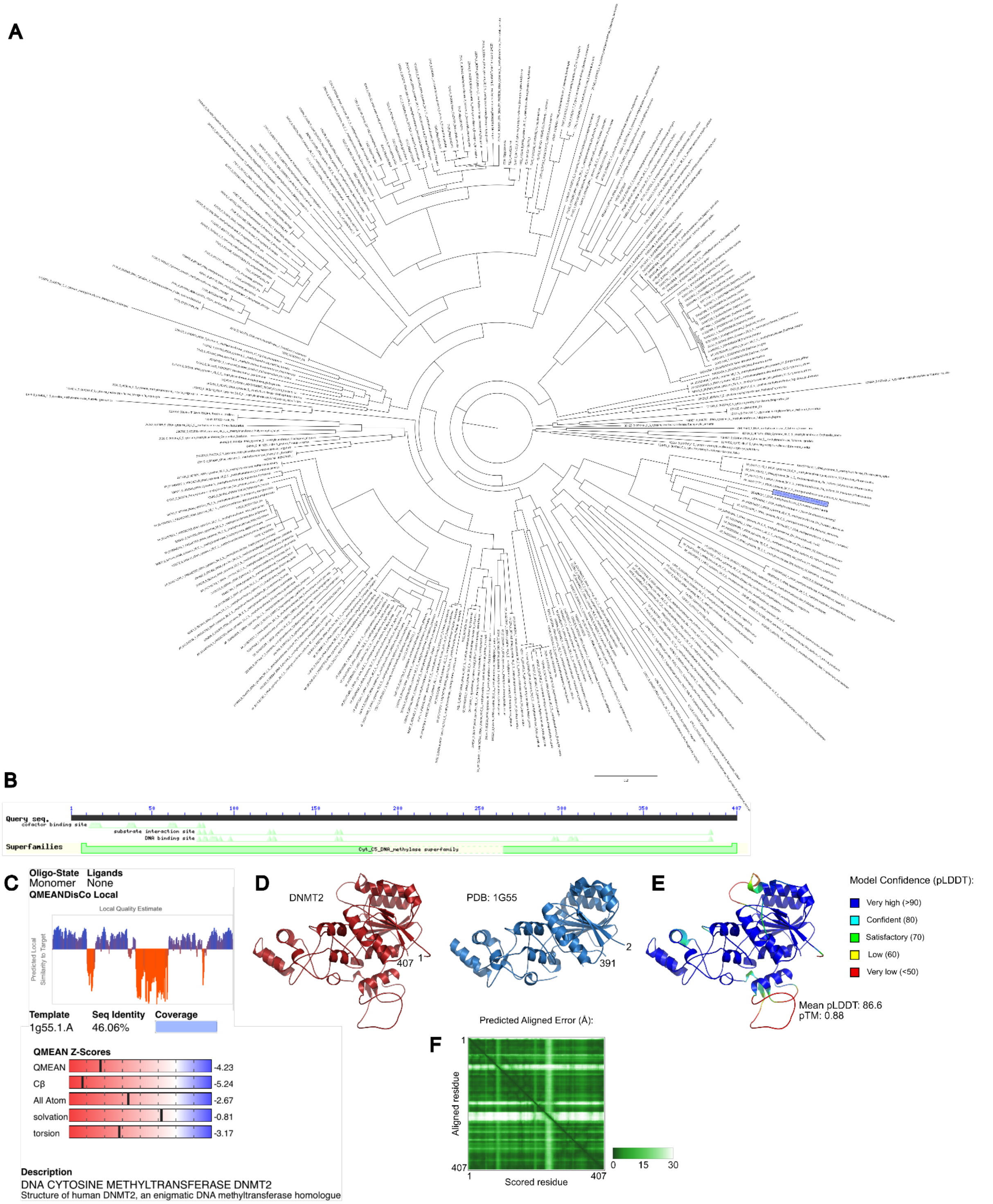


**Fig. S13.**

Comparative and functional analysis of a candidate gene encoding DNMT2 in *M. norvegica* (COM_SPAL_1_008092_XLOC_031468.8.p1). The gene candidate was first detected in the EnTAP annotations and then analyzed against arthropod homologs. (**A**) Phylogenetic analysis using orthologs from OrthoDB, EggNOG and NCBI. Blue label indicates the position of the candidate krill sequence. (**B**) Domains indicated from querying the peptide sequence against the NCBI Conserved Domains database. (**C**) The best homology hit (model 1g55.1.A) from querying the peptide sequence against the SWISS-MODEL database. (**D–F**) Modeling of the DNMT2 protein. (**D**) Comparison of the ColabFold predicted DNMT2 protein model (red) to the crystal structure of the human DNMT2 (PDB: 1G55, blue) (*177*). (**E**) Per-residue confidence coloring of the top ranked predicted model of DNMT2. The mean predicted local distance difference test (pLDDT) value and pTM score are annotated. (**F**) Residue-residue alignment plot of the predicted DNMT2 model.

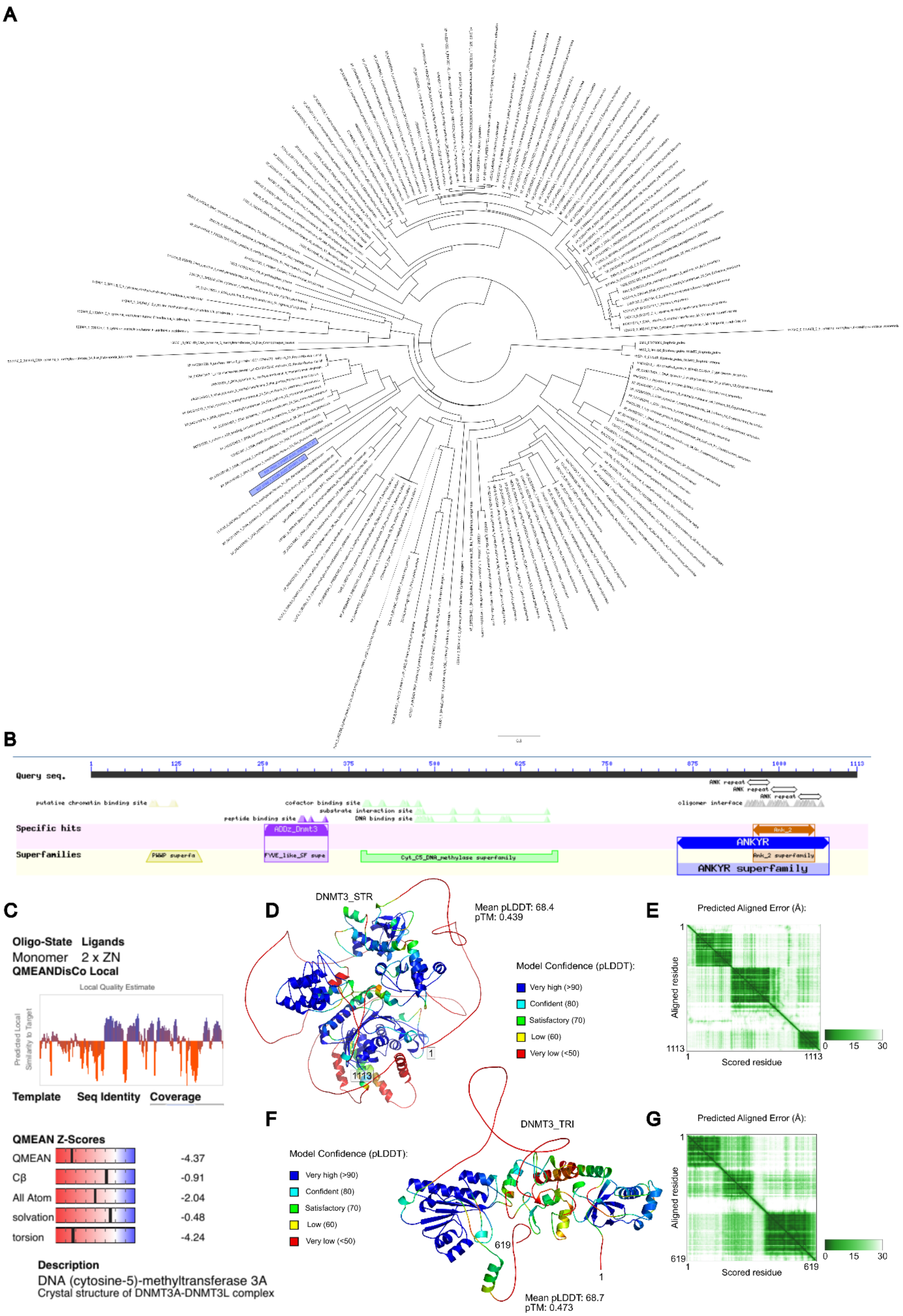
A
B
Query seq.
1
125
250
375
500
625
750
875
1000
1113
putative chromatin binding site
cofactor binding site
substrate interaction site
peptide binding site
DNA binding site
ANK repeat
ANK repeat
ANK repeat
oligomer interface
Specific hits
ADDz_Dnmt3
Ank_2
Superfamilies
PWWP superfa
FYVE_like_SF supe
Cyt_C5_DNA_methylase superfamily
ANKYR
Ank_2 superfamily
ANKYR superfamily
C
Oligo-State Ligands
Monomer 2 x ZN
QMEANDisCo Local
Local Quality Estimate
Predicted Local Similarity to Target
Template Seq Identity Coverage
QMEAN Z-Scores
QMEAN -4.37
Cβ -0.91
All Atom -2.04
solvation -0.48
torsion -4.24
Description
DNA (cytosine-5)-methyltransferase 3A
Crystal structure of DNMT3A-DNMT3L complex
D
DNMT3_STR
Mean pLDDT: 68.4
pTM: 0.439
Model Confidence (pLDDT):
Very high (>90)
Confident (80)
Satisfactory (70)
Low (60)
Very low (<50)
1
1113
E
Predicted Aligned Error (Å):
Aligned residue
Scored residue
1
1113
0 15 30
F
DNMT3_TRI
Model Confidence (pLDDT):
Very high (>90)
Confident (80)
Satisfactory (70)
Low (60)
Very low (<50)
619
1
Mean pLDDT: 68.7
pTM: 0.473
G
Predicted Aligned Error (Å):
Aligned residue
Scored residue
1
619
0 15 30

**Fig. S14.**

Comparative and functional analysis of two candidate genes encoding DNMT3 homologs in *M. norvegica* (REF_TRIN_3_01687_XLOC_055556_11.p1 and REF_STRG_1_22585_XLOC_074301_1.p1). The gene candidates were first detected in the EnTAP annotations and then analyzed against arthropod homologs. (**A**) Phylogenetic analysis using orthologs from OrthoDB, EggNOG and NCBI. One dotted branch has been shortened for presentation. Blue label indicates the position of the candidate krill sequence. (**B**) Domains indicated from querying the REF_STRG_1_22585_XLOC_074301_1.p1 peptide sequence against the NCBI Conserved Domains database. (**C**) The best homology hit (model 4u7p.1.A) from querying the peptide sequence REF_STRG_1_22585_XLOC_074301_1.p1 against the SWISS-MODEL database. (**D–F**) Modeling of the REF_STRG_1_22585_XLOC_074301_1 protein. (**D**) Per-residue confidence coloring of the top ranked predicted model of DNMT3_STR. The mean predicted local distance difference test (pLDDT) value and pTM score are annotated. (**E**) Residue-residue alignment plot of the predicted DNMT3_STR model. (**F–G**) Modeling of the REF_TRIN_3_01687_XLOC_055556_11 protein. (**F**) Per-residue confidence coloring of the top ranked predicted model of DNMT3_TRI. The mean predicted local distance difference test (pLDDT) value and pTM score are annotated. (**G**) Residue-residue alignment plot of the predicted DNMT3_TRI model.

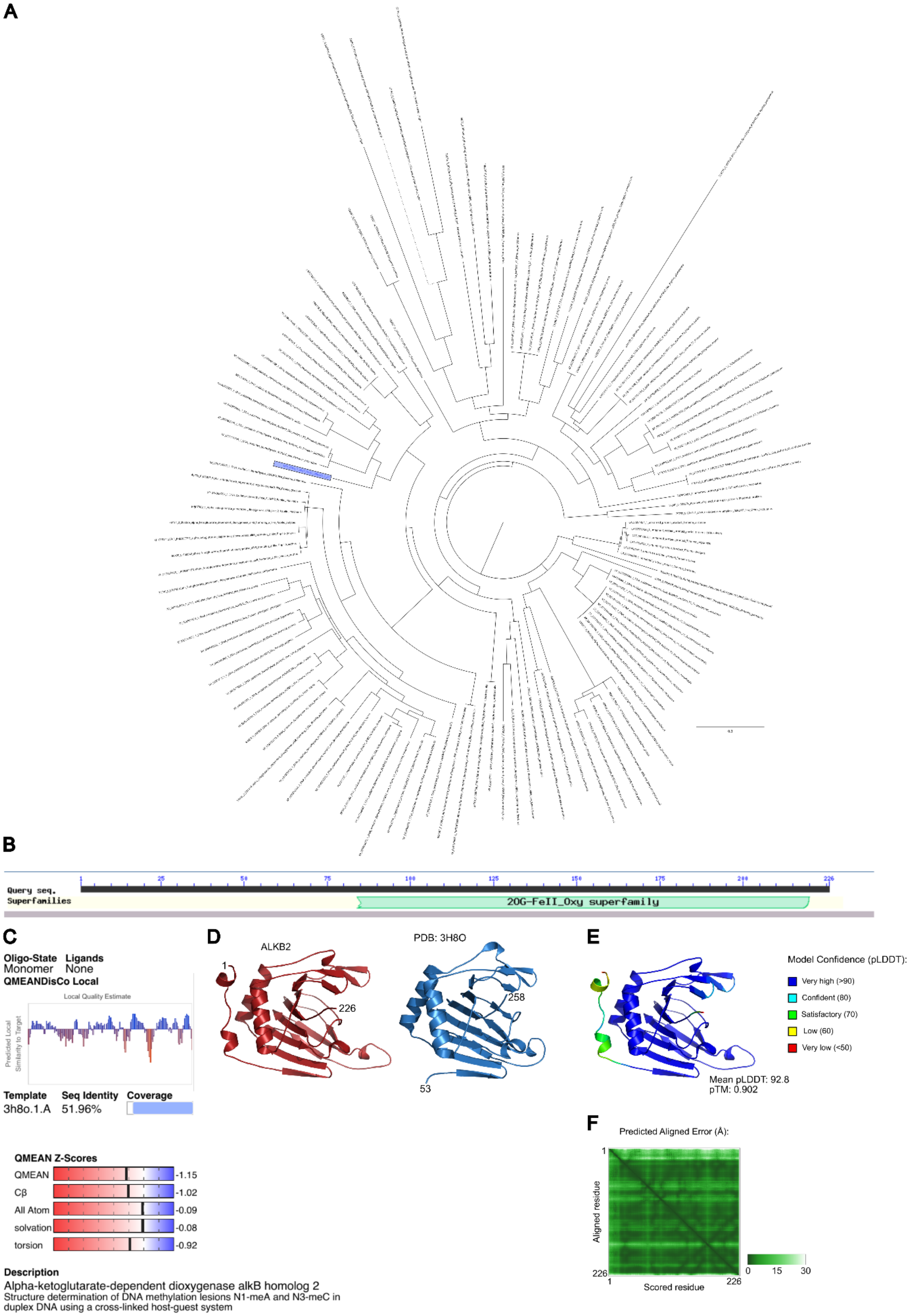
A
B
Query seq.
Superfamilies
1
25
50
75
100
125
150
175
200
226
2OG-FeII_Oxy superfamily
C
Oligo-State
Ligands
Monomer
None
QMEANDisCo Local
Local Quality Estimate
Predicted Local Similarity to Target
Template
Seq Identity
Coverage
3h8o.1.A
51.96%
QMEAN Z-Scores
QMEAN -1.15
Cβ -1.02
All Atom -0.09
solvation -0.08
torsion -0.92
Description
Alpha-ketoglutarate-dependent dioxygenase alkB homolog 2
Structure determination of DNA methylation lesions N1-meA and N3-meC in duplex DNA using a cross-linked host-guest system
D
ALKB2
1
226
PDB: 3H8O
258
53
E
Model Confidence (pLDDT):
Very high (>90)
Confident (80)
Satisfactory (70)
Low (60)
Very low (<50)
Mean pLDDT: 92.8
pTM: 0.902
F
Predicted Aligned Error (Å):
Aligned residue
Scored residue
1
226
0
15
30

**Fig. S15.**
Comparative and functional analysis of a candidate gene encoding ALKB2 in *M. norvegica* (COM_SPAL_1_008092_XLOC_031468.8.p1). The gene candidate was first detected in the EnTAP annotations and then analyzed against arthropod homologs. (**A**) Phylogenetic analysis using orthologs from OrthoDB and NCBI. Blue label indicates the position of the candidate krill sequence. (**B**) Domains indicated from querying the peptide sequence against the NCBI Conserved Domains database. (**C**) The best homology hit (model 3h8o.1.A) from querying the peptide sequence against the SWISS-MODEL database. (**D–F**) Modeling of the ALKB2 protein. (**D**) Comparison of the ColabFold predicted ALKB2 protein model (red) to the crystal structure of the human ALKB2 (PDB: 3H8O, blue) (*178*). (**E**) Per-residue confidence coloring of the top ranked predicted model of ALKB2. The mean predicted local distance difference test (pLDDT) value and pTM score are annotated. (**F**) Residue-residue alignment plot of the predicted ALKB2 model.

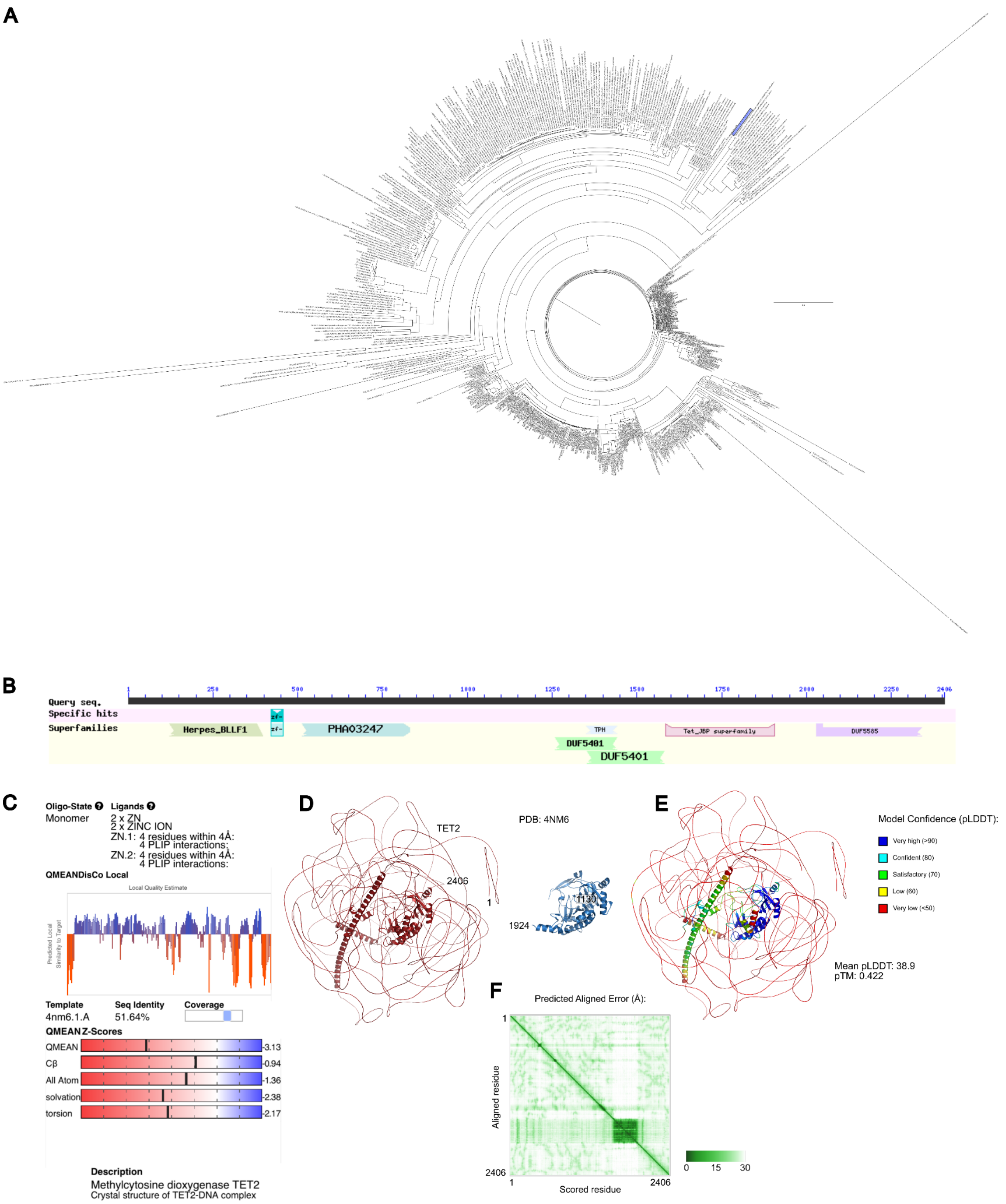


**Fig. S16.**

Comparative and functional analysis of a candidate gene encoding TET2 in *M. norvegica* (REF_STRG_1_37441_XLOC_109451.1.p1). The gene candidates were first detected in the EnTAP annotations and then analyzed against arthropod homologs. (**A**) Phylogenetic analysis using orthologs from OrthoDB and EggNOG. Blue label indicates the position of the candidate krill sequence. (**B**) Domains indicated from querying the peptide sequence against the NCBI

Conserved Domains database. (**C**) The best homology hit (model 4nm6.1.A) from querying the peptide sequence against the SWISS-MODEL database. (**D–F**) Modeling of the TET2 protein. (**D**) Comparison of the ColabFold predicted TET2 protein model (red) to the crystal structure of the human TET2 (PDB: 4NM6, blue) (*179*). (**E**) Per-residue confidence coloring of the top ranked predicted model of TET2. The mean predicted local distance difference test (pLDDT) value and pTM score are annotated. (**F**) Residue-residue alignment plot of the predicted TET2 model.

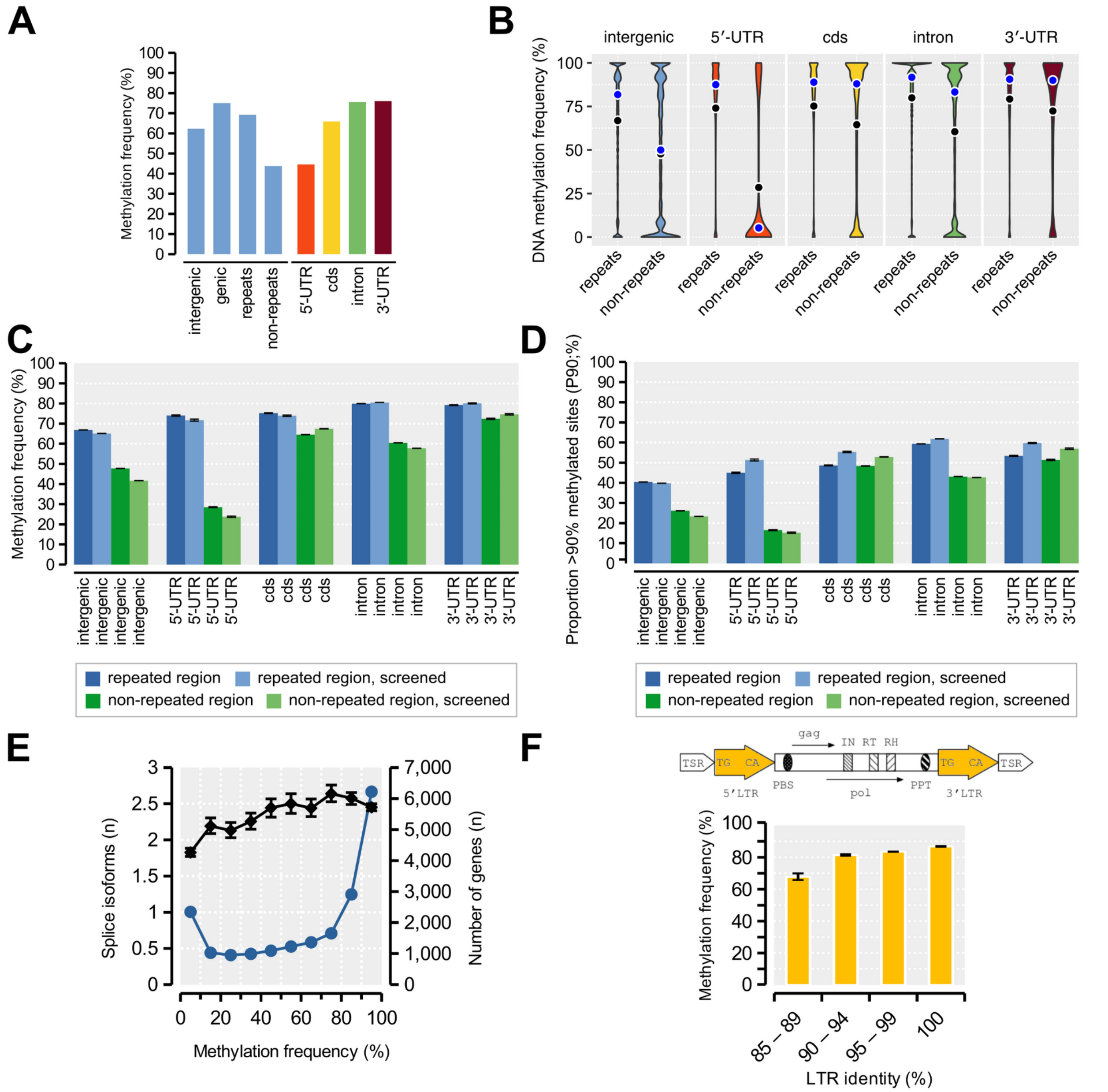


**Fig. S17.**

Cytosine methylation frequencies detected at CpG sites through Nanopore signal analysis. (**A**) Overall DNA methylation frequencies (proportion of methylated Nanopore reads) in genic vs. intergenic regions, repeated vs. non-repeated regions and gene-body regions. (**B**) Full distributions of DNA methylation rates across repeated vs. non-repeated regions, partitioned by intergenic and genic regions. (**C**) Mean DNA methylation frequencies across intergenic and genic gene regions. Screened bars indicate measurement only across CpG sites that have been screened to exclude: i) heterozygous genotypes in the reference specimen; ii) regions occurring outside of the approved read-depth boundaries in the population genomic dataset. Bar plot whiskers indicate 95% confidence intervals generated from 200 non-parametric bootstrap pseudo-replicates. (**D**) As (C) but showing P90 values as defined in (A). P90 is the proportion of

CpG sites with >90% methylated reads. (**E**) The average number of RNA splice isoforms per gene as a function of average methylation rate in exons binned for intervals of 10% (y1-axis). Whiskers indicate 95% confidence intervals generated from 1000 non-parametric bootstrap pseudo-replicates. Number of genes in each bin shown on the y2-axis. (**F**) Average CpG-methylation rate across 1,706 putative LTR retrotransposons detected with LTR_Retriever and tested to contain at least one expected LTR domain. The model of a retrotransposon (top) indicates the location of the 5’-LTR and 3’-LTR regions (yellow; modified after a model in the LTR_Finder program manual) that were measured for identity for each element and used to bin elements into intervals of 5% divergence (low divergence between LTRs represent evolutionarily young repeats that may recently have been inserted into their respective genomic locations). Bar plot whiskers indicate 95% confidence intervals generated from 200 bootstrap replicates. The group of retrotransposons with 100% identical LTRs were found to have 1.28× times higher than those with 85–90% LTR identities (87% vs. 68%; $p<0.05$).

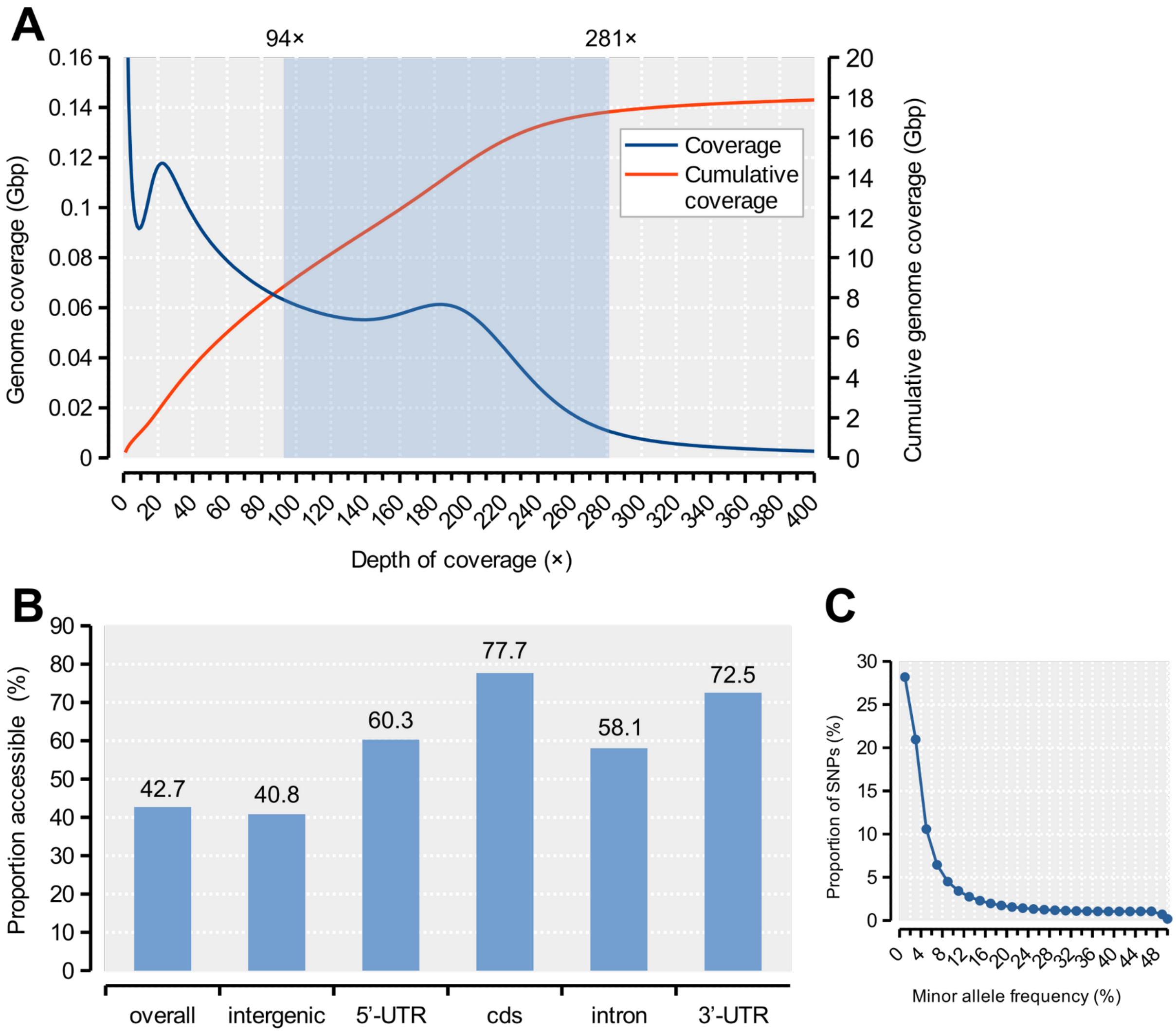


**Fig. S18.**
Regions of the genome used to estimate levels and patterns of variation from SNPs. (**A**) A putative diploid peak of mapped short-read re-sequencing data was identified at ~188× depth of coverage (with minimum mapping qualities at 10 or more). Upper and lower thresholds at +50% (281×) and -50% (94×) around the peak were used to define the region of the genome accessible for analysis of variation (blue area). (**B**) In addition to the threshold in (A), a second criterion of at least 50% of samples being genotyped at a site was applied. The bars indicate the proportion of accessible sites in different genome regions given the thresholds specified in (A) and the minimum genotyping rate, spanning 8.4 Gb in total. (**C**) The resulting folded allele frequency spectrum of all SNPs after applying the quality, depth and sample coverage filters.

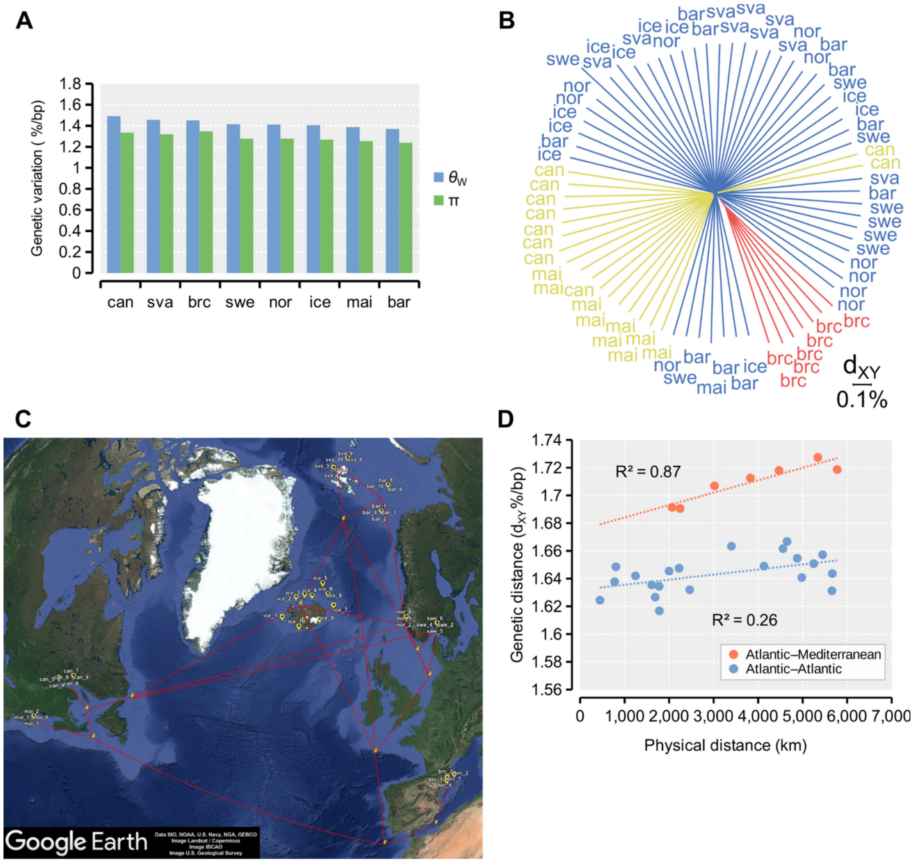


**Fig. S19.**

Levels of genetic variation and genetic distances among populations. (**A**) Genome-wide mean levels of genetic variation in each population estimated using the population mutation rate ($\vartheta_W$) and nucleotide diversity (π) (can=Canada/Gulf of St. Lawrence; mai=USA/Gulf of Maine; ice=Iceland; sva=Svalbard; bar=Barents Sea; nor=Norway/Masfjord; swe=Sweden/Gullmarsfjord; brc=Spain/Catalan Sea). (**B**) Neighbor-joining tree computed from genome-average per-base pairwise genetic distances ($d_{XY}$) between all specimens (labels as in A). (**C**) Physical distance vectors among populations and ocean regions, including mid- or branchpoints. (**D**) Genetic distance compared to physical distance. The average pairwise $d_{XY}$ distances from (A) between all populations over physical distance (North Atlantic Ocean vs. Mediterranean Sea populations: n=7 pairs; North Atlantic vs. North Atlantic: n=21 pairs). Dotted lines indicate linear regressions fitted to the distributions and $R^2$s are the respective goodness of fit for each comparison.

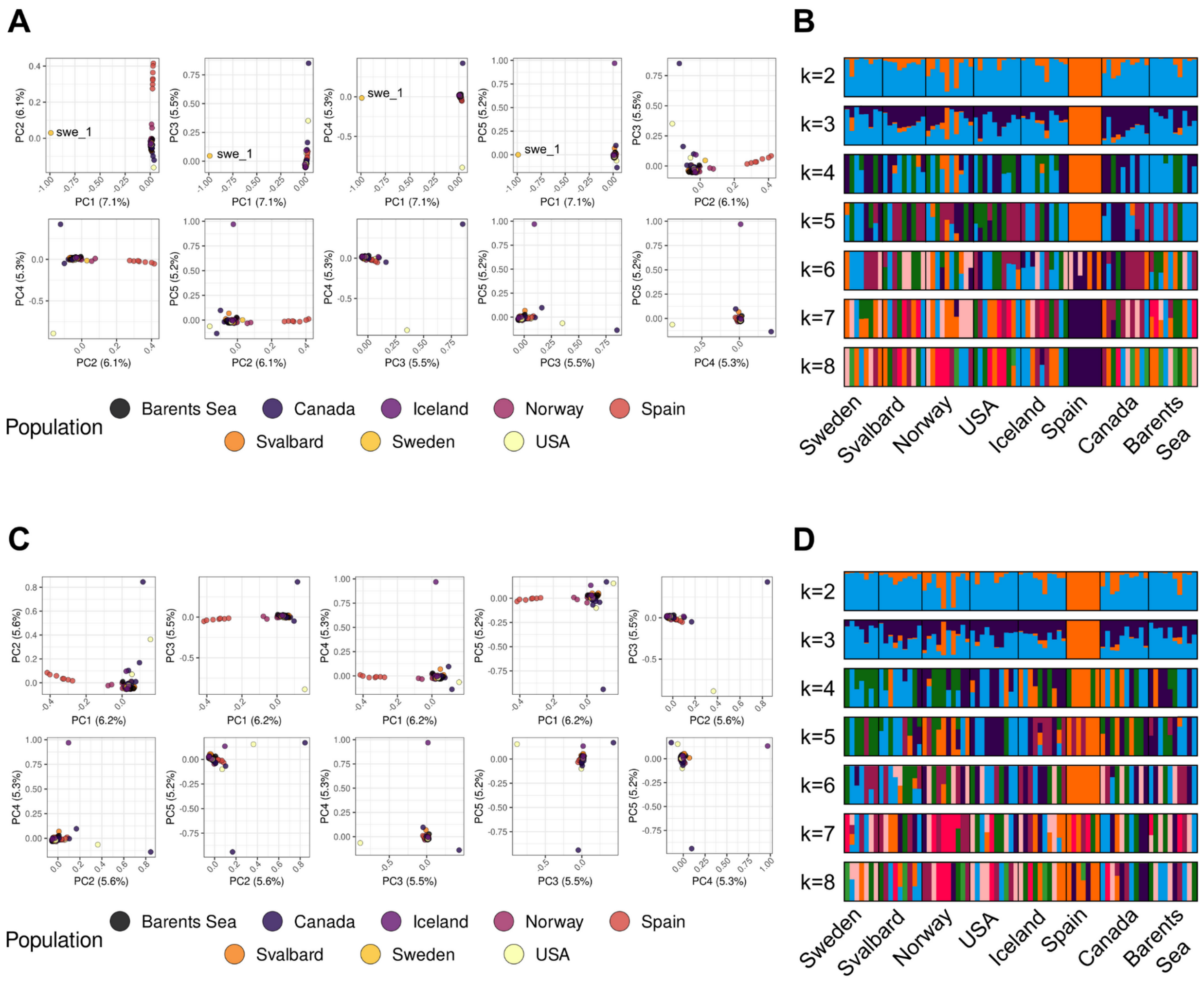


**Fig. S20.**

Signatures of population structure and ancestry components from Principal Component Analysis (PCA) and ADMIXTURE analyses. Top panels (A, B): results including the reference individual, which was sequenced to greater depth than the resequencing individuals. (**A**) The reference individual accounts for most of the variation in PC1, owing to an excess of called and private variants compared to the other individuals. PC2 shows some separation of the Spanish individuals. (**B**) ADMIXTURE results suggest Spain as a separate population for K=2–5, but no clear substructure for remaining populations. Bottom panels (C, D): results excluding the reference individual. (**C**) Separation of Spanish individuals observed in PC1. Higher components show no clear population grouping. (**D**) ADMIXTURE results suggest Spain as a separate population for K=2–3, but no further substructure is observed.

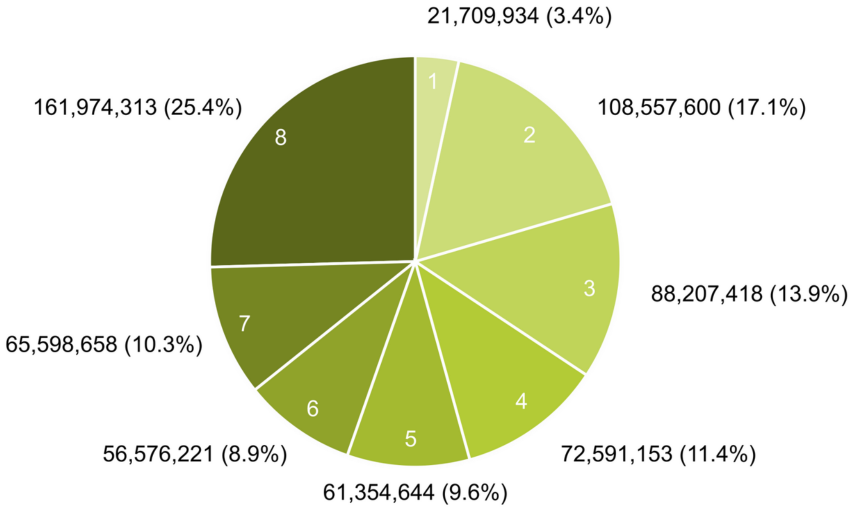


**Fig. S21.**
Allele sharing across the eight studied Northern krill populations. Variants with allele counts of 2 or higher across samples were counted (i.e. excluding singleton alleles). Labels 1–8 indicate in how many populations a particular SNP is polymorphic (as counts or proportions of all SNPs). In total, 60% of the non-singleton variation is shared by five or more populations and only 3% is private.

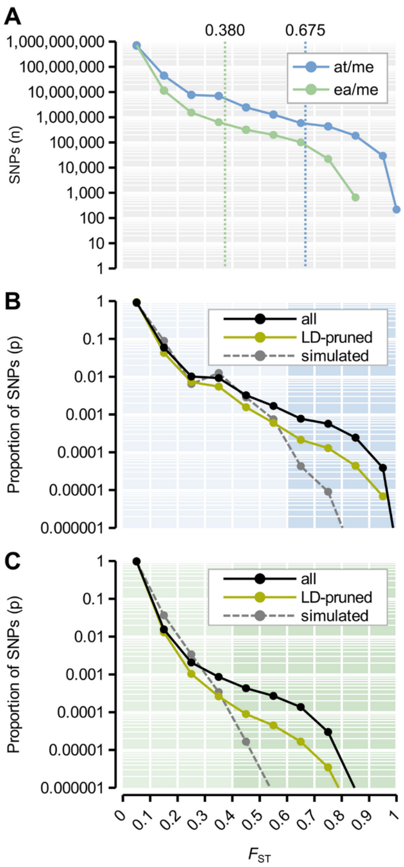


**Fig. S22.**

Genetic divergence between krill sampled across the Atlantic Ocean and Mediterranean Sea. The two contrasts: i) "at/me"=North Atlantic Ocean samples (n=67) vs. the Mediterranean Sea samples (n=7); ii) "ea/we"=North Eastern samples from Iceland, the Barents Sea, Svalbard and Scandinavia (n=47) vs. South Western samples from Canada and the USA (n=20). (**A**) The numbers of SNPs at particular levels of allele frequency divergence in each contrast, as measured using the Fixation index $F_{ST}$ at each SNP. SNPs were binned in $F_{ST}$ intervals of 0.1. $F_{ST}$-scale goes from 0 to 1 as in (C). The top 0.1% percentiles for $F_{ST}$ are indicated for each contrast. (**B**) Observed vs. simulated divergence in the "at/me" contrast. Dotted lines indicate the proportion of SNPs at particular $F_{ST}$ levels for all data ("all"; n=760,436,665 SNPs), a subset of putatively independent SNPs selected to have low levels of linkage disequilibrium ("LD-pruned"; n=7,326,444 SNPs) and neutral SNPs simulated using the same overall divergence as the LD-pruned SNPs ($F_{ST}$=0.0561) under a simple scenario of population subdivision ("simulated; n=7,326,444 SNPs). SNPs were binned as in (A). For the shaded area of divergent SNPs ($F_{ST}$>0.6), the proportions of "all" and "LD-pruned" observed SNPs are 31.1× and 7.5× higher than simulated SNPs ($p_{all}$=0.0016; $p_{LD\text{-}pruned}$=0.000393; $p_{simulated}$=5.21e$^{-05}$), respectively. (**C**) Observed vs. simulated divergence in the "ea/we" contrast using the same approach is in (B), comparing all data (n=738,107,753 SNPs), a subset SNPs with low levels of linkage disequilibrium (n=7,211,757 SNPs) and simulated SNPs (n=7,211,757 SNPs; overall $F_{ST}$=0.0168). SNPs were binned as in (A). For the shaded area of divergent SNPs ($F_{ST}$>0.4), the proportions of "all" and "LD-pruned" observed SNPs are 9.6× and 8.8× higher than simulated SNPs ($p_{all}$=0.00017; $p_{LD\text{-}}$

$_{pruned}$=0.00015; $p_{simulated}$=1.756e$^{-05}$), respectively.

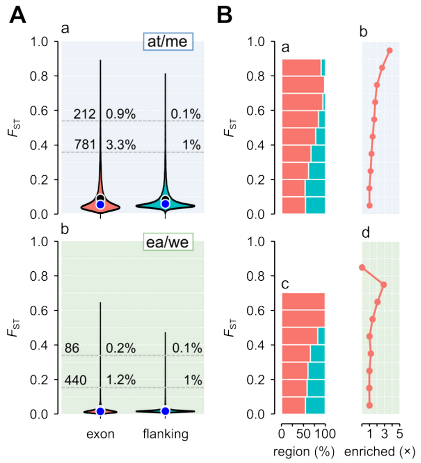


**Fig. S23.**

Genetic divergence in exonic and flanking regions for the subset of ~25,000 functionally annotated genes. First contrast: “at/me”=North Atlantic Ocean samples (n=67) vs the Mediterranean Sea samples (n=7). Second contrast: “ea/we”=North Eastern samples from Iceland, the Barents Sea, Svalbard and Scandinavia (n=47) vs South Western samples from Canada and the USA (n=20). (**A**) The distribution of per-gene $F_{ST}$ values computed across the exons of genes (cds+UTRs) vs. flanking intergenic regions 50–100 kbp away from genes (circles: black=mean; blue=median) (a: n=23,554 vs. n=19,501; b: n=23,542 vs. n=19,060). Gray lines are the 1% and 0.1% percentiles of flanking $F_{ST}$. The number and proportion of genes at that $F_{ST}$-level are indicated. (**B**) First contrast: (a) The relative proportion of genes vs. flanking regions at each level of $F_{ST}$. (b) The enrichment of exonic vs. flanking windows (100 bp) at increasing divergence vs. undifferentiated regions ($F_{ST}$=0.0–0.1). Second contrast: b–c statistics as in a–b.

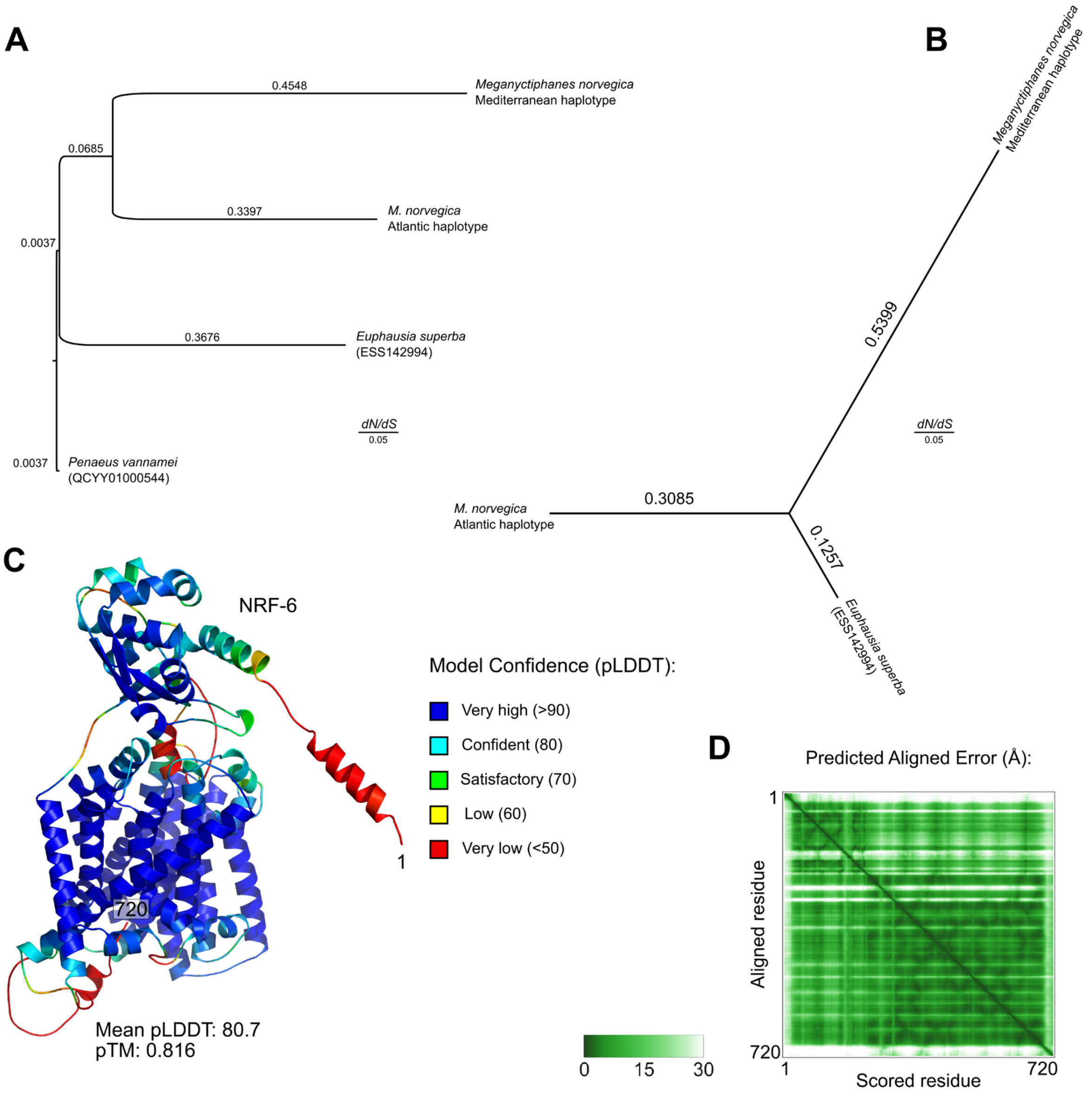


**Fig. S24.**

Molecular evolution and topology of the *nrf-6* gene and encoded protein. (**A**) Maximum likelihood tree scaled by *dN/dS* along each branch (free-ratio model in PAML). The tree includes the Mediterranean and Atlantic Ocean haplotypes of the sequence in the Northern krill, as well as the homologous sequence in the Antarctic krill *Euphausia superba* and the shrimp *Penaeus vannamei*. (**B**). As in (A) but without the shrimp outgroup sequence. (**C**) Per-residue confidence coloring of the top ranked predicted model of NRF-6. The mean predicted local distance difference test (pLDDT) value and pTM score are annotated. (**D**) Residue-residue alignment plot of the predicted NRF-6 model.

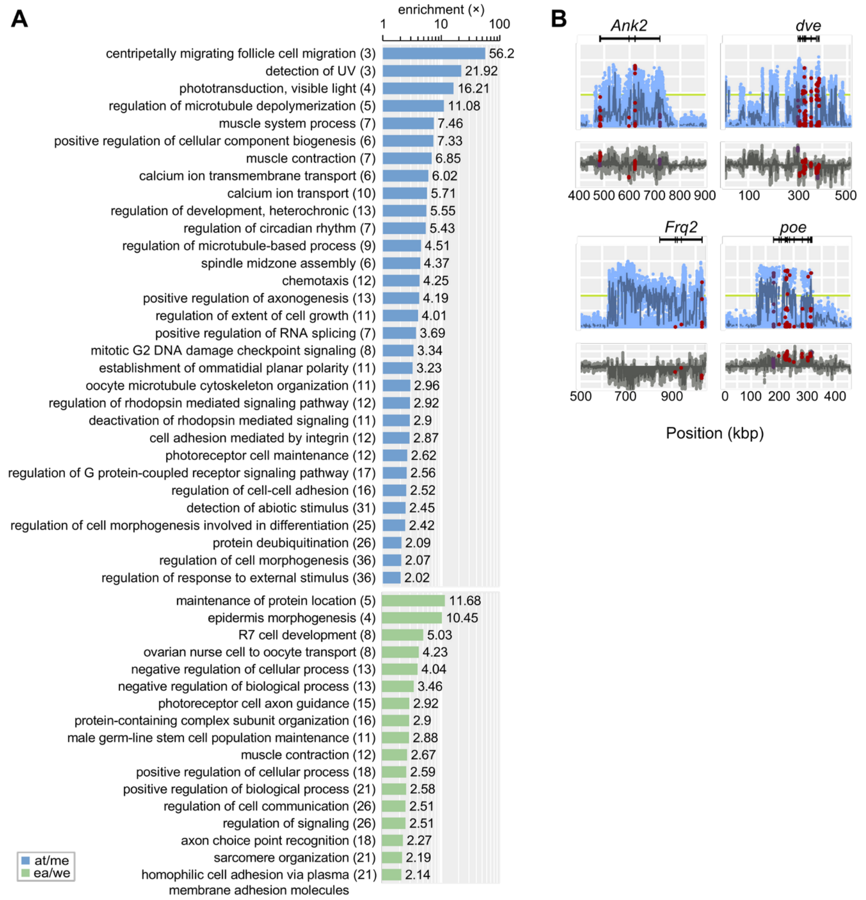


**Fig. S25.**

Gene ontology enrichments (inferred with GOrilla) among divergent genes and examples of loci from the Atlantic contrast after removing small (<3 genes) or redundant ontologies (using Revigo). (**A**) For each of the two contrasts (at/me=Atlantic vs. Mediterranean krill; ea/we=North-East North Atlantic vs South-West North Atlantic krill), the top enriched gene ontology terms from gene-lists ranked by exonic $F_{ST}$. Each list contains non-redundant terms summarized by Revigo after removal of enriched ontologies with few genes (less than three) or low enrichment compared to background (less than 2×). Number of genes associated with each term in parenthesis. (**B**) Examples of divergent genes in the North-East North Atlantic vs South-West North Atlantic contrast associated with regulation and signaling are homologs of fly genes: *Ankyrin 2* (*Ank2*; low identity score towards *Drosophila* homolog: BLAST e-value=0.75; *defective proventriculus* (*dve*); *Frequenin 2* (*Frq2*); *purity of essence* (*poe*).

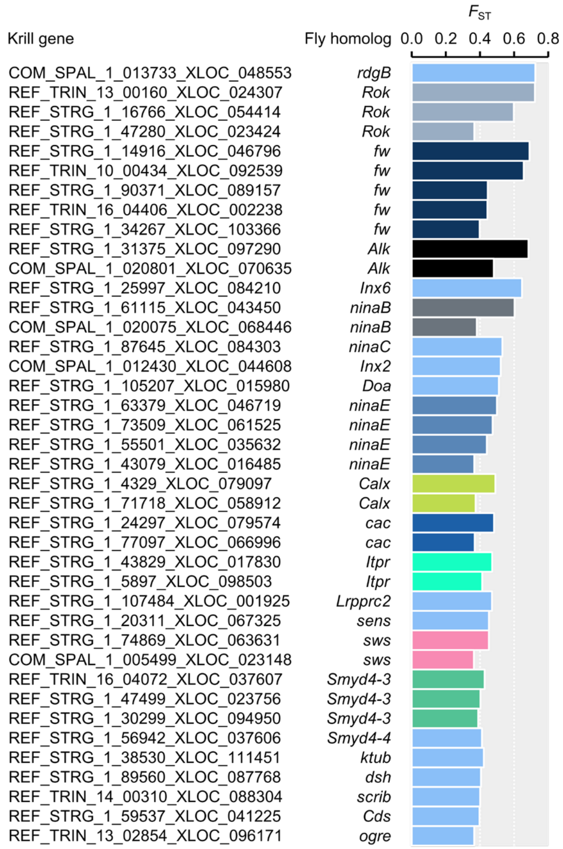


**Fig. S26.**

Visual gene candidates for positive selection in the Atlantic vs. Mediterranean contrast (n=40) and their matching *Drosophila* homologs. Genes associated with significantly enriched eye functions (e.g. phototransduction, compound eye photoreceptor development, establishment of ommatidial planar polarity) were ranked by exon-wide $F_{ST}$ and grouped according to fly homolog. 27 out of 40 genes in this group of genes are putative gene family paralogs. See table S13 for additional homology and ontology details.

**Supplementary tables**

***Table S1.*** See separate file.

Northern krill *M. norvegica* samples sequenced for genome assembly or population genomics.

***Table S2.***

Krill samples sequenced for genome assembly and population genetic analyses.

| Sample | Sequencing | Usage |
|---|---|---|
| K20 ("swe_1" in population dataset)<br>*M. norvegica* reference specimen | Long-read, linked-read and short-read DNA sequencing; RNA/cDNA long-read and short-read sequencing | Genome assembly, scaffolding, polishing and annotation; population genetic analyses |
| K4<br>*M. norvegica* specimen collected together with K20 | Long-read DNA trial sequencing | Preliminary mitochondrial draft assembly |
| Population dataset<br>74 *M. norvegica* specimens | Short-read sequencing | Depth-of-coverage estimation for haplotig identification in genome assembly; population genetic analyses |
| TI787-MN-M (adult male) | RNA short-reads (NCBI SRA: SRR3657321) (*19*) | RNA-based scaffolding and gene annotation |
| TI787-MN-F (adult female) | RNA short-reads (NCBI SRA: SRR3657320) (*19*) | RNA-based scaffolding and gene annotation |

***Table S3.*** See separate file.

Sequence data produced to assemble and annotate the *M. norvegica* genome.

***Table S4.***

Statistics for the *M. norvegica* draft genome assembly and predicted genes.

| Feature | Count |
|---|---|
| Total length w/wo gaps (Gb) | 19.73 / 19.20 |
| Number of genome sequences (n) | 216,568 |
| Scaffold mean (bp) | 91,118 |
| Scaffold N50 / L50 (bp) | 220,856 / 24,545 |
| Scaffold N50 genic scaffolds (bp) | 421,132 |
| Scaffold N50 non-genic scaffolds (bp) | 132,640 |
| Scaffold N90 / L90 (bp) | 38,152 / 111,072 |
| Longest scaffold (bp) | 2,859,246 |
| Contig N50 (bp) | 46,549 |

***Table S5.*** See separate file.

Repeats detected in the genome of *M. norvegica*.

***Table S6.***

Comparative gene models used in SPALN for gene detection.

| Species | Sequences (n) | Source |
|---|---|---|
| *Euphausia superba* (Antarctic krill) | 149,654 | Transcriptome; KrillDB (*63*) |
| *Homarus americanus* (American lobster) | 40,732 | NCBI GCF_018991925.1 (*180*) |
| *Penaeus monodon* (Black tiger shrimp) | 31,641 | https://www.biotec.or.th/pmonodon/index.php (*181*) |
| *Penaeus vannamei* (Whiteleg shrimp) | 33,273 | NCBI GCF_003789085.1 (*182*) |
| *Procambarus virginalis* (Marbled crayfish) | 22,206 | http://marmorkrebs.dkfz.de/ (*183*) |
| *Cherax quadricarinatus* (Australian red claw crayfish) | 19,494 | https://www.frontiersin.org/articles/10.3389/fgene.2020.00201/full#supplementary-material (*184*) |
| *Eriocheir sinensis* (Chinese mitten crab) | 7,549 | http://gigadb.org/dataset/100186 (*185*) |

***Table S7.*** See separate file.

Gene annotations using TransDecoder and EnTAP.

***Table S8.***

Extra gene models used in EnTAP for functional gene annotation.

| Species | Sequences (n) | Source |
|---|---|---|
| *Homarus americanus* (American lobster) | 40,732 | NCBI GCF_018991925.1 (*180*) |
| *Procambarus virginalis* (Marbled crayfish) | 22,206 | http://marmorkrebs.dkfz.de/ (*183*) |
| *Cherax quadricarinatus* (Australian red claw crayfish) | 19,494 | https://www.frontiersin.org/articles/10.3389/fgene.2020.00201/full#supplementary-material (*184*) |
| *Eriocheir sinensis* (Chinese mitten crab) | 7,549 | http://gigadb.org/dataset/100186 (*185*) |
| *Parhyale hawaiensis* (amphipod) | 28,666 | https://research.janelia.org/pavlopoulos/ (*186*) |
| *Daphnia pulex* (water flea) | 30,611 | ENSEMBL https://metazoa.ensembl.org/Daphnia_pulex/Info/Index |
| *Locusta migratoria* (migratory locust) | 29,872 | JAMg (536c4c1) i5k https://i5k.nal.usda.gov/locusta-migratoria |

***Table S9.***

Crustacean species and published resources used for orthology searches in ProteinOrtho and SwiftOrtho.

| Species | Genome size estimate (Gb) in publication | Non-redundant sequences (n) | 1:1 orthologues | Source |
|---|---|---|---|---|
| *Homarus americanus* (American lobster) | 3.06–4.64 | 22,355 | 7,150 | NCBI GCF_018991925.1 (*180*) |
| *Cherax quadricarinatus* (Australian red claw crayfish) | 5 | 19,494 | 4,939 | https://www.frontiersin.org/articles/10.3389/fgene.2020.00201/full#supplementary-material (*184*) |
| *Procambarus virginalis* (Marbled crayfish) | 3.5 | 21,773 | 4,027 | http://marmorkrebs.dkfz.de/ (*183*) |
| *Penaeus monodon* (Black tiger shrimp) | 2.59 | 24,079 | 7,084 | https://www.biotec.or.th/pmonodon/index.php (*181*) |
| *Penaeus vannamei* (Whiteleg shrimp) | 2.45 | 24,974 | 6,914 | NCBI GCF_003789085.1 (*182*) |
| *Hyalella azteca* (amphipod) | 1.05 | 18,608 | 5,835 | NCBI GCF_000764305.1 (*187*) |
| *Parhyale hawaiensis* (amphipod) | 3.6 | 28,666 | 6,093 | https://research.janelia.org/pavlopoulos/ (*186*) |
| *Eurytemora affinis* (copepod) | 0.59–0.69 | 20,716 | 4,551 | NCBI GCF_000591075.1 |

| | | | | |
|---|---|---|---|---|
| | | | | (*188*) |
| *Daphnia magna* | 0.15 | 16,878 | 4,533 | NCBI GCF_020631705.1 |

***Table S10.*** See separate file.

Summary statistics for comparison of the lengths of genes and coding sequences.

***Table S11.*** See separate file.

Analysis of gene family evolution.

***Table S12.*** See separate file.

Datasets used in detecting candidate genes for DNA methylation in the Northern krill.

***Table S13.*** See separate file.

Divergence across genes

***Table S14.*** See separate file.

High $F_{ST}$ variants ($F_{ST}$>0.5) in the coding region of nrf-6

***Table S15.*** See separate file.

Gene enrichments
